# Widespread PRC barrel proteins play critical roles in archaeal cell division

**DOI:** 10.1101/2023.03.28.534520

**Authors:** Shan Zhao, Kira S Makarova, Wenchao Zheng, Yafei Liu, Le Zhan, Qianqian Wan, Han Gong, Mart Krupovic, Joe Lutkenhaus, Xiangdong Chen, Eugene V Koonin, Shishen Du

**Affiliations:** Department of Microbiology, Hubei Key Laboratory of Cell Homeostasis, College of Life Sciences, Wuhan University, Wuhan, Hubei, China; National Center for Biotechnology Information, National Library of Medicine, Bethesda, MD, USA; Institut Pasteur, Unité Biologie Moléculaire du Gène chez les Extrêmophiles, Paris, France; Department of Microbiology, Molecular Genetics and Immunology, University of Kansas Medical Center, Kansas City, Kansas, USA; State Key Laboratory of Virology, College of Life Sciences, Wuhan University, Wuhan, Hubei, China

**Author notes:** These authors contributed equally, and sequence was determined alphabetically (Shan Zhao, Kira S Makarova). <u>To whom correspondence should be addressed:</u> Shishen Du, Department of Microbiology, Hubei Key Laboratory of Cell Homeostasis, College of Life Sciences, Wuhan University, Wuhan, Hubei, China; Eugene Koonin, National Center for Biotechnology Information, National Library of Medicine, Bethesda, MD, USA.

## Abstract

Cell division is fundamental to all cellular life. Most of the archaea employ one of two alternative division machineries, one centered around the prokaryotic tubulin homolog FtsZ and the other around the endosomal sorting complex required for transport (ESCRT). However, neither of these mechanisms has been thoroughly characterized in archaea. Here, we show that three of the four PRC (Photosynthetic Reaction Center) barrel domain proteins of *Haloferax volcanii* (renamed <u>C</u>ell <u>d</u>ivision <u>p</u>roteins B1/2/3 (CdpB1/2/3)), play important roles in division. CdpB1 interacts directly with the FtsZ membrane anchor SepF and is essential for division, whereas deletion of *cdpB2* and *cdpB3* causes a major and a minor division defect, respectively. Orthologs of CdpB proteins are also involved in cell division in other haloarchaea. Phylogenetic analysis shows that PRC barrel proteins are widely distributed among archaea, including the highly conserved CdvA protein of the crenarchaeal ESCRT-based division system. Thus, diverse PRC barrel proteins appear to be central to cell division in most if not all archaea. Further study of these proteins is expected to elucidate the division mechanisms in archaea and their evolution.

## Main

As Francois Jacob famously quipped, “the dream of every cell is to become two cells.” In most bacteria, the FtsZ protein, a tubulin homolog, polymerizes into a dynamic ring-like structure (Z ring) at the division site to recruit other division proteins, forming the division machinery that splits the cell in two^1–7^. Archaea employ multiple division mechanisms, the two most common ones are centered around FtsZ or the endosomal sorting complex required for transport system (ESCRT-III system); some archaea lack both of these systems, suggesting yet to be discovered division mechanisms^8–17^. Recent studies of the Cdv system revealed that it shares structural similarities to the eukaryotic ESCRT machinery and possibly functions in a similar manner^17–23^. Although FtsZ was identified in archaea decades ago^8,15,16^, its function in cell division was only recently studied in detail. Two FtsZ paralogs, FtsZ1 and FtsZ2, are essential for normal cell division in the euryarchaeon *H. volcanii* and perform distinct functions^24^. Two independent studies showed that the highly conserved SepF protein serves as a membrane anchor for FtsZ and is essential for division in both *H. volcanii* and the methanogen *Methanobrevibacter smithii*^25,26^. Phylogenomic analyses show that FtsZ and SepF date back to the Last Universal Cellular Ancestor (LUCA), suggesting that the FtsZ-SepF-based system is the ancestral division apparatus in archaea^13,26^. However, apart from these findings, relatively little is known about the FtsZ-based cell division system in archaea and its regulation because most proteins involved in this process have not yet been identified.

Here, we show that the widespread archaeal PRC (Photosynthetic Reaction Center) barrel proteins, previously predicted to be involved in RNA processing^27^, play critical roles in haloarchaeal cell division. Evolutionary analysis indicates that the PRC barrel domain is widely distributed in archaea, and is present in the CdvA protein of the ESCRT-based division system, suggesting that the PRC barrel domain proteins are widely conserved in cell division.

## Results

### Identification of HVO_1691 as a candidate cell division protein in *H. volcanii*

In search for *H. volcanii* proteins involved in cell division, we used replica plating to screen for proteins that were toxic when overexpressed and discovered an insert containing a portion of the *HVO_1691* gene (Supplementary Figs 1–3). Overexpression of an intact HVO_1691 slightly impaired colony growth but did not cause obvious division and morphological changes (Supplementary Fig. 3). Nonetheless, because expression of this gene has been predicted to be regulated by CdrS^28^, the master regulator of the cell cycle in many archaea^28,29^, we fused HVO_1691 to GFP or mCherry to analyze its subcellular localization and found that it localized to midcell and constriction sites (Supplementary Fig. 4). Moreover, HVO_1691 perfectly co-localized with the known cell division proteins FtsZ1, FtsZ2 and SepF (Fig. 1a). To confirm the involvement of HVO_1691 in division, we determined if its localization depended on FtsZ1, FtsZ2 and SepF using depletion strains in which the expression of these proteins is regulated by the tryptophan-inducible promoter *P_tna_*^24,30^. The depletion of FtsZ1 or FtsZ2 by removing tryptophan from the cultures did not affect the co-localization of HVO_1691 with the other cell division proteins (Fig. 2 and Extended Data Fig. 1). However, in the absence of both FtsZ1 and FtsZ2, HVO_1691 was mostly diffuse in the cytoplasm of the giant misshapen cells with some bright foci (Extended Data Fig. 3a). When SepF was depleted, HVO_1691 also became evenly distributed, whereas FtsZ1 and FtsZ2 still formed ring like structures in the filamentous, misshapen cells (Fig. 1b and Extended Data Fig. 3b). Altogether, these results indicate that HVO_1691 is a component of the FtsZ-based division apparatus. Therefore, we renamed HVO_1691 CdpB1 (<u>C</u>ell <u>d</u>ivision <u>p</u>rotein <u>B1</u>) and its two paralogs CdpB2 and CdpB3 (see below).

**Figure 1.**
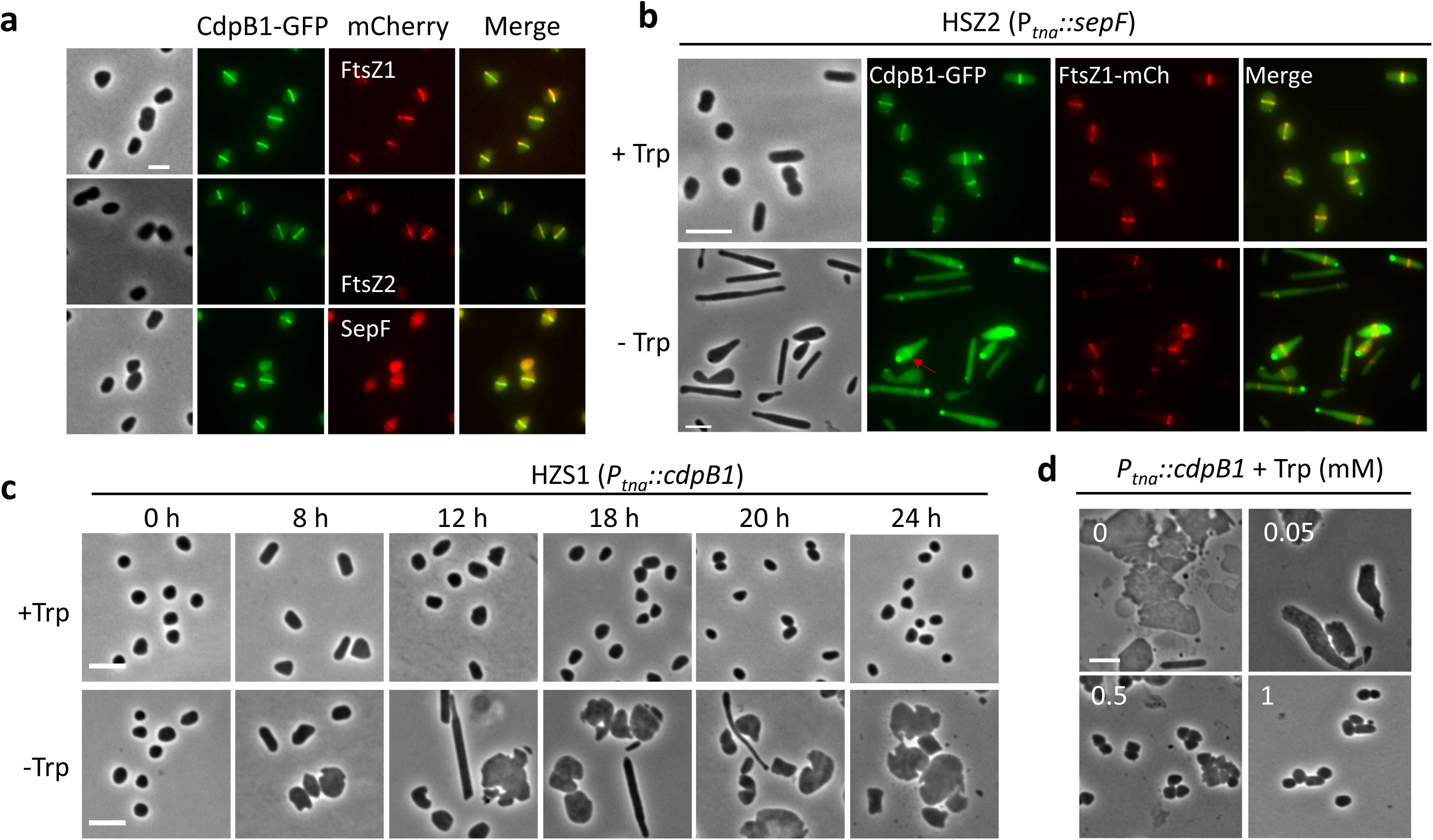
Identification of CdpB1 as a candidate cell division protein in *H. volcanii*. **a**. CdpB1 co-localizes with known cell division proteins. Exponential phase cultures of *H. volcanii* H26 carrying plasmid pZS105 (*P_tna_::cdpB1-gfp*-*ftsZ1-mCherry*), pZS106 (*P_tna_::cdpB1-gfp*-*ftsZ2-mCherry*) or pZS107 (*P_tna_::cdpB1-gfp*-*sepF-mCherry*) were diluted 1:100 in fresh Hv.Cab medium with 0.2 mM Trp, and grown at 45°C to OD_600_ ~ 0.2. 2μL of the cultures was spotted on BSW agarose pads for phase-contrast and fluorescence microscopy. **b**. CdpB1 depends on SepF for localization. An exponential phase culture of strain HZS2 (H98, *P_tna_::sepF*) harboring plasmid pZS239 (*P_phaR_::cdpB1-gfp*-*ftsZ1-mCherry*) was washed with fresh Hv.Cab medium three times to remove tryptophan and then resuspended in fresh Hv.Cab medium. The tryptophan-free culture was then diluted 1:100 in Hv.Cab medium with or without tryptophan and cultured to an OD_600_ and about 0.2. 2 μL of the cultures was spotted on a BSW agarose pad for phase-contrast and fluorescence microscopy. **c**. Depletion of CdpB1 results in severe division and morphological defects. An exponential phase culture of strain HZS1 (H98, *P_tna_::cdpB1*) was treated similarly as panel **1b** to deplete CdpB1. Samples were taken at the indicated times post removal of tryptophan and spotted on a BSW agarose pad for photography. **d**. CdpB1 depleted cells resume normal cell shape and size following restoration of CdpB1. HZS1 grown in Hv.Cab (+50 μg/mL uracil) without tryptophan for 24 hours was diluted 1:100 in fresh medium with different concentrations of tryptophan. 15 hours later, samples were spotted onto a BSW agarose pad for visualization of the cell morphology. Scale bar 5 μm.

**Figure 2.**
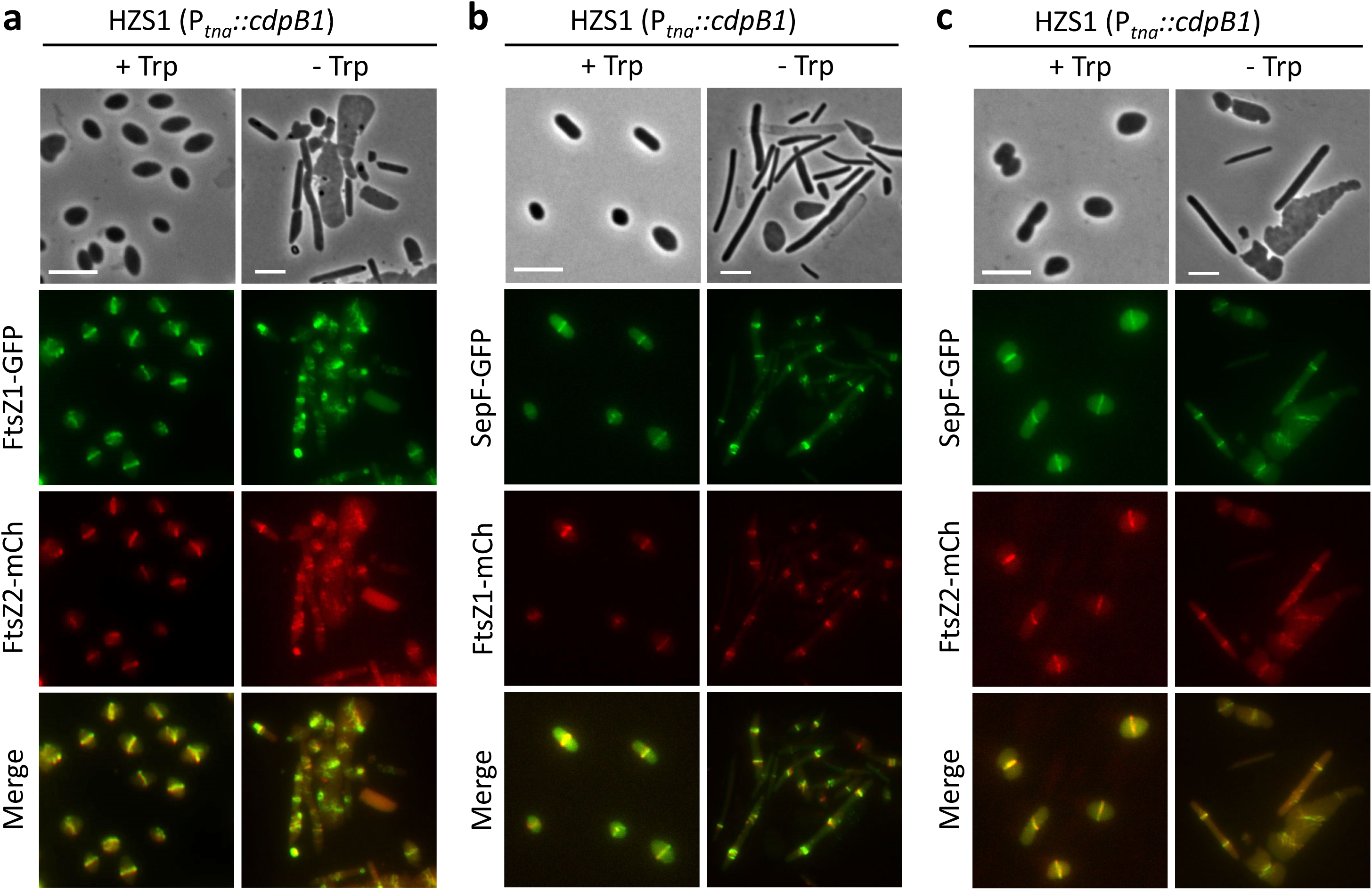
Depletion of CdpB1 does not affect co-localization of FtsZ1, FtsZ2 and SepF. **a.** FtsZ1 co-localizes well with FtsZ2 in the absence of CdpB1. An exponential phase culture of strain HZS1 (H98, *P_tna_::cdpB1*) harboring plasmid pZS289 (*P_phaR_::ftsZ1-gfp*-*ftsZ2-mCherry*) was depleted of CdpB1 as in Fig. 1b and examined by phase-contrast and fluorescence microscopy. **b**. SepF co-localizes well with FtsZ1 in the absence of CdpB1. An exponential phase culture of strain HZS1 harboring plasmid pZS322 (*P_phaR_::sepF-gfp*-*ftsZ1-mCherry*) was treated as in panel **a** and examined by phase-contrast and fluorescence microscopy. **c**. SepF co-localizes well with FtsZ2 in the absence of CdpB1. An exponential phase culture of strain HZS1 harboring plasmid pZS324 (*P_phaR_::sepF-gfp-ftsZ2-mCherry*) was treated as in panel **a** and examined by phase-contrast and fluorescence microscopy. Scale bar 5 μm.

### CdpB1 is important for cell division and cell shape in *H. volcanii*

To test whether CdpB1 was essential for cell division in *H. volcanii*, we attempted to generate a *cdpB1* deletion mutant by the standard pop-in/pop-out approach^30^. However, we failed to obtain a *cdpB1* deletion strain after numerous attempts. Therefore, we constructed a CdpB1 depletion strain by replacing its native promoter with the *P_tna_* promoter^31^ so that the expression level of CdpB1 was regulated by tryptophan (Supplementary Fig. 5). The depletion strain grew well in the presence of tryptophan but cells gradually became enlarged and misshapen in the absence of tryptophan (Fig. 1c and Supplementary Fig. 6), indicating that CdpB1 plays an important role in division and cell morphology. The CdpB1 depleted cells could still grow on plates or in liquid medium without tryptophan (Supplementary Fig. 6), presumably, due to leaky expression of CdpB1 from the *P_tna_* promoter. The CdpB1 depleted cells resumed division and normal shape in a tryptophan dependent manner. At 1 mM tryptophan, the size and morphology of the cells were comparable to those of wild type cells (Fig. 1d). Thus, CdpB1 is likely essential, its level is important for normal cell division and cell shape in *H. volcanii* and the effects of its depletion are entirely reversible.

### CdpB1 is not required for the localization of FtsZs and SepF to the division site

To explore the function of CdpB1 in haloarchaeal cell division, we checked if its depletion affected the co-localization of FtsZ1, FtsZ2 and SepF. FtsZ1, FtsZ2 and SepF co-localized even in the filamentous and enlarged misshapen cells that formed upon CdpB1 depletion (Fig. 2). In the filamentous cells, FtsZ1, FtsZ2 and SepF formed both clustered spiral ring-like structures as well as normal Z rings, whereas in the giant misshapen cells, FtsZ1, FtsZ2 and SepF mostly formed patches of filamentous structures (Fig. 2). These observations indicate that the abnormal localization of FtsZs and SepF in CdpB1 depleted cells reflect the division block and altered cell morphology rather than a direct effect of CdpB1 on Z ring formation.

### Depletion of CdpB1 mimics depletion of FtsZ2 or SepF

The localization of FtsZs and SepF in the CdpB1 depleted cells resembled that observed when SepF or FtsZ2 was depleted^24,25^, suggesting that CdpB1 is involved in the same division step(s) as FtsZ2 and SepF. To test this hypothesis, we compared the cell morphology and FtsZ1 localization in cells depleted of CdpB1, FtsZ2 or SepF. Cells of all three depletion strains began to elongate and enlarge 6 hours post tryptophan removal, with FtsZ1 localizing in disorganized spiral ring-like structures (Extended Data Fig. 4). Around 9 hours post depletion, FtsZ1 formed one loose spiral Z ring and abnormal structures in the filamentous and misshapen cells (Extended Data Fig. 4). By 12 hours post depletion, multiple regularly spaced spiral FtsZ1 rings or abnormal FtsZ1 structures were observed in the cell filaments or giant cells. These abnormal phenotypes exacerbated with time so that, by 21 hours post depletion, many giant abnormal cells along with cell debris were observed (Extended Data Fig. 4). Thus, depletion of CdpB1, similar to the depletion of FtsZ2 and SepF, results in a gradual loss of normal FtsZ1 localization, severe impairment of cell division and, as a result, abnormal cell morphology.

### CdpB1 interacts with SepF *in vivo* and *in vitro*

To test if CdpB1 interacted with SepF and FtsZ1 or FtsZ2 *in vivo*, we employed the Split-FP (Fluorescent Protein) assay^32,33^. In this assay, the superfolder GFP is split into three parts: the complementary detector GFP1-9, and two twenty amino acids long tags, GFP10 and GFP11. Protein partners of interest are fused to GFP10 and GFP11, respectively. If the putative protein partners interact, GFP10 and GFP11 are brought in proximity to self-associate with GFP1-9, reconstituting a functional GFP. In our case, because the protein pairs were involved in cell division, we would not only detect fluorescence but also localization at the division site. Indeed, we detected strong fluorescence and observed fluorescent rings in cells when CdpB1 and SepF were fused to the GFP tags but not in cells carrying GFP tags only or when only one of the two proteins was tagged, indicating that CdpB1 and SepF strongly interact *in vivo* (Fig. 3a-b and Extended Data Fig. 5). Cells expressing CdpB1 and FtsZ2 or FtsZ1 fused to the GFP tags also displayed fluorescent rings at midcell (Extended Data Fig. 5b), however, given that neither FtsZ2 nor FtsZ1 were individually required for CdpB1 localization, this signal was likely due to the interaction between CdpB1 and SepF which brought CdpB1 and FtsZ2/FtsZ1 close enough to generate the fluorescence.

**Figure 3.**
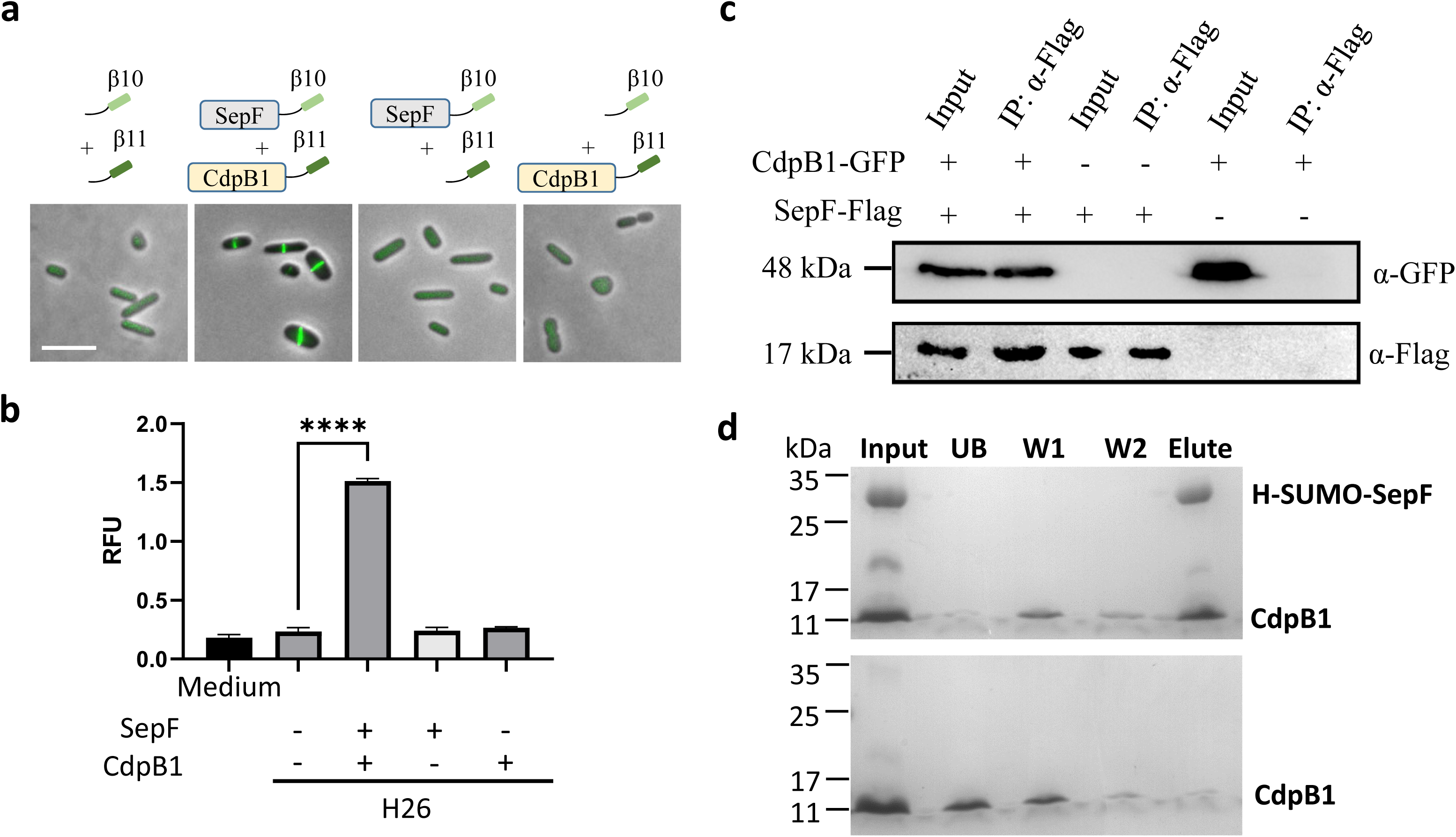
CdpB1 directly interacts with SepF *in vivo* and *in vitro*. **a.** Split-FP assay shows that CdpB1 interacts with SepF *in vivo*. An exponential culture of strain H26 (DS70, *ΔpyrE2*) harboring the split-FP plasmid was cultivated in Hv.Cab at 45°C with 0.2 mM Trp to an OD_600_ of about 0.2 followed by a shift to 37°C 3h. 2μL of the culture was spotted on a BSW agarose pad for phase-contrast and fluorescence microscopy. Scale bar 5 μm. **b.** Quantitation of the interaction signal between CdpB1 and SepF. An exponential culture of strain H26 (DS70, *ΔpyrE2*) harboring the split-FP plasmid was cultivated in Hv.Cab at 45°C with 0.2 mM Trp to an OD_600_ about 1.0 followed by 30°C overnight. 1 mL of the culture was washed and resuspended to OD_600_ about 1.0 with 18% BSW (includes calcium). Samples of 200 μL were analyzed with three technical replicates in a 96-well plate and evaluated using the Varioskan LUX multifunctional microplate detection system. The p-values were calculated using the Student t-test. **c.** Co-IP shows that CdpB1 interacts with SepF *in vivo*. Overnight cultures of *H. volcanii* carrying the indicated expression plasmid were diluted 1:100 in 40 mL fresh Hv.YPC medium, and cultivated at 45°C to OD_600_ about 1.0. Cells were collected and lysed by sonication and then centrifuged at 12,000 rpm for 5 min at 4°C to remove cell debris. Supernatants were incubated with anti-bodies coated magnetic beads. Immunocomplexes were eluted with boiling SDS-PAGE loading buffer and then analyzed by immunoblot. **d**. CdpB1 interacts with SepF *in vitro*. H-SUMO-SepF and CdpB1 were incubated and treated according to the pull-down assay described in Methods. All fractions were collected during the procedure and analyzed by SDS-PAGE.

To validate the interaction between CdpB1 and SepF *in vivo*, we performed co-immunoprecipitation (Co-IP) experiments with cells expressing a GFP-tagged CdpB1 and a Flag-tagged SepF. CdpB1-GFP was detected in immunocomplexes isolated with anti-Flag antibodies in cells expressing both SepF-Flag and CdpB1-GFP but not in cells expressing only one of the fusion proteins (Fig. 3c), and vice versa (Supplementary Fig. 7). These results indicate that CdpB1 interacts with SepF under physiological conditions.

To further test the interaction between CdpB1 and SepF *in vitro*, we purified these proteins in *Escherichia coli* using the SUMO-tag protein purification system^34^. The SUMO and His tags were removed from SUMO-CdpB1 but not from SepF (SUMO-SepF) so that we could test the interaction between the two proteins using a pull-down assay. When CdpB1 was incubated with SUMO-SepF, both proteins were retained on Ni-NTA beads and most of the CdpB1 was found in the eluate (Fig. 3d). In contrast, when CdpB1 was incubated with the Ni-NTA beads alone, most of the protein was found in the unbound fraction and none was detected in the eluate (Fig. 3d), indicating that retention of CdpB1 on the beads was due to specific interaction with SUMO-SepF. Taken together, these results demonstrate that CdpB1 directly interacts with SepF to participate in cell division.

### CdpB1 is a member of the conserved PRC barrel protein family widely present in archaea

CdpB1 is a small protein of 97 amino acids that belongs to the vast PRC barrel protein family widely present in archaea, bacteria and plants (Pfam ID: PF05239)^27^. Although most of the PRC barrel proteins remain poorly characterized, four distinct functions have been described: 1) H subunit of photosynthetic reaction center (PRC)^35^; 2) bacterial RimM protein involved in 30S ribosome maturation^36^; 3) sporulation proteins YlmC/YmxH in Firmicutes^37^ and 4) CdvA component of the crenarchaeal and thaumarchaeal ESCRT-based cell division system^18^. Prompted by the role of CdpB1 in haloarchaeal cell division, we reexamined the PRC barrel protein family, focusing on archaeal proteins previously found to be monophyletic in the phylogeny of the PRC barrel family and hypothesized to be involved in translation by analogy with RimM^27^ (see Methods). Initially, five arCOGs (2155, 2157, 2158, 5740, 8023) were assigned to the PRC barrel family, but by extensive sequence comparison performed with sensitive methods (see Methods), we identified two additional arCOGs (8931 and 10234) and expanded arCOG04054 (CdvA) by identifying CdvA orthologs in several *Thermoproteales* genomes (Supplementary Table 1). Altogether, at least one PRC barrel domain containing protein was identified in 509 of the 524 searched archaeal genomes (Supplementary Table 1). Phylogenetic analysis of the PRC barrel family revealed three major branches (Fig. 4a, Supplementary File 1), each of which included representatives of several archaeal lineages from two major phyla (Euryarchaeota and Asgardarchaeota) and two superphyla (TACK and DPANN), suggesting that the last archaeal common ancestor encoded three distinct PRC barrel proteins.

**Figure 4.**
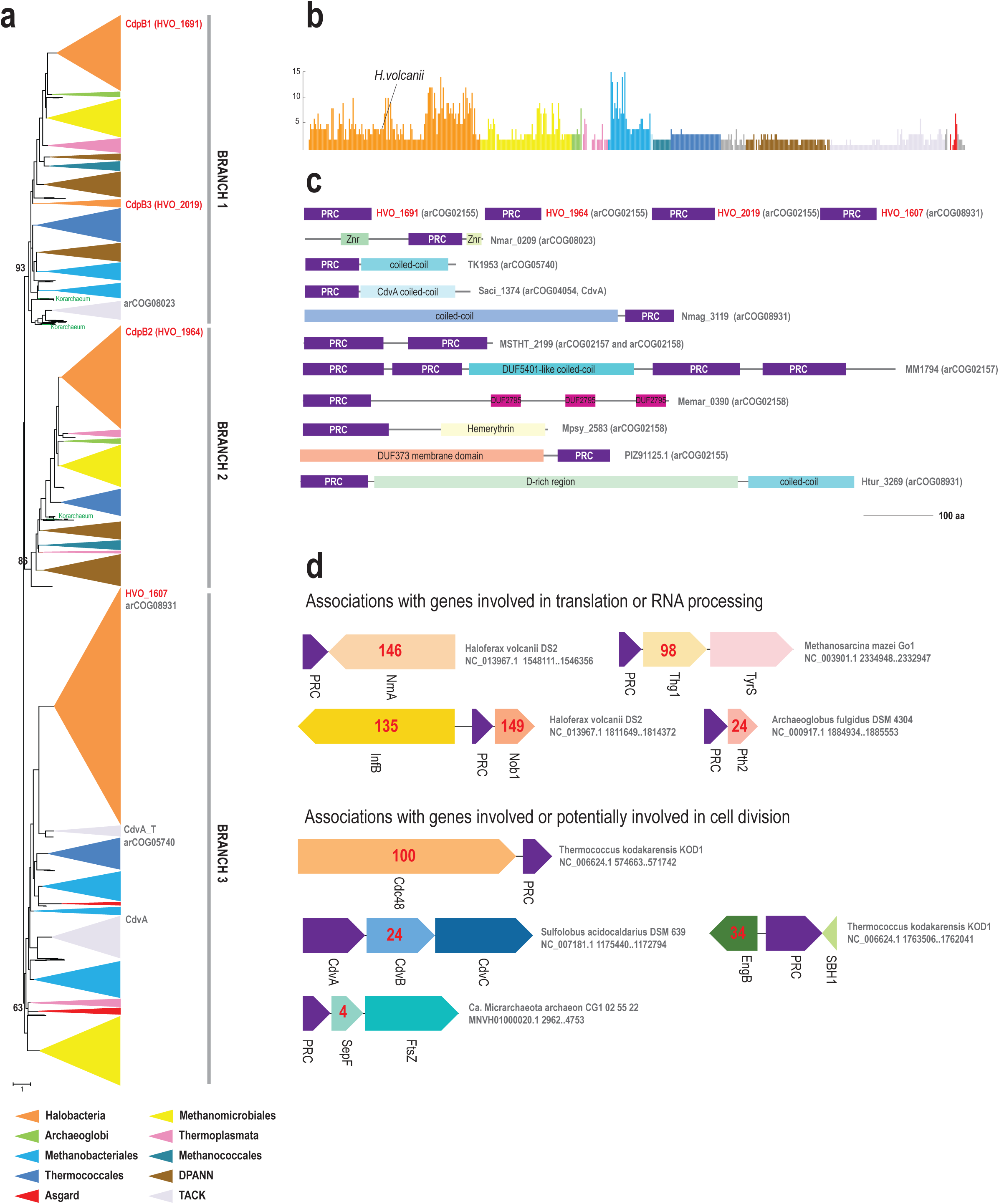
Phylogeny, comparative genomics, domain architectures and gene neighborhood analysis of PRC barrel family in archaea. **a.** Schematic representation of phylogenetic tree of the PRC barrel domain family. The phylogenetic tree was built using the FastTree approximate maximum likelihood method as described under Methods. Branches corresponding to major archaeal phyla are collapsed and colored according to the color key below the dendrogram. Three bootstrap values supporting major branches are shown. Korarchaeal sequences are highlighted in green. Several distinct arCOGs corresponding to collapsed branches and *H. volcanii* PRC barrel proteins identifiers, belonging to the respective branches, are indicated on the right. The complete tree in Newick format is available in Supplementary File 1. **b**. Numbers of PRC barrel domain proteins in 524 archaeal genomes from the arCOG database. The plot shows the number of distinct PRC barrel domains in all 524 genomes from arCOG database grouped according to their taxonomy and colored using the same color code as in A, except for the gray color, indicating unclassified genomes. **c**. Selected most frequent domain architectures of PRC barrel proteins. The proteins are shown to scale as indicated. Distinct domains are shown by colored rectangles. The respective protein identifier and arCOG number are indicated on the right. **d**. Selected neighborhoods of PRC barrel genes. Genes are shown as arrows. Protein name is indicated below each arrow. The numbers inside the arrows indicate the counts of the occurrences of the respective gene in 1675 PRC barrel gene neighborhoods (Supplementary Table 2). Genome description, nucleotide accession and coordinates of the neighborhood are indicated on the right. Abbreviations: Nob1, endonuclease Nob1; Cdc48, ATPase of the AAA+ class; NrnA, nanoRNase/pAp phosphatase; Thg1, tRNA-His guanylyltransferase; InfB, translation initiation factor 2; EngB, cell division controlling GTPase; Pth2, Peptidyl-tRNA hydrolase; SepF, cell division protein; FtsZ, cell division GTPase; CdvB, cell division protein ESCRT III family; CdvC, Vps4 family ATPase, component of ESCRT cell division system; CdvA, component of ESCRT cell division system; TyrS, Tyrosyl-tRNA synthetase; SBH1, preprotein translocase subunit Sec61beta.

The number of genes encoding PRC barrel proteins varies greatly among archaea even within the same lineage, such as *Halobacteria* or *Methanomicrobia*, with up to 15 genes in several *Methanobacteriales* species (Fig. 4b). *H. volcanii* encodes 4 PRC barrel proteins: CdpB1 (HVO_1691) and HVO_2019 in Branch 1, HVO_1964 in Branch 2 and a more distant paralog HVO_1607 (arCOG08931) in Branch 3 (Fig. 4a and 4c). Although the majority of the PRC barrel proteins contain a single domain, proteins with duplicated PRC barrel domains or a PRC barrel fused to other domains are also widespread including CdvA^9,18^ in which the PRC barrel is fused to a coiled-coil domain (Fig. 4c). The majority of PRC barrel proteins are encoded by standalone genes, but many are embedded in conserved neighborhoods or putative operons (Fig. 4d, Supplementary Table 2). Notably, PRC barrel proteins are often found in the vicinity of genes encoding translation systems components or RNA processing enzymes. Specifically, *cdpB1* is encoded next to *nrnA*, a 5’-3’ exonuclease involved in processing of short RNA substrates^38^, and HVO_1964 is encoded divergently to *infB*, translation initiation factor 2. These two genes are located in conserved neighborhoods in haloarchaea (Supplementary Table 2). In *Methanomicrobia*, the most highly conserved neighborhood includes *thg1*, a tRNA-His guanylyltransferase^39^ which might be co-expressed with the PRC barrel gene (Fig. 4d). In *Archaeoglobi* a PRC barrel gene is encoded in a putative operon with peptidyl-tRNA hydrolase *pth2* (Supplementary Table 2). Several other, relatively frequent neighborhoods include genes that are potentially involved in cell division control, in particular, Cdc48 family ATPase and *engB* GTPase^40,41^, both of which are implicated in cell cycle regulation but have not been characterized in archaea (Fig. 4d, Supplementary Table 2). CdvA is encoded in the *cdv* cluster along with other genes involved in cell division (Fig. 4d). Surprisingly, we found only one genome, *Micrarchaeota* archaeon from the DPANN superphylum, where a PRC barrel protein is encoded in the vicinity of genes encoding components of the FtsZ-based division machinery, in a putative operon with *sepF* and *ftsZ* (Fig. 4d, Supplementary Table 2). Thus, PRC barrel proteins might be involved in other house-keeping functions, in addition to cell division. Overall, our analysis shows that the PRC family is actively evolving in archaea and some subfamilies are likely to be sub- and neofunctionalized to participate in diverse cellular processes.

### CdpB1 paralogs are also involved in cell division in *H. volcanii*

The three other PRC barrel proteins in *H. volcanii* (HVO_1607, HVO_1964 and HVO_2019) are relatively distant paralogs of CdpB1 (25-35% sequence identity), but the predicted structures of their respective PRC barrels are nearly identical (Extended Fig. 6). To test if these proteins were also involved in cell division, we fused them to GFP and examined their localization. HVO_1607 showed a diffusive localization pattern, but HVO_1964 and HVO_2019 formed midcell ring-like structures (Fig. 5a), suggesting that these two proteins were involved in cell division. Thus, we renamed them CdpB2 and CdpB3, respectively. However, unlike *cdpB1*, *cdpB2* and *cdpB3* could be knocked out. Deletion of *cdpB2* caused severe division and shape defects in the semi-defined Hv-Cab medium where Casamino acids were used as the carbon and energy source, however, in the complex medium Hv-YPC, the division and shape defects of *ΔcdpB2* cells were less pronounced (Fig. 5b). By contrast, deletion of CdpB3 only resulted in a minor division defect in either media (Fig. 5b). We found that the CdpB proteins were recruited to the Z ring in a sequential manner: CdpB1 was required for the midcell localization of CdpB2 and CdpB3 whereas CdpB2 was necessary for the localization of CdpB3 but not CdpB1, and absence of CdpB3 did not affect the localization of CdpB1 or CdpB2 (Extended Data Fig. 7). Overexpression of CdpB2 or CdpB3 alone or in combination did not suppress the division and morphological defects of the CdpB1 depleted cells (Fig. 5c), whereas overexpression of CdpB1 but not CdpB3 largely suppressed the defects of the *ΔcdpB2* cells (Fig. 5d). These results indicate that, although all three CdpB proteins participate in cell division, they have distinct functions, with CdpB1 playing a dominant role.

**Figure 5.**
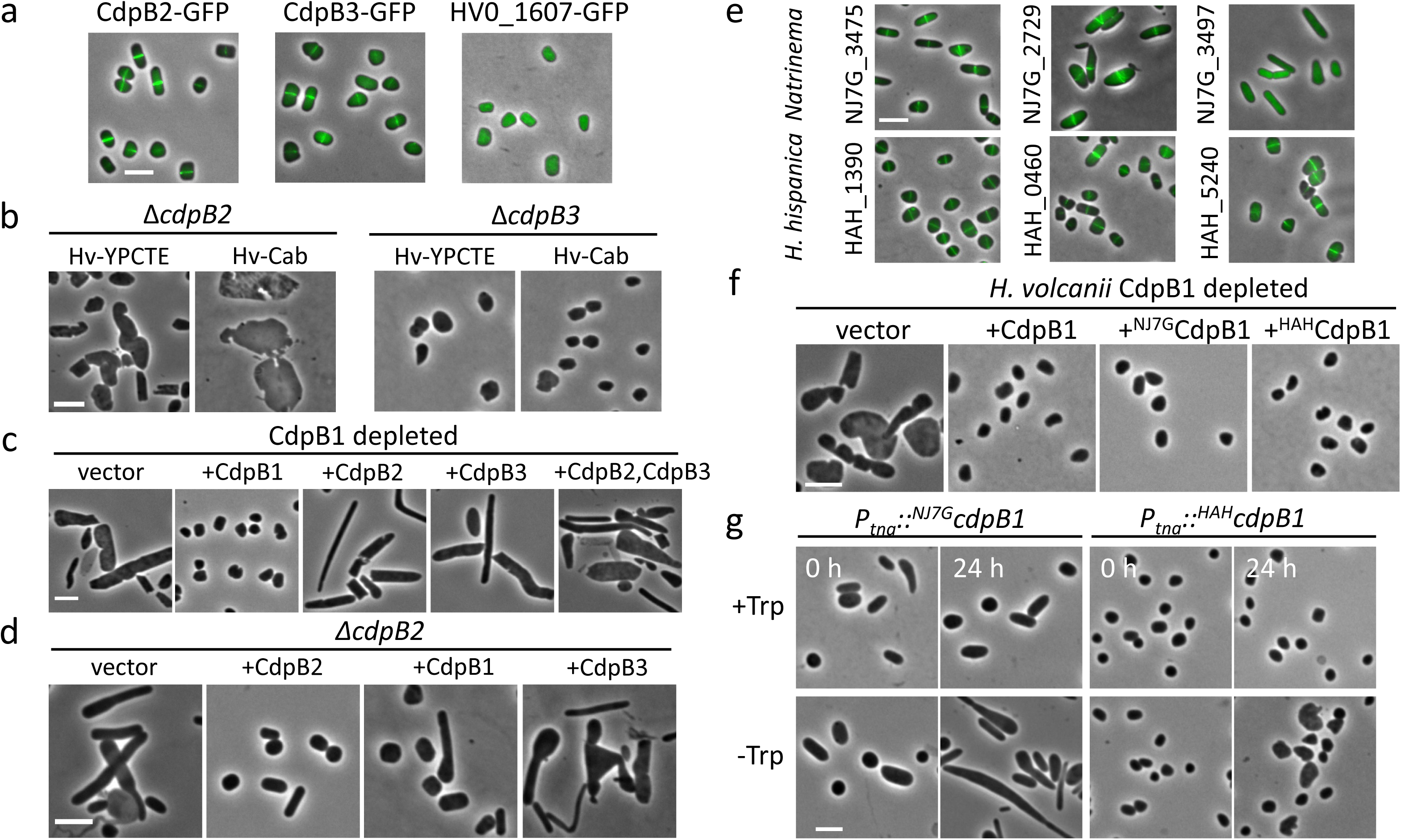
CdpB proteins are involved in cell division in diverse haloarchaea. **a.** CdpB1 paralogs localize as midcell ring-like structures. Exponential phase cultures of *H. volcanii* H26 carrying plasmid pZS336 (*P_tna_::cdpB2-gfp*) or pZS337 (*P_tna_::cdpB3-gfp*) were diluted 1:100 in fresh Hv.Cab medium with 0.2 mM Trp, and grown at 45°C to OD_600_ ~ 0.2. 2 μL of the cultures was spotted on BSW agarose pads for phase-contrast and fluorescence microscopy. **b.** *cdpB2* and *cdpB3* deletion strains display severe and minor cell division and shape defects, respectively. The single colony of HZS5 (Δ*cdpB2*) and HZS6 (Δ*cdpB3*) were inoculated into fresh Hv-YPCTE or Hv.Cab medium with 50 μg/mL uracil and grown at 45°C to OD_600_ about 0.2. 2 μL of the cultures was spotted on a BSW agarose pad for photography. **c**. Overexpression of CdpB2 and CdpB3 alone or in combination cannot suppress the division and shape defect of CdpB1 depleted cells. Exponential phase cultures of HZS1 (H98, *P_tna_::cdpB1*) carrying plasmid pZS236 (*P_native_::cdpB1*), pZS339 (*P_native_::cdpB2*), pZS338 (*P_native_::cdpB3*) or pZS390 (*P_native_::cdpB2*-*P_native_::cdpB3*) were treated similarly as Figure 1b to deplete CdpB1. Samples were spotted onto a BSW agarose pad for visualization of the cell morphology. **d**. Overexpression of CdpB1 but not CdpB3 largely suppresses the division and shape defect of Δ*cdpB2* cells. Exponential phase cultures of HZS5 carrying plasmid pZS236 (*P_native_::cdpB1*), pZS339 (*P_native_::cdpB2*) or pZS338 (*P_native_::cdpB3*) were diluted 1:100 in fresh Hv-Cab medium, and grown at 45°C to OD_600_ about 0.2 to examine their impact on cell division and morphology. **e.** PRC barrel proteins localize to the midcell as a ring in *Natrinema sp*. J7 and *H. hispanica* cells, respectively. Exponential phase culture of strain *Natrinema sp*. CJ7-F harboring plasmid pZS217 (*P_tna_:: NJ7G_3475-gfp*), pZS384 (*P_tna_::NJ7G_2729-gfp*), pZS385 (*P_tna_::NJ7G_3497-gfp*) or *H. hispanica* DF60 carrying plasmid pZS306 (*P_tna_::HAH_1390-gfp*), pZS388 (*P_tna_::HAH_0460-gfp*), pZS389 (*P_tna_::HAH_5240-gfp*) were treated as in Figure 5a for phase contrast and fluorescence microscopy. Hv-Cab medium was used for *Natrinema sp. J7* and AS-168-M medium was used for *H. hispanica* DF60 strain. **f.** ^NJ7G^CdpB1 and ^HAH^CdpB complement CdpB1 depleted *H. volcanii* cells. Exponential phase cultures of HZS1 carrying plasmid pZS236 (*P_native_::cdpB1*), pZS234 (*P_native_::^NJ7G^cdpB1*) or pZS276 (*P_native_::^HAH^cdpB1*) were treated similarly as Figure 1b to check their impact on cell division and morphology. **g.** Depletion of CdpB1 homologs causes severe and modest division defects in *Natrinema sp*. J7 and *H. hispanica* cells, respectively. Strain HZS4 (CJ7-F, *P_tna_*::*^NJ7G^cdpB1*) or HZS3 (DF60, *P_tna_*::*^HAH^cdpB1*) was treated as in Fig. 1c, but HZS4 was grown in AS-168-M medium. Scale bar 5 μm.

### CdpB homologs are critical for cell division in diverse haloarchaea

Similar to *H. volcanii*, many haloarchaea encode multiple PRC barrel proteins (Fig. 4a and Extended Data Fig. 6). To determine if these proteins were also involved in division, we tested if they localized to midcell as ring-like structures in *Natrinema sp. J7* and *Haloarcula hispanica*. As shown in Fig. 5e, all the tested PRC barrel proteins from *Natrinema sp. J7* and *H. hispanica* formed ring-like structures in the respective species except for NJ7G_3497, indicating that most of the PRC proteins participate in cell division in these two species. We focused on the CdpB1 orthologs, NJ7G_3475 (renamed ^NJ7G^CdpB1) and HAH_1390 (renamed ^HAH^CdpB1), and tested if they could complement the CdpB1 depleted *H. volcanii* cells. As shown in Fig. 5f, expression of either ^NJ7G^CdpB1 or ^HAH^CdpB1 in CdpB1 depleted *H. volcanii* cells restored normal cell division and morphology, indicating that CdpB1 homologs from other haloarchaea likely function in a similar manner. To further confirm their role in cell division, we tried to generate their deletion strains in *Natrinema sp. J7* and *H. hispanica*, but these attempts failed consistent with the results for CdpB1 in *H. volcanii*. Thus, we again used the *P_tna_* promoter to replace the native promoter regions, in order to obtain depletion strains (Supplementary Fig. 8). Similar to the depletion of CdpB1 in *H. volcanii*, depletion of ^NJ7G^CdpB1 in *Natrinema sp. J7* cells resulted in a severe cell division defect (Fig. 5e). By contrast, depletion of ^HAH^CdpB1 in *H. hispanica* cells only caused a modest cell division and morphological defect. Also, both ^NJ7G^CdpB1 and ^HAH^CdpB1 depletion strains could grow on plates without tryptophan (Supplementary Fig. 6), presumably, due to the leaky expression from the *P_tna_* promoter. Nonetheless, given that ^NJ7G^CdpB1 and ^HAH^CdpB1 could complement the CdpB1 depleted *H. volcanii* cells and these genes could not be deleted from the chromosomes of the respective species, CdpB1-like proteins likely play a critical role in cell division in diverse haloarchaea.

## Discussion

Recent work on archaeal cell division revealed interesting mechanisms for controlling this process and cell organization, and provided insights into the diversifying evolution of the division machinery of LUCA^13,14,21, 24–26^. However, compared to the extensive knowledge on bacterial and eukaryotic cell division, relatively little is known about this process in archaea. In this work, we find that the PRC barrel proteins of *H. volcanii* play important, but distinct roles in cell division. CdpB1 is essential for division, whereas its two paralogs, CdpB2 and CdpB3, are not. Our analysis indicates that CdpB1 is recruited to the Z ring by SepF, via a direct interaction between the two proteins, and it functions as the recruiter of CdpB2 which in turn recruits CdpB3 (Fig. 6). Moreover, the function of CdpB proteins are highly conserved in halophiles. In addition, PRC barrel proteins are widely distributed in archaea, including the DPANN superphylum which consists of symbiotic archaea with small genomes. Notably, the highly conserved CdvA protein of the ESCRT-based division system also contains a PRC barrel domain (Fig. 4c), suggesting that the PRC barrel domain is important for cell division in both the FtsZ-based and ESCRT-based division systems. Similar findings by a completely different approach are reported in a complementary study from the Albers lab.

**Figure 6.**
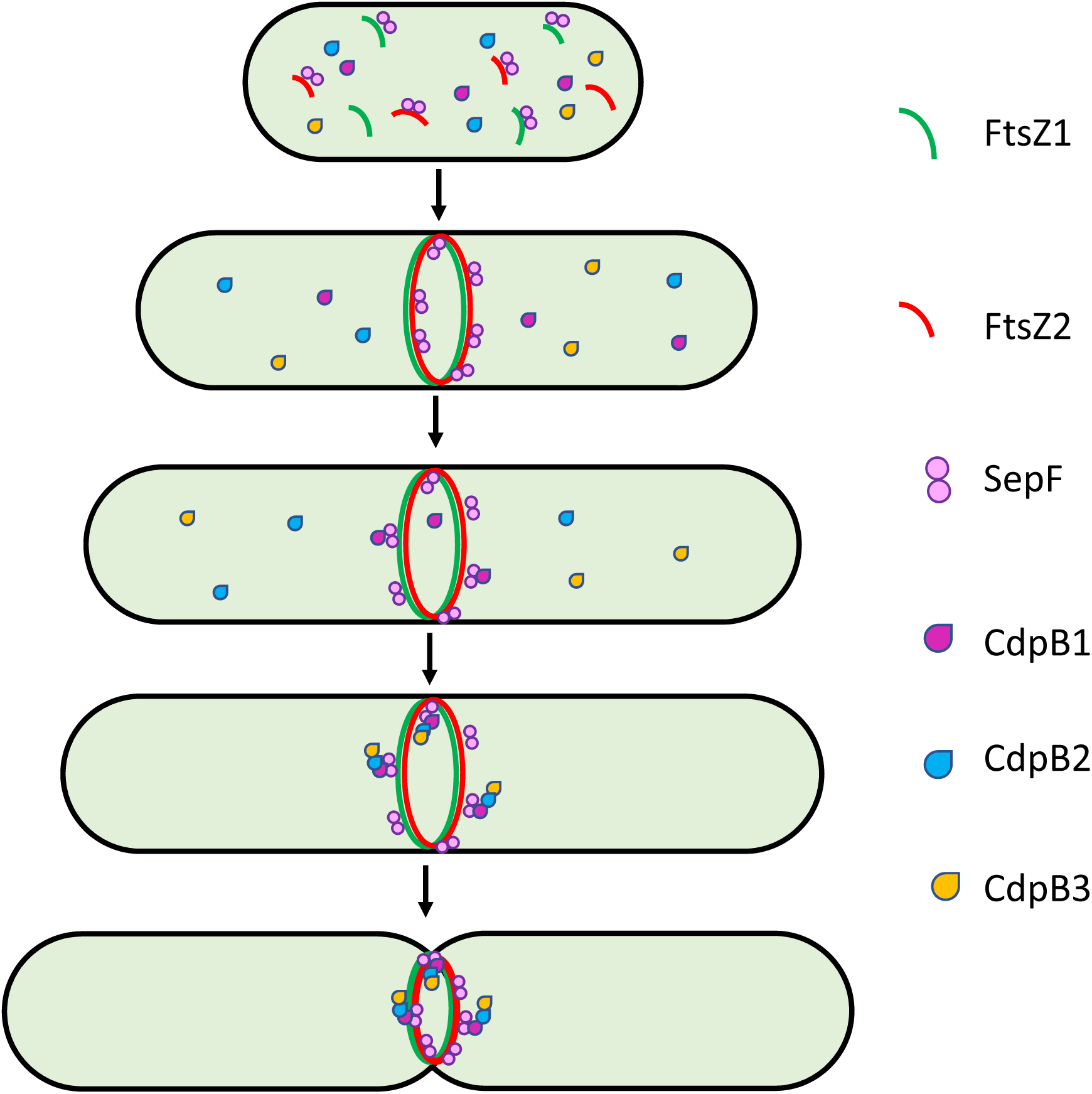
A proposed model for the functions of CdpB proteins. In *H. volcanii* cells, FtsZ1, FtsZ2 and SepF assemble into a Z ring at midcell as the cell is about the divide. CdpB1 is first recruited by SepF and itself recruits CdpB2, which then recruits CdpB3 to the Z ring. The CdpB proteins likely recruit additional cell division proteins to form the complete divisome complex. Once the divisome is fully assembled, cell membrane constricts, yielding two daughter cells. The absence of CdpB1, CdpB2 or CdpB3 causes failure to recruit additional essential or accessory division proteins to the Z ring, ultimately leading to different extents of division and cell shape defects.

Although this work establishes the CdpB proteins as important components of the FtsZ-dependent division machinery in haloarchaea, their mechanisms remain to be elucidated. CdpB1 interacts with SepF *in vivo* and *in vitro* and its depletion results in the formation of filamentous and giant cells. However, in these abnormal cells, FtsZ and SepF still clearly co-localize and in some cases, form normal looking Z rings. Thus, CdpB1 is not essential for Z ring formation but appears to be important for subsequent steps of division. In line with this conclusion, CdpB1 is required for the localization of its two paralogs, CdpB2 and CdpB3. However, given that CdpB2 and CdpB3 are not essential for cell division, their recruitment is likely not the critical function of CdpB1. It seems more plausible that CdpB1 functions as a recruiter for other essential proteins involved in cell division that currently remain unidentified. This function of CdpB1 appears widely conserved in haloarchaea as demonstrated by finding that CdpB1 orthologs from *Natrinema sp. J7* and *H. hispanica* were also involved in division and could complement CdpB1 depletion in *H. volcanii*. Future studies using CdpB1 as a bait may enable the identification of additional essential cell division proteins.

Unlike CdpB1, CdpB2 and CdpB3 are not critical for cell division in haloarchaea. Moreover, these proteins are recruited to the Z ring by CdpB1, and their overexpression could not suppress the division and morphological defects of CdpB1 depleted cells, whereas overexpression of CdpB1 suppressed the defects of CdpB2 knockout cells. These observations indicate that the three CdpB proteins perform distinct functions in archaeal cell division. The localization dependency of the CdpB proteins suggests that CdpB2 directly interacts with CdpB1 and CdpB3, likely via their PRC barrel domains because this domain has been shown to mediate protein-protein interactions^27^. However, how these interactions affect cell division, is not clear. Future studies will be necessary to characterize the interactions between the CdpB proteins and elucidate their functions in cell division.

Phylogenetic analysis of the PRC barrel domain containing proteins shows that they are widely distributed in archaea and formed three large branches. In many archaeal genomes, the genes encoding PRC barrel domain containing proteins, including CdpB1 and CdpB2, are adjacent to genes for components of the translation systems, consistent with previous predictions that the PRC barrel domain is involved in ribosome maturation and RNA metabolism^27^. However, given that at least one protein subfamily in each major clade is involved in cell division, the role of PRC barrel proteins in archaeal division is likely ancestral and the functions of PRC barrel domain diversified extensively in archaeal evolution. It is noteworthy that not all the standalone PRC barrel protein are involved in division despite pronounced similarity to CdpB proteins, such as HVO_1607 in *H. volcanii*, and NJ7G_3497 in *Natrinema sp. J7*. Many genes encoding PRC barrel domain proteins are adjacent to genes encoding Cdc48 ATPase, a subfamily of AAA+ ATPase involved in cell cycle regulation and protein degradation^42^. It remains to be tested whether the PRC barrel domain proteins in other archaea lineages are involved in cell division, cell cycle regulation, protein translation or other processes. Many bacterial genomes also harbor uncharacterized genes encoding PRC barrel domain containing proteins that are distinct from the prototypical RimM (COG0806), and their involvement in division remains to be tested.

Of special note is the presence of a PRC barrel domain in the CdvA protein, an essential component of the ESCRT-based division system. The function of the PRC barrel domain remains unknown because it is not required for the interaction of CdvA with the ESCRT-III-like CdvB protein^18^. Our finding here that the PRC barrel proteins likely function as recruiters for other cell division proteins in the FtsZ-based division system implies that the PRC barrel domain of CdvA might play a similar role in recruiting other division proteins to the ESCRT-based machinery. The identification of CdvA orthologs in *Thermoproteales*, which lack orthologs of FtsZ and ESCRT proteins and thus are thought to divide via a distinct, currently unknown mechanism^11,13^, implies that PRC barrels might be (nearly) universal components of archaeal cell division systems.

Overall, we identified the PRC barrel proteins as conserved division proteins in the archaeal FtsZ-based division system that likely function as adaptors for the recruitment of other division proteins. The wide spread distribution of the PRC barrel among archaea and in particular its presence in CdvA suggests that the role of PRC barrel domain in cell division is ancestral in archaea. Search for interaction partners of the PRC barrel proteins can be expected to advance the exploration of archaeal cell division and shed light on its evolution.

## Methods

### Strains and growth conditions

All strains used in this study were listed in Supplementary Table 3 in the Supplemental Information. *E. coli* strains were grown in LB medium (1% tryptone, 0.5% yeast extract, 0.5% NaCl and 0.05 mg/ml thymine) at indicated temperatures. When needed, ampicillin was added to a final concentration of 100 µg/mL. *H. volcanii* and *Natrinema* sp. CJ7-F strains were grown aerobically at 45°C and 200 rpm in Hv.YPC medium or Hv.Ca medium, or in media containing expanded trace elements and vitamin solution referred to as Hv.YPCTE or Hv.Cab medium^43,44^. When auxotrophic markers were used, media were supplemented with uracil (10 µg/mL or 50 µg/mL) for *ΔpyrE2* strains, or thymidine and hypoxanthine (40 µg/mL each) for *ΔhdrB* strains. Cultures were generally maintained in continuous logarithmic growth (OD_600_ < 0.8) for at least 2 days prior to sampling for analysis of mid-log cultures, unless otherwise indicated. To control gene expression via the *P_tna_* promoter, L-tryptophan (Trp) was added at the indicated concentration in cultures.

*H. hispanica* strains were cultured at 45°C in nutrient rich AS-168 medium (200 g of NaCl, 2 g of KCl, 20 g of MgSO_4_· 7H_2_O, 3 g of trisodium citrate, 1 g of sodium glutamate, 5 g of Bacto casamino acids, 5 g of yeast extract, 50 mg of FeSO_4_·7H_2_O, and 0.36 mg of MnCl_2_·4H_2_O per liter, pH 7.2) with uracil at a final concentration of 50 µg/mL. Strains carrying the expression plasmid was cultured in the modified AS-168-M medium without yeast extract to provide selection pressure.

### Plasmid construction

Plasmids used in this study were listed in Supplementary Table 4, and the primers utilized for plasmid construction were listed in Supplementary Table 5. Plasmids pTA962^45^, pIDJL40 (containing *gfp*)^43^ or pIDJL114 (containing *mCherry*)^24^ were used as the backbones to construct plasmids for controlled expression of the *cdpB* genes or modified versions in *H. volcanii.* The *cdpB* ORFs were amplified and cloned between the NdeI and BamHI sites of pIDJL40 or pIDJL114 to create *-gfp* or *-mCherry* fusions, respectively. Additionally, plasmid pZS214 was created, allowing expression of CdpB1-GFP under the control of the native promoter of *cdpB1*. Similarly, NJ7G_3475 and HAH_1390 (homolog of CdpB1) were inserted into plasmids pFJ6-*P_tna_* and pWL502^46^, respectively, to construct fluorescent fusions in *Natrinema sp.* CJ7-F and *H. hispanica* strains.

For dual expression of the various division genes, a fragment containing *ftsZ-mCherry* or *sepF-mCherry* was ligated into the NotI-cut (klenow blunt end) of the above plasmids (containing *-gfp* or –*mCherry* fusion). In order to be used in the depletion strains, the *P_tna_* promoter of the above plasmids was replaced by the *P_phaR_* promoter, which is constitutively active.

To detect the interaction between CdpB1 and other cell division proteins, we applied the tripartite split-GFP system^33^ in *H. volcanii.* The *sfGFP10* or *sfGFP11* fragment was fused to the respective reading frames encoding the cell division proteins. The *sfGFP* fragments and cell division proteins coding sequences are separated by two kinds of flexible linkers. The longer linkers of 30-mer (GFP10 tag) and 25-mer (GFP11 tag) were used for interaction assays of FtsZ1 and FtsZ2, respectively. The shorter linkers of 15-mer (GFP10 tag) and 17-mer (GFP11 tag) were used for interaction assays of CdpB1 and SepF. We constructed intermediate vectors carrying *sfGFP1-9*, *sfGFP10* and *sfGFP11* with different linkers under the control of *P_tna_*. The three cassettes were separated by several restriction enzyme sites including EcoRI, HindIII, XbaI and NheI. The target genes were cloned to the restriction sites to fuse with the *sfGFP10* or *sfGFP11* fragments in different directions. A series of plasmids and corresponding control plasmids were constructed to examine the interaction between CdpB1 and other cell division proteins.

For detailed construction of every plasmid, please check the Supplementary Information. All plasmids were demethylated by passage through *E. coli* JM110 and re-purified prior to transfer to haloarchaea by PEG-mediated spheroplast transformation^30^.

### Genomic modification

To construct the depletion strain of *cdpB1* in *H. volcanii,* a non-replicating plasmid was first constructed that can recombine at the *cdpB1* locus using two-step homologous recombination thereby replacing the promoter region of *cdpB1* with the specific tryptophan-inducible *P_tna_* promoter^31^. The upstream and downstream flanking sequences on either side of *cdpB1*’s start codon were PCR amplified from *H. volcanii* DS70 genomic DNA (upstream flank) and plasmid pZS103 (contains the L11e transcription terminator followed by the *P_tna_::cdpB1-gfp* cassette) with primers listed in Supplementary Table 5, respectively. The upstream and downstream fragments were joined by overlap-extension PCR and the products were digested with HindIII and BamHI and then ligated to pTA131^30^ (at HindIII-BamHI), giving rise to pZS98. The fragment *P_fdx_::hdrB* from pTA1185^24^ was inserted between the upstream and downstream fragments of pZS98 at the SphI site. Clones containing the *P_fdx_::hdrB* oriented with the downstream *P_tna_::cdpB1* cassette were selected and named pZS111. Demethylated pZS111 was transformed into *H. volcanii* H98^30^ (DS70, *ΔpyrE2 ΔhdrB*) and transformants were selected on agar medium without uracil. The resultant colonies were expected to contain the plasmid integrated between the upstream or downstream of the genomic *cdpB1* locus by a single-crossover (“pop-in”). After growth of single colonies in liquid Hv.YPC media, cells were plated onto Hv.Ca agar containing 100 μg/mL of uracil, 50 μg/mL of 5-fluoroorotic acid (FOA) and 1 mM Trp to select for excision of the plasmid (“pop-out”). Single colonies were streaked onto the same medium and arising colonies were then screened by allele-specific PCR and sequencing. Colonies containing *P_tna_::cdpB1* as the only copy of *cdpB1* were saved and named as HZS1 (Supplementary Table 3). The SepF depletion strain HZS2 (H98, *P_tna_::sepF*) was constructed similarly.

The construction process for the *nj7g_3475* and *hah_1390* depletion strains in *Natrinema* sp. CJ7-F^47^ and *H. hispanica* DF60^48^, respectively, was similar to the above procedure for *cdpB1* in *H. volcanii*. The upstream fragment (upstream of *nj7g_3475*) and downstream fragments (the L11e transcription terminator and *P_tna_::nj7g_3475* cassette) were amplified, joined by overlap-extension PCR, and the products were digested with BamHI and AflII and ligated to pNBK-F^49^ (at BamHI-AflII), giving rise to pZS253 (*P_tna_*::*nj7g_3475*). Similarly, the upstream fragment (upstream of *hah_1390*) and downstream fragment (containing the L11e transcription terminator and *P_tna_::hah_1390* cassette) of *hah_1390* were amplified, joined by overlap-extension PCR and inserted into plasmid pHAR^48^ (at KpnI and HindIII) to obtain vector pZS280 (*P_tna_*::*hah_1390*). Demethylated pZS253 and pZS280 were transformed into *Natrinema* sp. CJ7-F and *H. hispanica* DF60 strain separately to generate NJ7G_3475 depletion strain in *Natrinema sp. J7*-F and HAH_1390 depletion strain in *H. hispanica,* respectively.

*cdpB2* and *cdpB3* deletion strains were also constructed by the pop-in/pop-out approach^31^ as above. The upstream fragment and downstream fragments of *cdpB2* were amplified, joined by overlap-extension PCR, and the products were digested with HindIII and BamHI and ligated to pTA131^30^ (at HindIII-BamHI), giving rise to a non-replicating plasmid pZS398. Similarly, a non-replicating plasmid pZS399 containing the upstream and downstream flanking sequences of *cdpB3* was constructed. The two respective non-replicating plasmids pZS398 and pZS399 were transformed into strain H26^30^ (DS70, *ΔpyrE2*) to generate the desired deletion strains, respectively.

### Construction of the genomic library of *H. volcanii*

To construct a genomic library of *H. volcanii*, the genomic DNA of *H. volcanii* was digested with Sau3AI and TaqI, fragments of about 1-5 kb were purified and then ligated into a derivative of pTA1228 (carrying the tryptonphan-inducible promoter *P_tna_*) digested with BamHI and ClaI. Ligation products were transformed into competent *E. coli* and transformants selected on LB plates with ampicillin. The plasmids from ten of the transformants were isolated and cut with restriction enzymes to determine if most of the plasmids contained genomic DNA fragments of *H. volcanii*. About 40,000 transformants were pooled together and the plasmids were extracted and saved as the library.

### Rationale for the screen for cell division proteins in *H. volcanii*

In bacteria, overproduction of proteins involved in essential cellular processes often impair cell growth by causing a malfunctioning of the corresponding machineries or disruption of metabolic pathways. For example, overexpression of cell division proteins or proteins regulating cell division often results in a division block and thereby preventing colony formation^50–54^. As haloarchaea also divide in an FtsZ-dependent manner^8,11,24^, we hypothesized that overexpression of haloarchaeal cell division proteins might interfere with cell division and inhibit cell growth. In line with this hypothesis, overexpression of CdrS, the master regulator of haloarcheal cell cycle, has been shown to cause severe cell division and morphological defects in multiple halophiles^29^. Thus, we screened for proteins whose overexpression was toxic to the cell in *H. volcanii* with a hope that some might cause division and morphological changes. To do this, we transformed the genomic library of *H. volcanii* into H26 and transformants were screened on plates with or without tryptophan by replica plating. Transformants displaying a growth defect on plates with tryptophan were confirmed for the tryptophan-dependent growth defect. The inserts in the selected transformants were determined by sequencing and then re-cloned into pTA1228 to confirm its toxicity to the cell and the effect on cell morphology in the presence of tryptophan. Using this approach, we found that many DNA segments inserted into pTA1228 blocked colony formation in the presence of tryptophan, but few would cause morphological changes to the cells, including one that harbored the first 82 amino acids of SepF. We also found that a fragment containing a part of HVO_1691 (aa1-56) inhibited the growth of *H. volcanii*, leading to its discovery. This indicated that this approach was working, although not very effective.

### Fluorescence microscopy

All phase contrast and fluorescence images were acquired using an Olympus BX53 upright microscopes with a Retiga R1 camera from QImaging, a CoolLED pE-4000 light source and a U Plan XApochromat phase contrast objective lens (100X, 1.45 numerical aperture [NA], oil immersion). Green and red fluorescence was imaged using the Chroma EGFP filter set EGFP/49002, mCherry/Texas Red filter set mCherry/49008, respectively. For microscopy, a 2 μL sample of cells were immobilized on 1.5% agarose pads equilibrated with 18% BSW (Hv.Ca medium without casamino acids and CaCl_2_) at room temperature, and a clean glass coverslip placed on top.

#### (1) Localization of CdpB proteins and HVO_1607 in *H. volcanii*

Overnight cultures of *H. volcanii* H26 carrying plasmid pZS103 (*P_tna_::cdpB1-gfp*), pZS101 (*P_tna_::cdpB1-mCherry*), pZS336 (*P_tna_::cdpB2-gfp*), pZS337 (*P_tna_::cdpB3-gfp*), pZS422 (*P_tna_::hvo_1607-gfp*) or pZS423 (*P_tna_::gfp-hvo_1607*) were diluted 1:100 in fresh Hv.Cab medium with 0.2 mM Trp, and grown at 45°C to OD_600_ about 0.2. 2 μL of the cultures was spot on BSW agarose pads for photograph.

#### (2) Co-localization of CdpB1-GFP with FtsZ1, FtsZ2 and SepF

Exponential phase cultures of *H. volcanii* H26 carrying plasmid pZS105 (*P_tna_::cdpB1-gfp*-*ftsZ1-mCherry*), pZS106 (*P_tna_::cdpB1-gfp*-*ftsZ2-mCherry*) or pZS107 (*P_tna_::cdpB1-gfp*-*sepF-mCherry*) were treated as in **(1)** to observe co-localization of protein fusions.

#### (3) Localization dependency of CdpB1

Overnight cultures of ID56^24^ (H98, *P_tna_::ftsZ1*) harboring plasmid pZS284 (*P_phaR_::*c*dpB1-gfp*-*ftsZ2-mCherry*) or pZS285 (*P_phaR_::*c*dpB1-gfp*-*sepF-mCherry*), ID57^24^ (H98, *P_tna_::ftsZ2*) harboring plasmid pZS239 (*P_phaR_::cdpB1-gfp*-*ftsZ1-mCherry*) or pZS285 (*P_phaR_::*c*dpB1-gfp*-*sepF-mCherry*), and HZS2 (H98, *P_tna_::sepF*) harboring plasmid pZS239 (*P_phaR_::cdpB1-gfp*-*ftsZ1-mCherry*) or pZS284 (*P_phaR_::*c*dpB1-gfp*-*ftsZ2-mCherry*) were diluted 1:100 in fresh Hv.Cab medium with 1mM Trp, and grown at 45°C to OD_600_ about 0.4. Cells were then collected by centrifugation and washed three times with fresh Hv.Cab medium to remove the tryptophan, followed by resuspension in the same volume of Hv.Cab medium. The tryptophan-free culture was then inoculated 1:100 in Hv.Cab medium with or without 1 mM tryptophan and cultured to OD_600_ about 0.2. 2 μL of the cultures was spot on BSW agarose pads for photograph. To check the localization of CdpB1-GFP in the *ftsZ1* and *ftsZ2* double deletion strain ID112^24^ (H98, *ΔftsZ1 ΔftsZ2*), overnight culture of ID112 carrying plasmid pZS103 (*P_tna_::cdpB1-gfp*) was diluted 1:100 in fresh Hv.Cab medium with 1mM Trp, and grown at 45°C to OD_600_ about 0.2. 2 μL of the cultures was spot on BSW agarose pads for photograph.

#### (4) Localization Interdependence of CdpB1, CdpB2 and CdpB3

Overnight cultures of HZS1 (H98, *P_tna_::cdpB1*) harboring plasmid pZS408 (*P_phaR_::*c*dpB2-gfp*-*sepF-mCherry*) or pZS407 (*P_phaR_::*c*dpB3-gfp*-*sepF-mCherry*), were treated as in **(3)** to examine the localization dependency on CdpB1 of CdpB2 and CdpB3. To check the localization of proteins in the *ΔcdpB2* and *ΔcdpB3* cells, overnight culture of HZS5 (H26, *ΔcdpB2*) harboring plasmid pZS417 (*P_phaR_::* c*dpB3-gfp*-*cdpB1-mCherry*), and HZS6 (H26, *ΔcdpB3*) harboring plasmid pZS418 (*P_phaR_::* c*dpB2-gfp*-*cdpB1-mCherry*) were diluted 1:100 in fresh Hv.Cab medium, and grown at 45°C to OD_600_ about 0.2. 2 μL of the cultures was spot on BSW agarose pads for photograph.

#### (5) Co-localization of FtsZ1, FtsZ2 and SepF in CdpB1 depleted cells

Overnight cultures of HZS1 (H98, *P_tna_::cdpB1*) carrying plasmid pZS289 (*P_phaR_::ftsZ1-gfp*-*ftsZ2-mCherry*), pZS322 (*P_phaR_::sepF-gfp*-*ftsZ1-mCherry*) or pZS324 (*P_phaR_::sepF-gfp*-*ftsZ2-mCherry*) were treated as in **(3)** to examine the localization of FtsZ1, FtsZ2 and SepF in CdpB1 depleted cells.

#### (6) Localization of the PRC barrel proteins in *Natrinema sp CJ7-F* and *H. hispanica*

Overnight cultures of *Natrinema sp.* CJ7-F carrying plasmid pZS217 (*P_tna_::NJ7G_3475-gfp*), pZS384 (*P_tna_:: NJ7G_2729-gfp*) or pZS385 (*P_tna_:: NJ7G_3497-gfp*) were diluted 1:100 in fresh Hv.Cab medium with 0.2 mM Trp, and grown at 45°C to OD_600_ about 0.2. Overnight cultures of *H. hispanica* DF60 carrying plasmid pZS306 (*P_tna_::HAH_1390-gfp*), pZS388 (*P_tna_:: HAH_0460-gfp*) or pZS389 (*P_tna_:: HAH_5240-gfp*) were diluted 1:100 in fresh AS-168-M medium with 0.2 mM Trp, and grown at 45°C to OD_600_ = 0.2. 2 μL of the cultures was spot on BSW agarose pads for photograph.

#### (7) Split-FP assay

Overnight cultures of *H. volcanii* H26 carrying the tripartite split-GFP system plasmids were diluted 1:100 in fresh Hv.Cab medium with 0.2 mM Trp, and cultivated at 45°C overnight followed by cultivation at 37°C for 3h. 2 μL of the culture was spot on BSW agarose pad for photograph.

### Quantification of Fluorescence

To quantify the protein-protein interaction signal between CdpB1 and other cell division proteins by Split-FP, the fluorescence of the *H. volcanii* transformants was quantified. In each case, 5 mL culture was cultivated at 45°C to an optical density of OD_600_ = 1-1.5. The culture was then brought to OD_600_=1 and kept shaking at 30°C overnight with 0.2 mM Trp. 1 mL of the culture was harvested by centrifugation (12000 × g, 2 min), washed and brought to OD_600_ = 1 with 18% BSW. 200 μL sample was analyzed in a 96-well plate and evaluated using the Varioskan LUX multifunctional microplate detection system. All experiments were performed with two biological samples and three technical replicates. The p-values were calculated using Student t-test.

### Protein expression and purification

The proteins were produced by heterologous expression in the *E. coli* strain BL21 (DE3) harboring plasmid pZS311 (H-SUMO-CdpB1) or pZS288 (H-SUMO-SepF). An overnight culture of each strain grown in LB with ampicillin (100 μg/mL) was diluted 1:100 into 300 mL fresh LB medium supplemented with ampicillin (100 μg/mL) and incubated at 37°C until OD_600_ reached about 0.4. IPTG was then added to the culture to a final concentration of 1 mM and incubated at 37°C for another 3h. Cells were collected by centrifugation, washed with 10 mM Tris-HCl (pH 7.9), and frozen at −80°C until used. On the day of purification, the cells were thawed and resuspended in 20 mL high salt lysis buffer (25 mM Tris HCl [pH 7.5], 2.5 M KCl, 5% glycerol, 0.1 mM dithiothreitol (DTT) and 20 mM imidazole) and lysed by sonication. The lysates were centrifuged at 12,000 rpm for 15 min at 4°C to remove cell debris. The supernatants were loaded onto pre-equilibrated Ni-NTA agarose. The column was washed once with high salt lysis buffer. The bound protein was eluted with elution buffer (25 mM Tris HCl [pH 7.5], 2.5 M KCl, 5% glycerol, 0.1 mM dithiothreitol (DTT) and 250 mM imidazole). Fractions were analyzed by SDS-PAGE gel and the ones with highest concentration of protein were pooled and dialyzed against storage buffer (25 mM Tris HCl [pH7.5], 2.5 M KCl, 5% glycerol, 0.1 mM dithiothreitol (DTT)), aliquoted and stored at −80°C.

The H-SUMO tag of CdpB1 was cleaved with purified 6xHis-tagged SUMO protease (Ulp1) for 1h at 30°C in the protein storage buffer with 200 mM KCl. The released tag and protease were removed by passing the reaction mixture through the pre-equilibrated Ni-NTA agarose. Untagged CdpB1 was collected in the flow through, dialyzed against protein storage buffer, concentrated and stored at −80°C.

### Pull-down assay

The pull-down assay was performed at 4°C. To test the interaction between H-SUMO-SepF and CdpB1, 50 μg of H-SUMO-SepF and 50 μg of purified CdpB1 were mixed in a total volume of 400 μL equilibrium buffer (25 mM Tris HCl [pH7.5], 2.5 M KCl, 5% glycerol, 0.1 mM dithiothreitol (DTT)), incubated at 4°C for 2h and then loaded into a gravity flow column with 200 μL pre-equilibrated Ni-NTA agarose. After incubation on ice for 10 min without agitation, the mixture was allowed to pass through the column by gravity. The column was then washed with 400 μL of wash buffer (25 mM Tris HCl [pH 7.5], 2.5 M KCl, 5% glycerol, 0.1 mM dithiothreitol (DTT) and 20 mM imidazole) twice. Proteins bound to the Ni-NTA beads were eluted with 400 μL of elution buffer (25 mM Tris HCl [pH 7.5], 2.5 M KCl, 5% glycerol, 0.1 mM dithiothreitol (DTT) and 250 mM imidazole). All fractions were collected during the procedure and analyzed by SDS-PAGE.

### Immunoprecipitation, western blot procedures and antibodies

Overnight cultures of *H. volcanii* carrying the expression plasmids were diluted 1:100 in 40 mL fresh Hv.YPC medium, and cultivated at 45°C to OD_600_ about 1.0. Cells were collected by centrifugation at 10,000 rpm for 10 min and resuspended in 2 mL high salt lysis buffer (25 mM Tris HCl [pH 7.5], 2.5 M KCl, 5% glycerol, 0.1 mM dithiothreitol (DTT) and 20 mM imidazole) containing an anti-protease cocktail (MCE) and lysed by sonication. The lysates were centrifuged at 12,000 rpm for 5 min at 4°C to remove cell debris. 400 μL of the supernatant was added to pre-prepared Ab-coated magnetic beads and incubated overnight at 4 °C. Magnetic beads-Ab-protein complexes were separated by centrifugation and then washed with 400 μL high salt wash buffer (25 mM Tris HCl [pH 7.5], 2.5 M KCl, 5% glycerol, 0.1 mM dithiothreitol (DTT), 20 mM imidazole, and 0.5% Tween-20) 5 times. The immunocomplexes were finally eluted with boiling SDS-PAGE Loading Buffer and were separated by SDS-PAGE. Following transfer onto NC membranes, proteins were revealed by immunoblot. The following antibodies, with their respective dilutions in 5% skimmed milk, were used: anti-GFP (AE078, ABclonal) 1/2,000, anti-Flag (AE004, ABclonal) 1/2,000, anti-GFP (HT801-01, Transgen) 1/10,000, anti-Flag (HT201-01, Transgen) 1/10,000, anti-mouse secondary antibody (HS201-01, Transgen) 1/10,000, anti-rabbit secondary antibody (HS101-01, Transgen) 1/10,000.

### Sequence comparison, phylogenetic analysis and gene neighborhood analysis for archaeal PRC barrel domain containing proteins

The arCOG database^55,56^ that includes annotated clusters of orthologous genes for 524 archaeal genomes covering all major archaeal lineages is available at https://ftp.ncbi.nih.gov/pub/wolf/COGs/arCOG/tmp.ar18/. PSI-BLAST^57^ search (e-value cutoff of 0.01, effective database size of 2*10^7^, no composition-based statistics and no low complexity filtering, 5 iterations) with several selected query sequences from each arCOG consisting of PRC barrel domain proteins (2155, 2157, 2158, 5740, 8023) was used to run searches against all proteins in the arCOG database to identify remotely similar homologs. Proteins identified using this approach but currently not annotated as containing the PRC barrel domain were additionally searched against PFAM, CDD and PDB profiles databases using HHpred^58^. If HHpred searches revealed similarity with known PRC barrel protein profiles with probability greater than 80%, then the query sequences and the respective arCOGs were assigned to the PRC barrel family (Supplementary Table 1). Muscle5 program^59^ with default parameters was used to construct a multiple sequence alignment of archaeal PRC barrel domains. For phylogenetic analysis, several poorly aligned sequences or fragments were discarded, and the remaining protein sequences were realigned. Columns in the multiple alignment were filtered for homogeneity value^60^ 0.05 or higher and gap fraction less than 0.667. This filtered alignment was used as an input for FastTree program^61^ to construct an approximate maximum likelihood phylogenetic tree with the WAG evolutionary model and gamma-distributed site rates (Supplementary File 1). The same program was used to calculate support values. HHpred and Marcoil^62^ were used to search for sequence similarity and prediction of coiled-coil regions, respectively, for protein domains fused to PRC barrel domain. For genome context analysis and search for putative operons, neighborhoods containing five upstream and five downstream genes were constructed for all identified genes encoding PRC barrel domain proteins (Supplementary Table 2).

## Data availability

Data generated and analyzed during this study are presented in the paper or in the supplementary information. Plasmids and strains that support the findings of this study are available from the corresponding authors on reasonable request.

## Acknowledgements

We thank members of the Du lab, Koonin lab, Chen lab and Krupovic lab for advice and helpful discussions to carry out this study. We thank Dr. Yan Liao and Dr. Iain Duggin at University of Technology Sydney for sending us the *ftsZ* depletion/deletion strains and plasmids for construction of fluorescent protein fusions. We thank Dr. Thorsten Allers at University of Nottingham and Dr. Xipeng Liu at Shanghai Jiao Tong University for sending us the *H. volcanii* strains and plasmids. We thank Dr. Ming Li and Dr. Hua Xiang at the Institute of Microbiology Chinese Academy of Sciences for providing us with the *H. hispanica* strains and plasmids. We would also like to thank Dr. Junfeng Liu and Yunfeng Yang at Shandong University for insightful discussions on the function of the PRC barrel domain of CdvA. This study was supported by National Natural Science Foundation of China (grant 32270049 and 32070032, http://www.nsfc.gov.cn/), the Fundamental Research Funds for the Central Universities (grant 2042021kf0198) and Wuhan University (https://www.whu.edu.cn/) to S.D.; K.S.M’s and E.V.K’s research is supported by the Intramural Research Program at the National Library of Medicine.

## Extended Data

**Extended Data Fig. 1.**
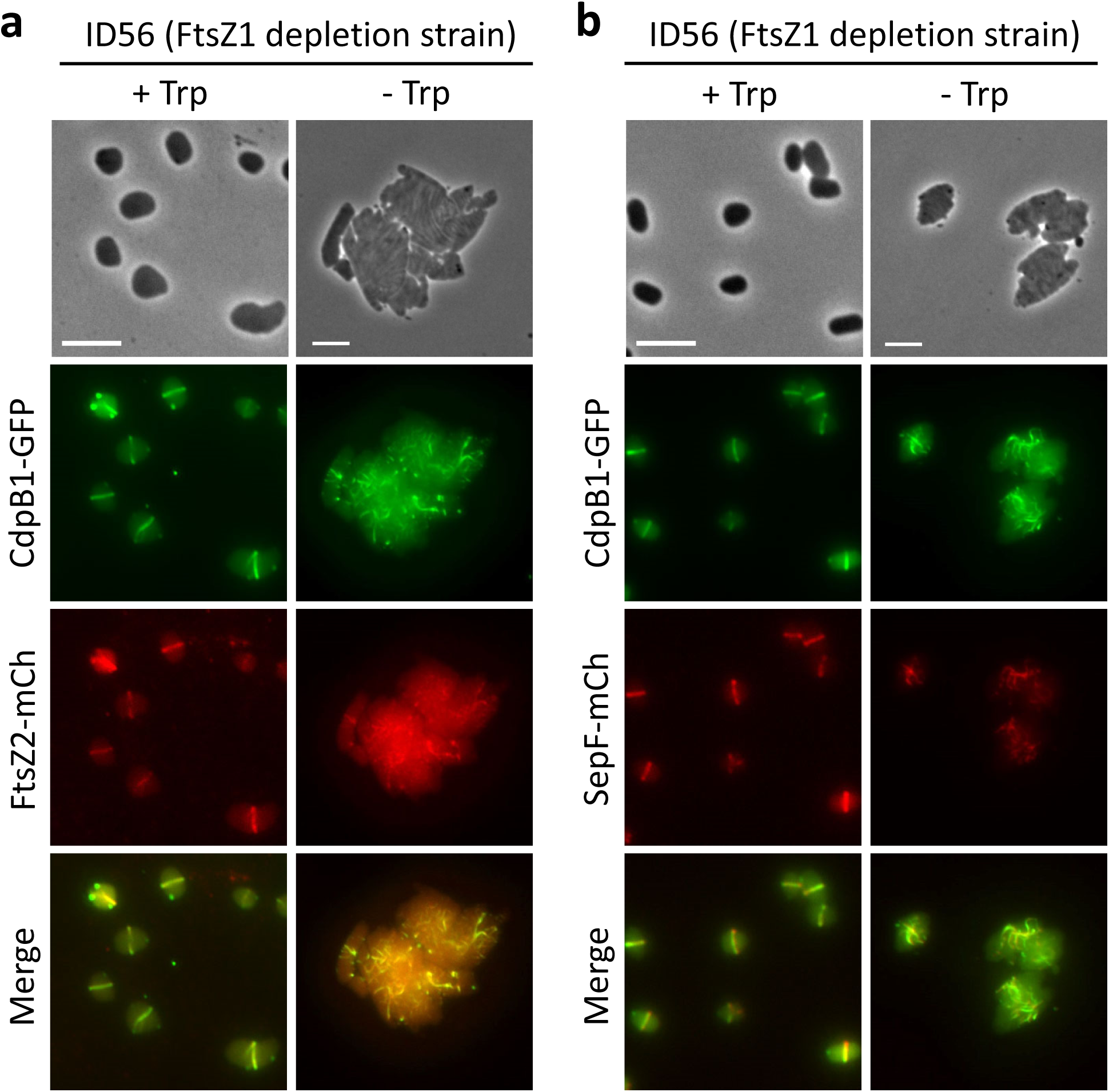
CdpB1 does not depend on FtsZ1 for co-localization with FtsZ2 and SepF. Exponential phase cultures of ID56 (H98, *P_tna_::ftsZ1*) harboring plasmid pZS284 (*P_phaR_::*c*dpB1-gfp*-*ftsZ2-mCherry*) or pZS285 (*P_phaR_::*c*dpB1-gfp*-*sepF-mCherry*) grown in Hv.Cab medium with or without tryptophan were treated as in Fig. 1b for phase-contrast and fluorescence microscopy. **a.** CdpB1 co-localizes well with FtsZ2 in the absence of FtsZ1. **b.** CdpB1 co-localizes well with SepF in the absence of FtsZ1. Scale bar 5 μm.

**Extended Data Fig. 2.**
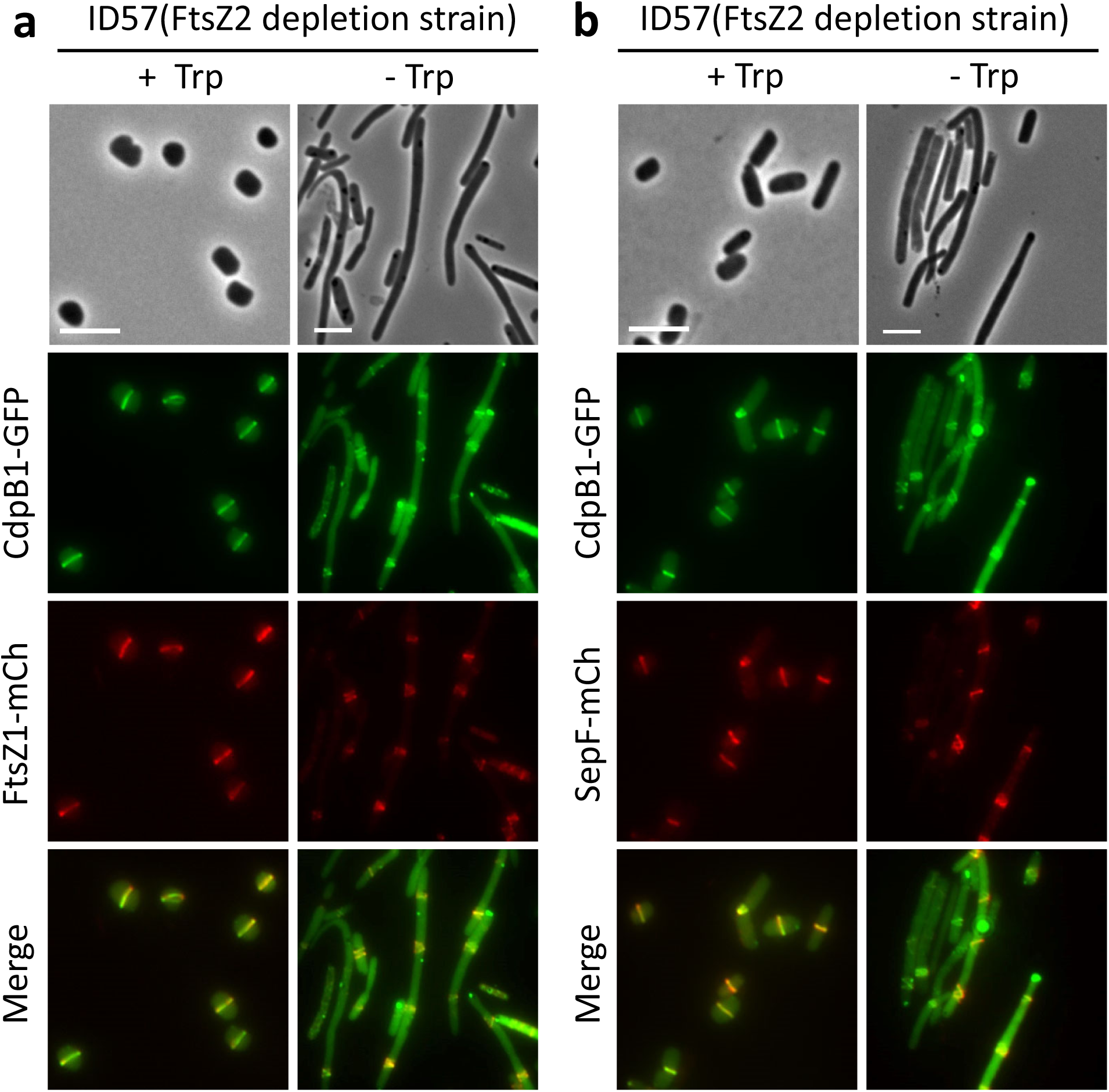
CdpB1 does not depend on FtsZ2 for co-localization with FtsZ1 and SepF. Exponential phase cultures of ID57 (H98, *P_tna_::ftsZ2*) harboring plasmid pZS239 (*P_phaR_::cdpB1-gfp*-*ftsZ1-mCherry*) or pZS285 (*P_phaR_::*c*dpB1-gfp*-*sepF-mCherry*) grown in Hv.Cab medium with or without tryptophan were treated as in Fig. 1b for phase-contrast and fluorescence microscopy. **a.** CdpB1 co-localizes well with FtsZ1 in the absence of FtsZ2. **b.** CdpB1 co-localizes well with SepF in the absence of FtsZ2. Scale bar 5 μm.

**Extended Data Fig. 3.**
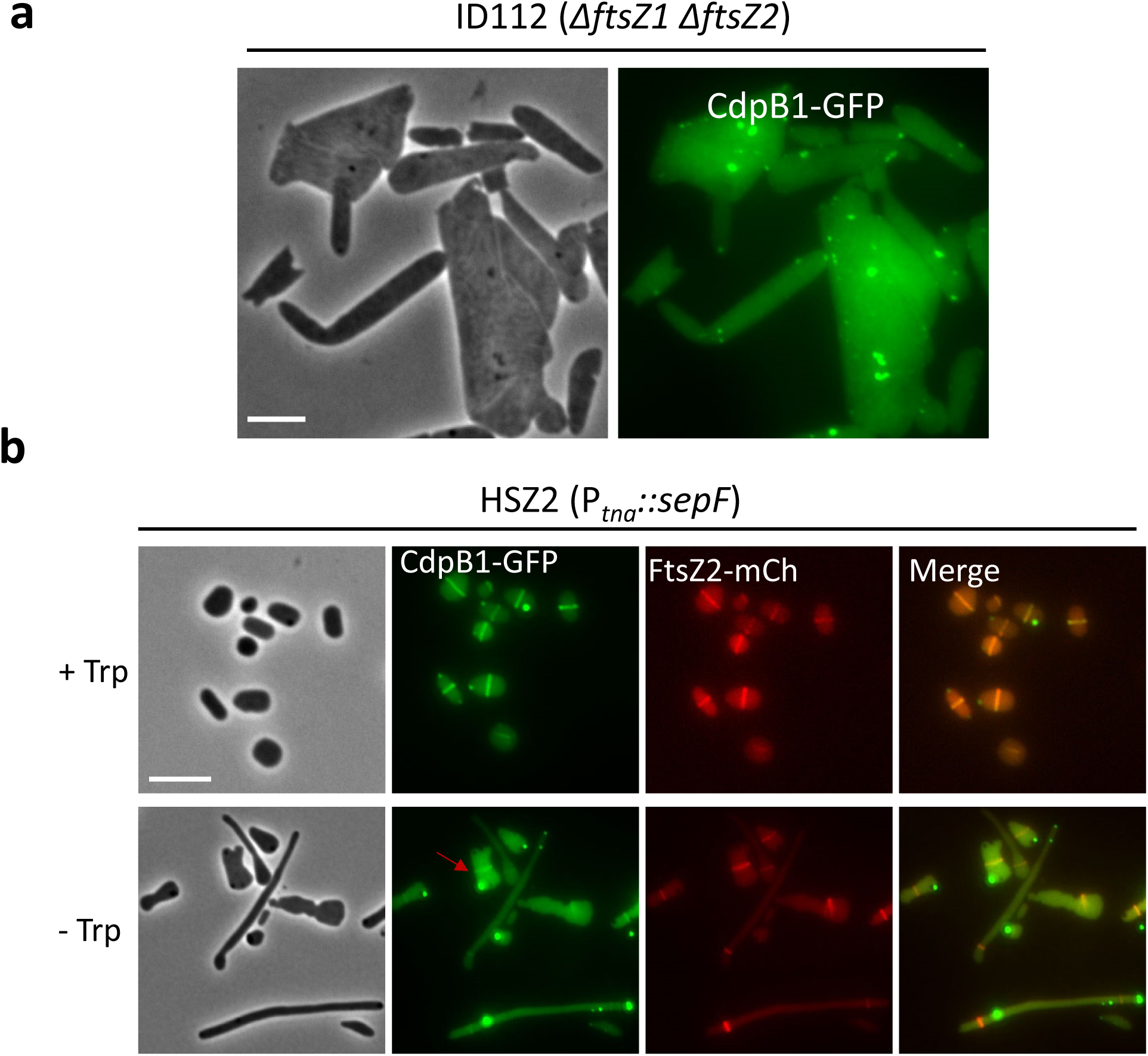
CdpB1 depends on the presence of both FtsZ1 and FtsZ2, and SepF for correct localization. **a.** CdpB1 depends on FtsZs for localization. Exponential phase cultures of ID112 (H98, Δ*ftsZ1*Δ*ftsZ2*) harboring plasmid pZS103 (*P_tna_::cdpB1-gfp*), were treated as in Fig. 1a for phase-contrast and fluorescence microscopy. **b.** CdpB1 depends on SepF for localization. Exponential phase culture of strain HZS2 (H98, *P_tna_::sepF*) harboring plasmid pZS284 (*P_phaR_:: cdpB1-gfp*-*ftsZ2-mCherry*) were treated as in Fig. 1b. 2 μL of the cultures was spotted on a BSW agarose pad for phase-contrast and fluorescence microscopy. Scale bar 5 μm.

**Extended Data Fig. 4.**
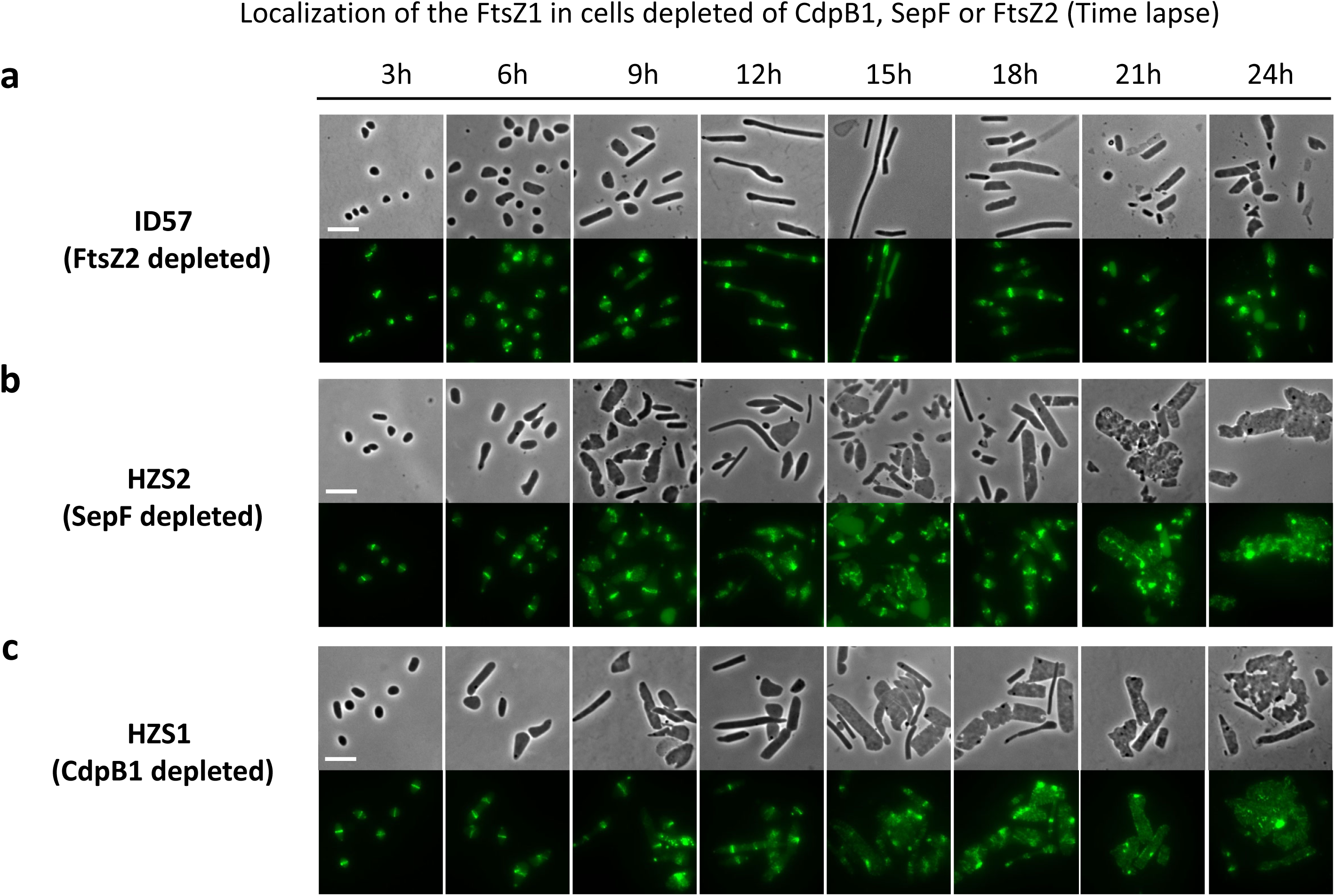
Cell morphology and FtsZ1 localization after depletion of FtsZ2, SepF or CdpB1. Exponential phase cultures of ID57 (H98,*P_tna_::ftsZ2*), HZS2 (H98, *P_tna_::sepF*) or HZS1 (H98, *P_tna_::cdpB1*) harboring plasmid pZS208 (*P_native_:: ftsZ1-gfp*) were treated as in Fig. 1b for phase-contrast and fluorescence microscopy at different time points post removal of tryptophan. Scale bar 5 μm.

**Extended Data Fig. 5.**
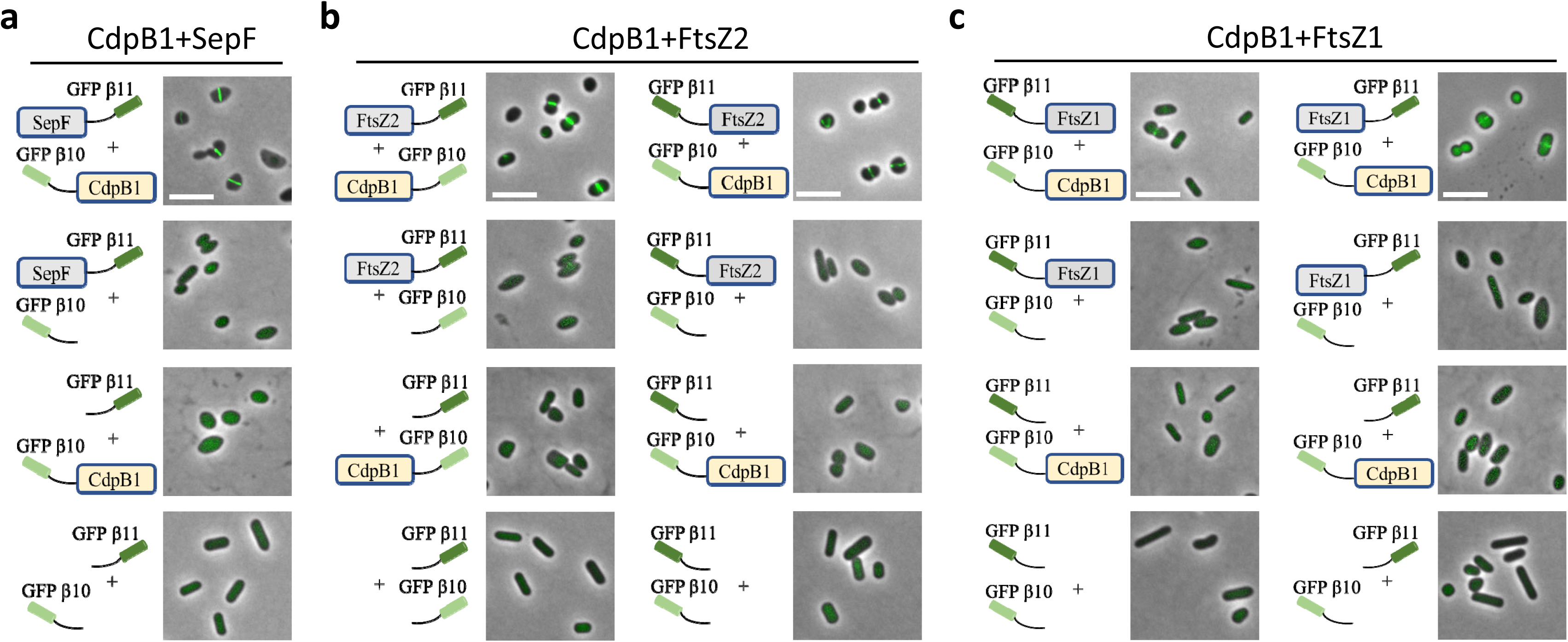
CdpB1 displays interaction with SepF and FtsZ2 as well as FtsZ1 by Slit-FP assay. An exponential culture of strain H26 (DS70, Δ*pyrE2*) harboring the split-FP plasmid expressing the indicated protein(s) was treated as in Fig. 3a and the fluorescence signal and localization were observed by microscopy. **a**. CdpB1 interacts with SepF. **b**. CdpB1 interacts with FtsZ2. **c**. CdpB1 displays weak interaction with FtsZ1. Scale bar 5 μm.

**Extended Data Fig. 6.**
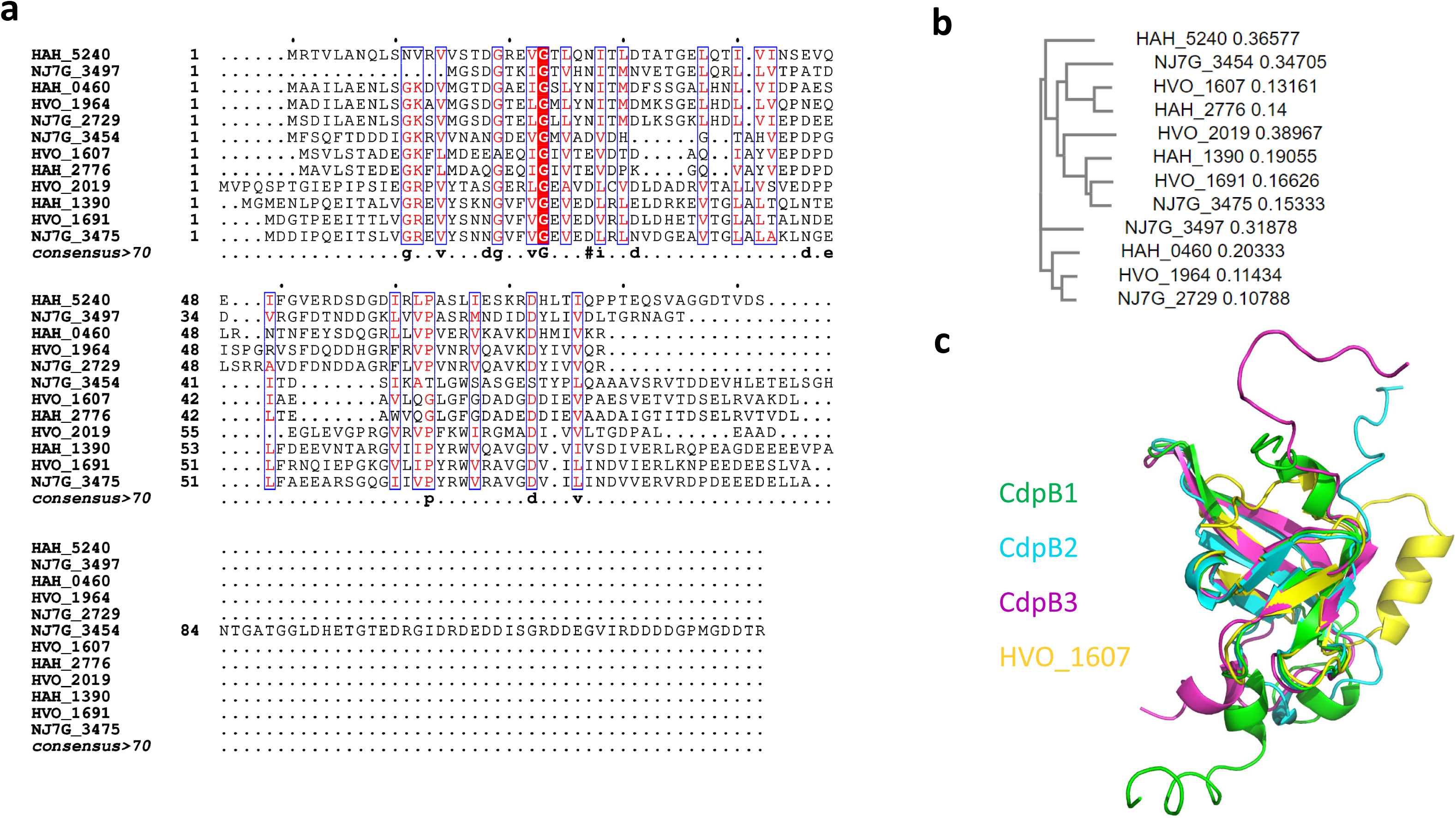
Alignment of CdpB1 with its paralogs and homologs from other haloarchaea and their predicted structures. **a.** Amino sequences of PRC barrel domain containing proteins from *H. volcanii*, *Natrinema sp. J7-1* and *H. hispanica* were downloaded from Uniprot: https://www.uniprot.org/, aligned by Clustal Omega: https://www.ebi.ac.uk/Tools/msa/clustalo/ and then depicted using ESPRIPT: http://espript.ibcp.fr/. **b**. Phylogenetic tree of the RC-barrel domain containing proteins from *H. volcanii*, *Natrinema sp. J7-1* and *H. hispanica* generated by Clustal Omega. **c**. Superimposition of the predicted structures of CdpB1, CdpB2,CdpB3 and HVO_1607 of *H. volcanii*. Structure models were generated by AlphaFold and were downloaded from Uniprot and aligned by PyMOL. CdpB1, green; CdpB2, cyan; CdpB3, magenta; HVO_1607, yellow.

**Extended Data Fig. 7.**
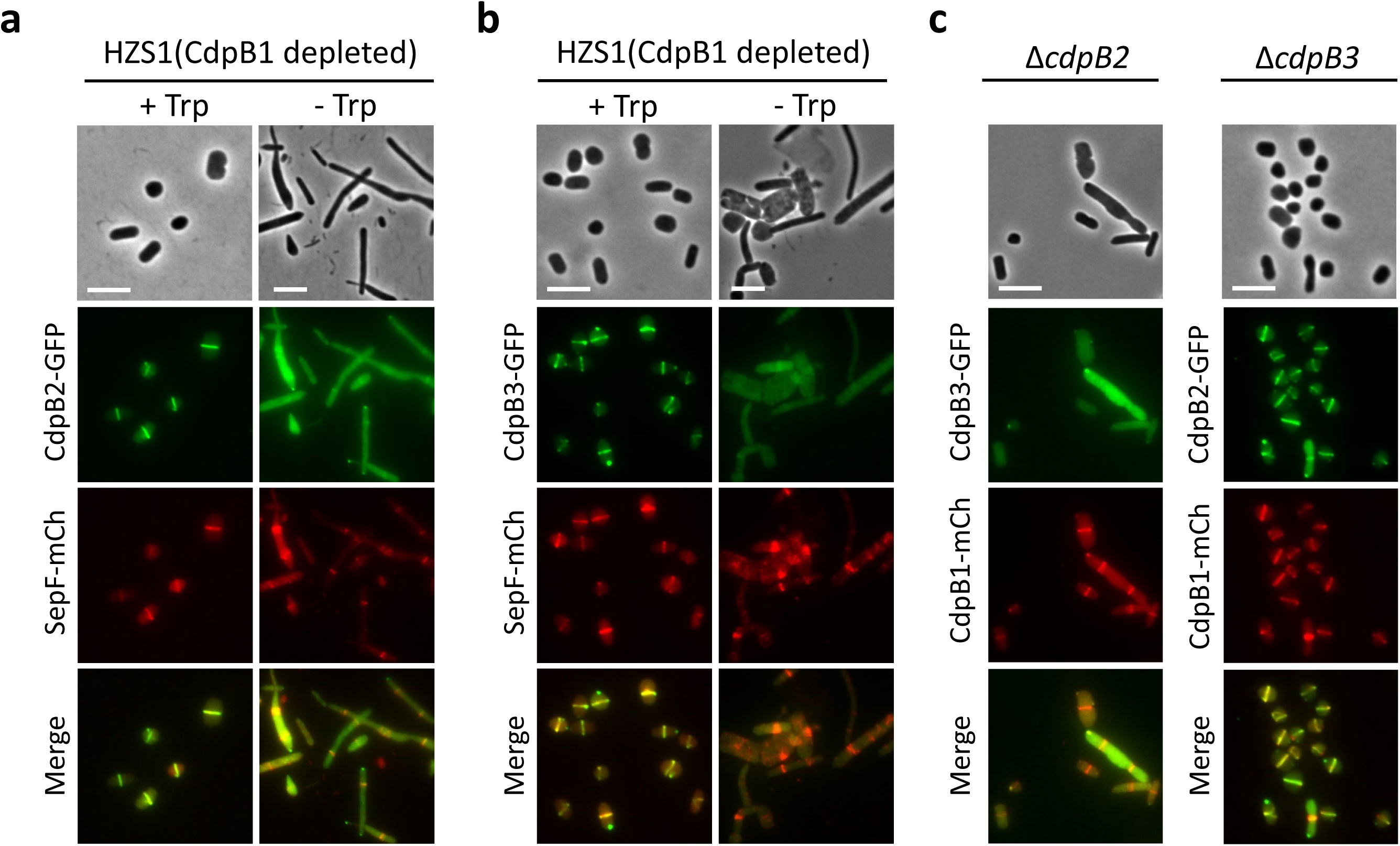
Localization interdependency of the CdpB proteins. **a.** CdpB2 depends on CdpB1 for localization to the division site. Exponential phase culture of HZS1 (H98, *P_tna_::cdpB1*) carrying plasmid pZS408 (*P_phaR_::*c*dpB2-gfp*-*sepF-mCherry*) was treated as in Fig. 1b to check cell morphology and protein localization. **b**. CdpB3 depends on CdpB1 for localization to the division site. Exponential phase culture of HZS1 carrying plasmid pZS407 (*P_phaR_::*c*dpB3-gfp*-*sepF-mCherry*) was treated as in Fig. 1b to check cell morphology and protein localization. **c**. CdpB3 depends on CdpB2 for correction localization, but CdpB1 does not require CdpB2 and CdpB3 for correct localization. Exponential phase cultures of HZS5 (H26, *ΔcdpB2*) carrying plasmid pZS417 (*P_phaR_::*c*dpB3-gfp*-*cdpB1-mCherry*) or HZS6 (H26, *ΔcdpB3*) carrying plasmid pZS418 (*P_phaR_::*c*dpB2-gfp*-*cdpB1-mCherry*) were treated as in Fig. 5d to check the localization of CdpB1 and CdpB3, or CdpB1 and CdpB2, respectively. Scale bar 5 μm.

**Extended Data Fig. 8.**
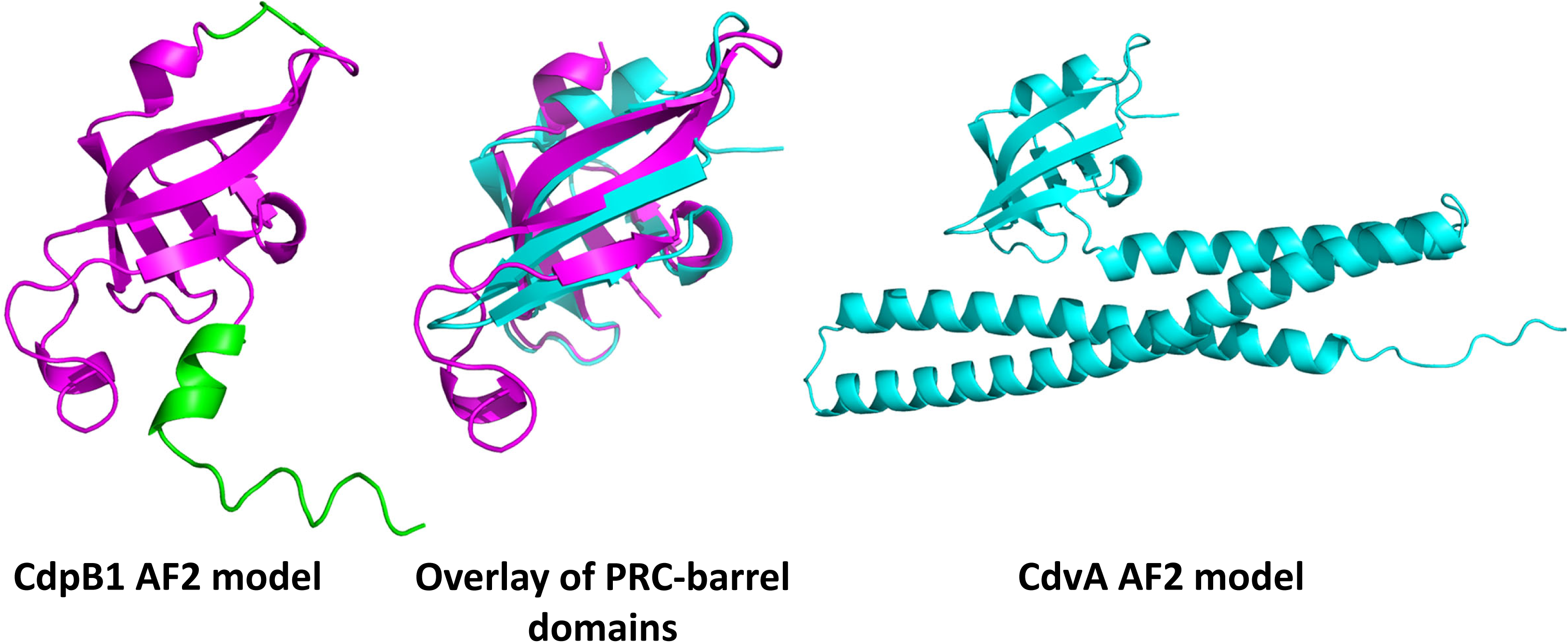
Comparison of CdpB1 with the PRC barrel domain of CdvA from *S. acidocaldricus*. Structural models of CdpB1 (D4H036) and CdvA (Q4J923) were downloaded from Uniprot and aligned with PyMOL.

## Supplementary Information

**Supplementary Fig. 1 A schematic diagram illustrating the procedure for screening for proteins that are toxic when overexpressed.** A genomic library harboring *H. volcanii* DNA segments expressed from a Trp-inducible *P_tna_* promoter was transformed into H26 and transformants were screened on plates with or without tryptophan by replica plating. Transformants displaying a growth defect on plates with tryptophan were inoculated into liquid medium without tryptophan and then re-streaked on plates with or without tryptophan to confirm the tryptophan-dependent growth defect. Sequence analysis was carried out to determine the inserts in the selected transformants, which were then re-cloned into pTA1228 to confirm toxicity, and to detect effects on cell morphology in the presence of tryptophan.

**Supplementary Fig. 2 An example of the tryptophan-dependent growth defect of transformants.** After transformation, the transformants on the original plate containing a proper amount of haloarchaeal colonies were replica plated to plates with and without tryptophan. The colonies indicated by the red arrow grew on the plate without tryptophan, but not on the plate with tryptophan.

**Supplementary Fig. 3 A fragment containing a part of HVO_1691 causes growth inhibition. a.** DNA fragment S-21 contains the first 82 amino acids of SepF, and its expression in the presence of tryptophan retards the growth of *H. volcanii*. **b.** DNA fragment S-295 contains parts of the HVO_1691 (the first 56 amino acids of HVO_1691) and HVO_1692 genes, and its expression in the presence of tryptophan compromises the growth of *H. volcanii*. Single clones of HVO_1691 and HVO_1692 also slow down the growth of *H. volcanii*.

**Supplementary Fig. 4 CdpB1 fluorescent protein fusions localize to the midcell as a ring.** Overnight cultures of *H. volcanii* H26 carrying plasmid pZS103 (*P_tna_::cdpB1-gfp*) or pZS101 (*P_tna_::cdpB1-mCherry*) were diluted 1:100 in fresh Hv.Cab medium with 0.2 mM Trp, and grown at 45°C to OD_600_ about 0.2. 2 μL of the cultures was spotted on a BSW agarose pad for photography. Scale bar 5 μm.

**Supplementary Fig. 5 A schematic diagram for the construction of a CdpB1 depletion strain.** A non-replicating plasmid pZS111 was constructed, and then transformed into *H. volcanii* H98 (DS70, Δ*pyrE2* Δ*hdrB*) after demethylation. Recombination at the *cdpB1* locus by two-step homologous recombination yields the genomic structure shown at the bottom. As a consequence, the transcription of *cdpB1* is now under the control of the specific tryptophan-inducible *P_tna_* promoter in HZS1 (H98, *P_tna_::cdpB1*).

**Supplementary Fig. 6 Growth of the CdpB1 depletion strain in the presence or absence of tryptophan. a.** Depletion of CdpB1 causes a minor growth defect 24 hours post removal of tryptophan. Growth curves (OD_600_) of H26 (black) and HZS1 (red) in Hv.Cab medium supplemented with 1mM tryptophan, while those of H26 (blue) and HZS1 (green) in the medium without tryptophan. **b.** CdpB1 depleted haloarchaeal cells still grow on plates without tryptophan. Spot tests of CdpB1 depleted cells. The mid-log liquid culture with the same optical density was serially diluted 10 times in Hv.Cab medium or the AS-168 medium, then 4 μL of each dilution was spotted on a plate with or without tryptophan. Before taking photos, the plates were cultured at 45°C for 4-5 days.

**Supplementary Fig. 7 Co-IP experiments shows that CdpB1 interacts with SepF.** The experiment was carried out as in Fig. 3c except that anti-GFP primary antibodies were used for immunoprecipitation.

**Supplementary Fig. 8 A schematic diagram for the construction of CdpB1 depletion strain of *Natrinema sp J7* and *H. hispanica*.** Non-replicating plasmids pZS253 (pNBK-F, containing *^NJ7G^cdpB1* upstream flanking sequence followed by the cassette *P_tna_::^NJ7G^cdpB1*) or pZS280 (pHAR, containing *^HAH^cdpB1* upstream flanking sequence followed by the cassette *P_tna_::^HAH^cdpB1*) were transformed into *Natrinema sp.* CJ7-F and *H. hispanica* DF60, respectively, resulting in HZS4 (CJ7-F, *P_tna_::^NJ7G^cdpB1*) and HZS3 (DF60, *P_tna_::^HAH^cdpB1*).

**Supplementary Table 1** Complete list of PRC barrel domain proteins in archaeal genomes from arCOG database.

**Supplementary Table 2** PRC barrel genes neighborhoods.

**Supplementary Table 3** Strains used in this study.

**Supplementary Table 4** Plasmids used in this study.

**Supplementary Table 5** Primers used in this study.

**Supplementary Table 6** Reagents and Chemicals used in this study.

**Supplementary File 1** The complete phylogenetic tree of PRC barrel genes in Newick format.

## References

1 Bi, E. F. & Lutkenhaus, J. FtsZ ring structure associated with division in Escherichia coli. Nature 354, 161–164, doi:10.1038/354161a0 (1991).

2 Du, S. & Lutkenhaus, J. At the Heart of Bacterial Cytokinesis: The Z Ring. Trends Microbiol 27, 781–791, doi:10.1016/j.tim.2019.04.011 (2019).

3 McQuillen, R. & Xiao, J. Insights into the Structure, Function, and Dynamics of the Bacterial Cytokinetic FtsZ-Ring. Annu Rev Biophys 49, 309–341, doi:10.1146/annurev-biophys-121219-081703 (2020).

4 Tsang, M. J. & Bernhardt, T. G. Guiding divisome assembly and controlling its activity. Curr Opin Microbiol 24, 60–65, doi:10.1016/j.mib.2015.01.002 (2015).

5 Haeusser, D. P. & Margolin, W. Splitsville: structural and functional insights into the dynamic bacterial Z ring. Nat Rev Microbiol 14, 305–319, doi:10.1038/nrmicro.2016.26 (2016).

6 Egan, A. J. F., Errington, J. & Vollmer, W. Regulation of peptidoglycan synthesis and remodelling. Nat Rev Microbiol 18, 446–460, doi:10.1038/s41579-020-0366-3 (2020).

7 Mahone, C. R. & Goley, E. D. Bacterial cell division at a glance. J Cell Sci 133, doi:10.1242/jcs.237057 (2020).

8 Wang, X. & Lutkenhaus, J. FtsZ ring: the eubacterial division apparatus conserved in archaebacteria. Mol Microbiol 21, 313–319, doi:10.1046/j.1365-2958.1996.6421360.x (1996).

9 Samson, R. Y., Obita, T., Freund, S. M., Williams, R. L. & Bell, S. D. A role for the ESCRT system in cell division in archaea. Science 322, 1710–1713, doi:10.1126/science.1165322 (2008).

10 Lindas, A. C., Karlsson, E. A., Lindgren, M. T., Ettema, T. J. & Bernander, R. A unique cell division machinery in the Archaea. Proc Natl Acad Sci U S A 105, 18942–18946, doi:10.1073/pnas.0809467105 (2008).

11 Makarova, K. S., Yutin, N., Bell, S. D. & Koonin, E. V. Evolution of diverse cell division and vesicle formation systems in Archaea. Nat Rev Microbiol 8, 731–741, doi:10.1038/nrmicro2406 (2010).

12 Caspi, Y. & Dekker, C. Dividing the Archaeal Way: The Ancient Cdv Cell-Division Machinery. Front Microbiol 9, 174, doi:10.3389/fmicb.2018.00174 (2018).

13 Ithurbide, S., Gribaldo, S., Albers, S. V. & Pende, N. Spotlight on FtsZ-based cell division in Archaea. Trends Microbiol 30, 665–678, doi:10.1016/j.tim.2022.01.005 (2022).

14 van Wolferen, M., Pulschen, A. A., Baum, B., Gribaldo, S. & Albers, S. V. The cell biology of archaea. Nat Microbiol 7, 1744–1755, doi:10.1038/s41564-022-01215-8 (2022).

15 Margolin, W., Wang, R. & Kumar, M. Isolation of an ftsZ homolog from the archaebacterium Halobacterium salinarium: implications for the evolution of FtsZ and tubulin. J Bacteriol 178, 1320–1327, doi:10.1128/jb.178.5.1320-1327.1996 (1996).

16 Baumann, P. & Jackson, S. P. An archaebacterial homologue of the essential eubacterial cell division protein FtsZ. Proc Natl Acad Sci U S A 93, 6726–6730, doi:10.1073/pnas.93.13.6726 (1996).

17 Blanch Jover, A. & Dekker, C. The archaeal Cdv cell division system. Trends Microbiol, doi:10.1016/j.tim.2022.12.006 (2023).

18 Samson, R. Y. et al. Molecular and structural basis of ESCRT-III recruitment to membranes during archaeal cell division. Mol Cell 41, 186–196, doi:10.1016/j.molcel.2010.12.018 (2011).

19 Moriscot, C. et al. Crenarchaeal CdvA forms double-helical filaments containing DNA and interacts with ESCRT-III-like CdvB. PLoS One 6, e21921, doi:10.1371/journal.pone.0021921 (2011).

20 Tarrason Risa, G. et al. The proteasome controls ESCRT-III-mediated cell division in an archaeon. Science 369, doi:10.1126/science.aaz2532 (2020).

21 Dance, A. The mysterious microbes that gave rise to complex life. Nature 593, 328–330, doi:10.1038/d41586-021-01316-0 (2021).

22 Hatano, T. et al. Asgard archaea shed light on the evolutionary origins of the eukaryotic ubiquitin-ESCRT machinery. Nat Commun 13, 3398, doi:10.1038/s41467-022-30656-2 (2022).

23 Hurtig, F. et al. The patterned assembly and stepwise Vps4-mediated disassembly of composite ESCRT-III polymers drives archaeal cell division. Sci Adv 9, eade5224, doi:10.1126/sciadv.ade5224 (2023).

24 Liao, Y., Ithurbide, S., Evenhuis, C., Lowe, J. & Duggin, I. G. Cell division in the archaeon Haloferax volcanii relies on two FtsZ proteins with distinct functions in division ring assembly and constriction. Nat Microbiol 6, 594–605, doi:10.1038/s41564-021-00894-z (2021).

25 Nussbaum, P., Gerstner, M., Dingethal, M., Erb, C. & Albers, S. V. The archaeal protein SepF is essential for cell division in Haloferax volcanii. Nat Commun 12, 3469, doi:10.1038/s41467-021-23686-9 (2021).

26 Pende, N. et al. SepF is the FtsZ anchor in archaea, with features of an ancestral cell division system. Nat Commun 12, 3214, doi:10.1038/s41467-021-23099-8 (2021).

27 Anantharaman, V. & Aravind, L. The PRC-barrel: a widespread, conserved domain shared by photosynthetic reaction center subunits and proteins of RNA metabolism. Genome Biol 3, RESEARCH0061, doi:10.1186/gb-2002-3-11-research0061 (2002).

28 Liao, Y. et al. CdrS Is a Global Transcriptional Regulator Influencing Cell Division in Haloferax volcanii. mBio 12, e0141621, doi:10.1128/mBio.01416-21 (2021).

29 Darnell, C. L. et al. The Ribbon-Helix-Helix Domain Protein CdrS Regulates the Tubulin Homolog ftsZ2 To Control Cell Division in Archaea. mBio 11, doi:10.1128/mBio.01007-20 (2020).

30 Allers, T., Ngo, H. P., Mevarech, M. & Lloyd, R. G. Development of additional selectable markers for the halophilic archaeon Haloferax volcanii based on the leuB and trpA genes. Appl Environ Microbiol 70, 943–953, doi:10.1128/AEM.70.2.943-953.2004 (2004).

31 Large, A. et al. Characterization of a tightly controlled promoter of the halophilic archaeon Haloferax volcanii and its use in the analysis of the essential cct1 gene. Mol Microbiol 66, 1092–1106, doi:10.1111/j.1365-2958.2007.05980.x (2007).

32 Hu, C. D. & Kerppola, T. K. Simultaneous visualization of multiple protein interactions in living cells using multicolor fluorescence complementation analysis. Nat Biotechnol 21, 539–545, doi:10.1038/nbt816 (2003).

33 Cabantous, S. et al. A new protein-protein interaction sensor based on tripartite split-GFP association. Sci Rep 3, 2854, doi:10.1038/srep02854 (2013).

34 Malakhov, M. P. et al. SUMO fusions and SUMO-specific protease for efficient expression and purification of proteins. J Struct Funct Genomics 5, 75–86, doi:10.1023/B:JSFG.0000029237.70316.52 (2004).

35 Tehrani, A., Prince, R. C. & Beatty, J. T. Effects of photosynthetic reaction center H protein domain mutations on photosynthetic properties and reaction center assembly in Rhodobacter sphaeroides. Biochemistry 42, 8919–8928, doi:10.1021/bi0346650 (2003).

36 Lovgren, J. M. et al. The PRC-barrel domain of the ribosome maturation protein RimM mediates binding to ribosomal protein S19 in the 30S ribosomal subunits. RNA 10, 1798–1812, doi:10.1261/rna.7720204 (2004).

37 Abecasis, A. B. et al. A genomic signature and the identification of new sporulation genes. J Bacteriol 195, 2101–2115, doi:10.1128/JB.02110-12 (2013).

38 Weiss, C. A. et al. NrnA is a 5’-3’ exonuclease that processes short RNA substrates in vivo and in vitro. Nucleic Acids Res 50, 12369–12388, doi:10.1093/nar/gkac1091 (2022).

39 Chen, A. W. et al. The Role of 3’ to 5’ Reverse RNA Polymerization in tRNA Fidelity and Repair. Genes (Basel) 10, doi:10.3390/genes10030250 (2019).

40 Moir, D., Stewart, S. E., Osmond, B. C. & Botstein, D. Cold-sensitive cell-division-cycle mutants of yeast: isolation, properties, and pseudoreversion studies. Genetics 100, 547–563, doi:10.1093/genetics/100.4.547 (1982).

41 Dassain, M., Leroy, A., Colosetti, L., Carole, S. & Bouche, J. P. A new essential gene of the ‘minimal genome’ affecting cell division. Biochimie 81, 889–895, doi:10.1016/s0300-9084(99)00207-2 (1999).

42 Maupin-Furlow, J. Proteasomes and protein conjugation across domains of life. Nat Rev Microbiol 10, 100–111, doi:10.1038/nrmicro2696 (2011).

43 Duggin, I. G. et al. CetZ tubulin-like proteins control archaeal cell shape. Nature 519, 362–365, doi:10.1038/nature13983 (2015).

44 de Silva, R. T. et al. Improved growth and morphological plasticity of Haloferax volcanii. Microbiology (Reading) 167, doi:10.1099/mic.0.001012 (2021).

45 Allers, T., Barak, S., Liddell, S., Wardell, K. & Mevarech, M. Improved strains and plasmid vectors for conditional overexpression of His-tagged proteins in Haloferax volcanii. Appl Environ Microbiol 76, 1759–1769, doi:10.1128/AEM.02670-09 (2010).

46 Cai, S. et al. Identification of the haloarchaeal phasin (PhaP) that functions in polyhydroxyalkanoate accumulation and granule formation in Haloferax mediterranei. Appl Environ Microbiol 78, 1946–1952, doi:10.1128/AEM.07114-11 (2012).

47 Ye, X., Ou, J., Ni, L., Shi, W. & Shen, P. Characterization of a novel plasmid from extremely halophilic Archaea: nucleotide sequence and function analysis. FEMS Microbiol Lett 221, 53–57, doi:10.1016/S0378-1097(03)00175-7 (2003).

48 Liu, H., Han, J., Liu, X., Zhou, J. & Xiang, H. Development of pyrF-based gene knockout systems for genome-wide manipulation of the archaea Haloferax mediterranei and Haloarcula hispanica. J Genet Genomics 38, 261–269, doi:10.1016/j.jgg.2011.05.003 (2011).

49 Wang, J. et al. A novel family of tyrosine integrases encoded by the temperate pleolipovirus SNJ2. Nucleic Acids Res 46, 2521–2536, doi:10.1093/nar/gky005 (2018).

50 de Boer, P. A., Crossley, R. E. & Rothfield, L. I. A division inhibitor and a topological specificity factor coded for by the minicell locus determine proper placement of the division septum in E. coli. Cell 56, 641–649, doi:10.1016/0092-8674(89)90586-2 (1989).

51 Bernhardt, T. G. & de Boer, P. A. SlmA, a nucleoid-associated, FtsZ binding protein required for blocking septal ring assembly over Chromosomes in E. coli. Mol Cell 18, 555–564, doi:10.1016/j.molcel.2005.04.012 (2005).

52 Hale, C. A. & de Boer, P. A. Direct binding of FtsZ to ZipA, an essential component of the septal ring structure that mediates cell division in E. coli. Cell 88, 175–185, doi:10.1016/s0092-8674(00)81838-3 (1997).

53 Pichoff, S. & Lutkenhaus, J. Identification of a region of FtsA required for interaction with FtsZ. Mol Microbiol 64, 1129–1138, doi:10.1111/j.1365-2958.2007.05735.x (2007).

54 Du, S. & Lutkenhaus, J. SlmA antagonism of FtsZ assembly employs a two-pronged mechanism like MinCD. PLoS Genet 10, e1004460, doi:10.1371/journal.pgen.1004460 (2014).

55 Makarova, K. S., Wolf, Y. I. & Koonin, E. V. Archaeal Clusters of Orthologous Genes (arCOGs): An Update and Application for Analysis of Shared Features between Thermococcales, Methanococcales, and Methanobacteriales. Life (Basel) 5, 818–840, doi:10.3390/life5010818 (2015).

56 Makarova, K. S., Wolf, Y. I. & Koonin, E. V. Towards functional characterization of archaeal genomic dark matter. Biochem Soc Trans 47, 389–398, doi:10.1042/BST20180560 (2019).

57 Altschul, S. F. et al. Gapped BLAST and PSI-BLAST: a new generation of protein database search programs. Nucleic Acids Res 25, 3389–3402, doi:10.1093/nar/25.17.3389 (1997).

58 Zimmermann, L. et al. A Completely Reimplemented MPI Bioinformatics Toolkit with a New HHpred Server at its Core. J Mol Biol 430, 2237–2243, doi:10.1016/j.jmb.2017.12.007 (2018).

59 Edgar, R. C. Muscle5: High-accuracy alignment ensembles enable unbiased assessments of sequence homology and phylogeny. Nat Commun 13, 6968, doi:10.1038/s41467-022-34630-w (2022).

60 Esterman, E. S., Wolf, Y. I., Kogay, R., Koonin, E. V. & Zhaxybayeva, O. Evolution of DNA packaging in gene transfer agents. Virus Evol 7, veab015, doi:10.1093/ve/veab015 (2021).

61 Price, M. N., Dehal, P. S. & Arkin, A. P. FastTree 2--approximately maximum-likelihood trees for large alignments. PLoS One 5, e9490, doi:10.1371/journal.pone.0009490 (2010).

62 Delorenzi, M. & Speed, T. An HMM model for coiled-coil domains and a comparison with PSSM-based predictions. Bioinformatics 18, 617–625, doi:10.1093/bioinformatics/18.4.617 (2002).

