## Supplemental Information for "Widespread PRC barrel proteins play critical roles in archaeal cell division"

**Affiliation**

* To whom correspondence should be addressed:

Shishen Du

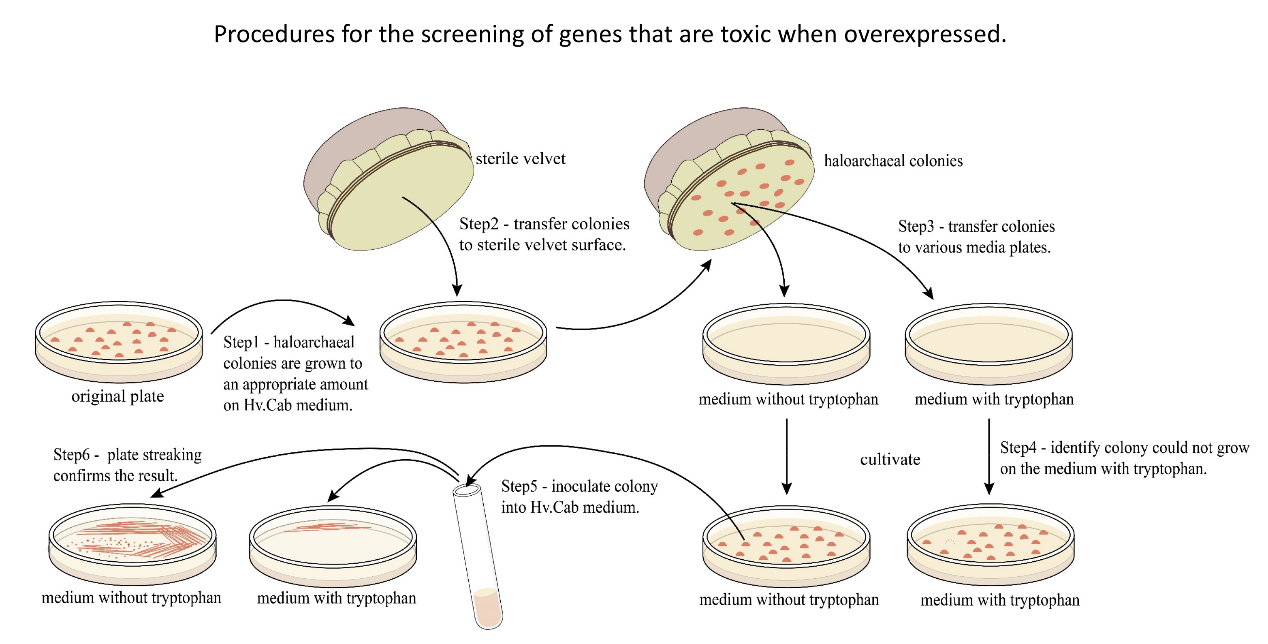

**Supplementary Figure S1. A schematic diagram illustrating the procedure for screening for proteins that are toxic when overexpressed.** A genomic library harboring *H. volcanii* DNA segments expressed from a Trp-inducible *P_tna_* promoter was transformed into H26 and transformants were screened on plates with or without tryptophan by replica plating. Transformants displaying a growth defect on plates with tryptophan were inoculated into liquid medium without tryptophan and then re-streaked on plates with or without tryptophan to confirm the tryptophan-dependent growth defect. Sequence analysis was carried out to determine the inserts in the selected transformants, which were then re-cloned into pTA1228 to confirm toxicity, and to detect effects on cell morphology in the presence of tryptophan.

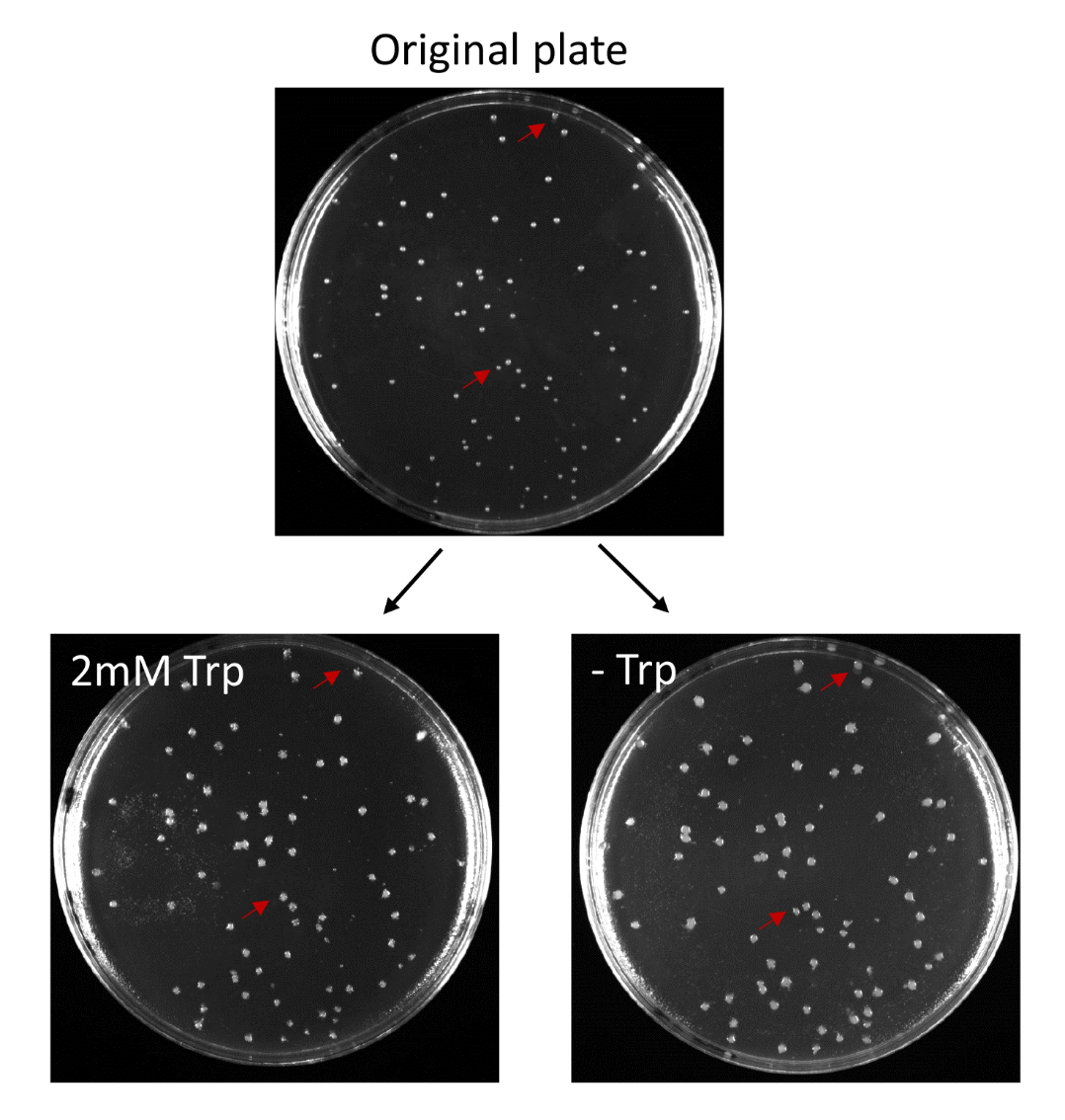

**Supplementary Figure S2. An example of the tryptophan-dependent growth defect of transformants.** After transformation, the transformants on the original plate containing a proper amount of haloarchaeal colonies were replica plated to plates with and without tryptophan. The colonies indicated by the red arrow grew on the plate without tryptophan, but not on the plate with tryptophan.

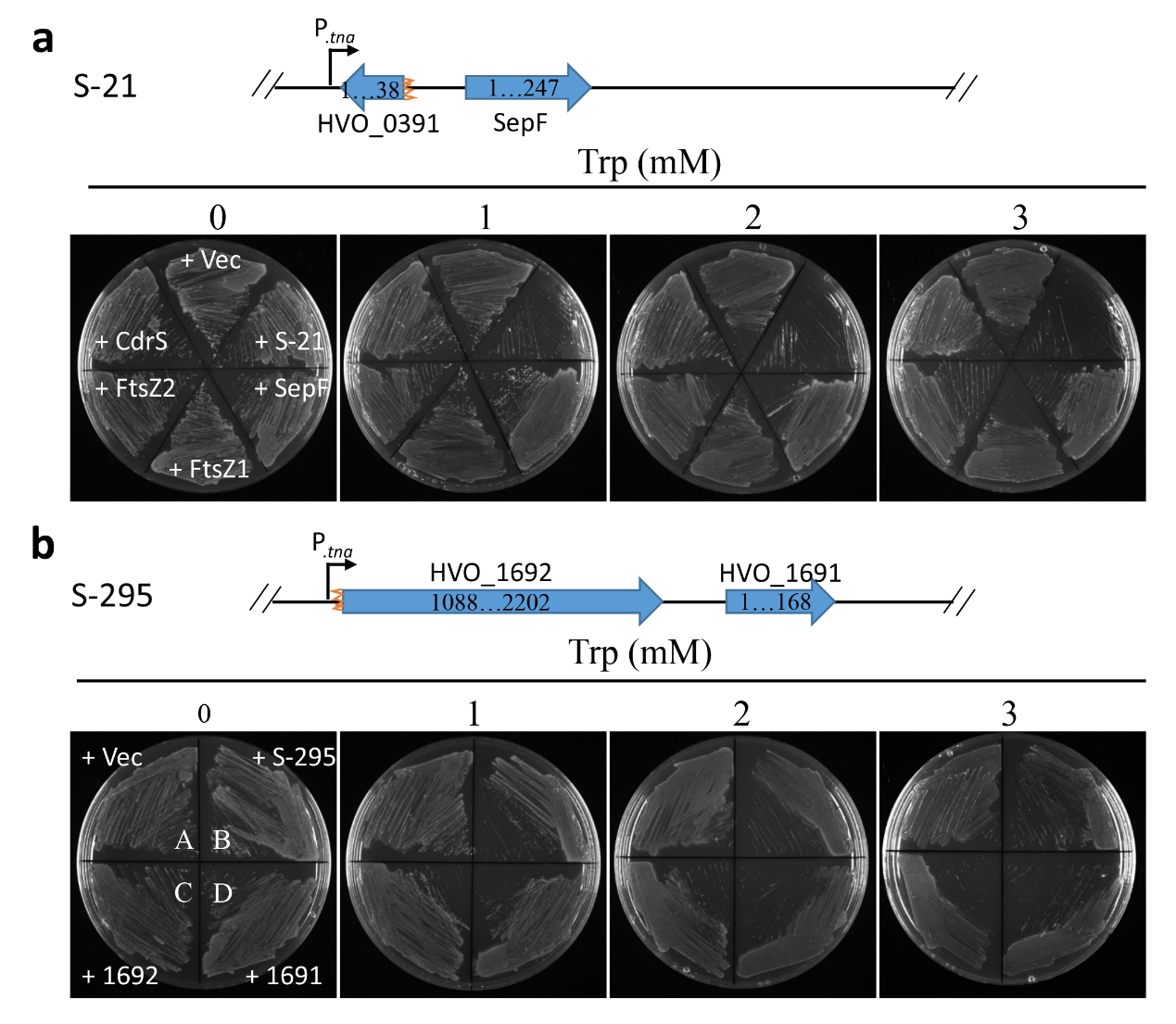

**Supplementary Figure S3. A fragment containing a part of HVO_1691 causes growth inhibition.** **a.** DNA fragment S-21 contains the first 82 amino acids of SepF, and its expression in the presence of tryptophan retards the growth of *H. volcanii*. **b.** DNA fragment S-295 contains parts of the HVO_1691 (the first 56 amino acids of HVO_1691) and HVO_1692 genes, and its expression in the presence of tryptophan compromises the growth of *H. volcanii*. Single clones of HVO_1691 and HVO_1692 also slow down the growth of *H. volcanii*.

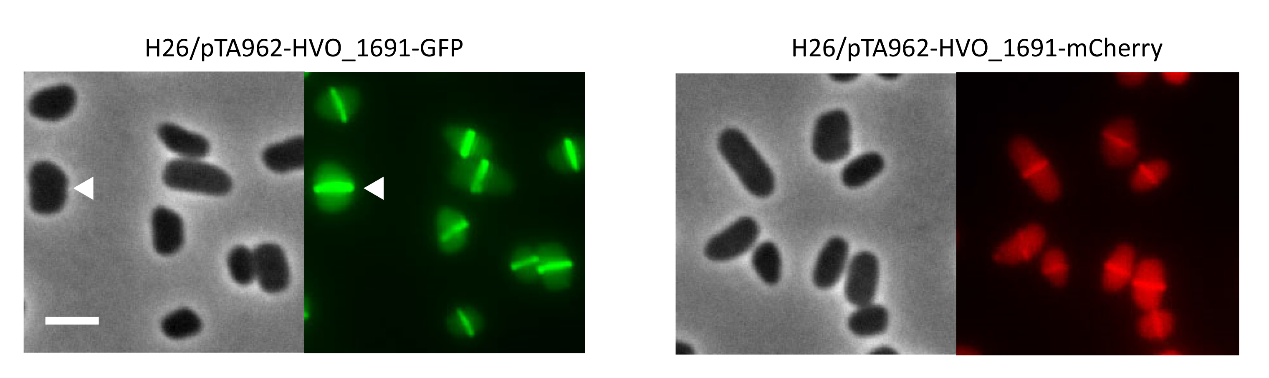

**Supplementary Figure S4. CdpB1 fluorescent protein fusions localize to the midcell as a ring.** Overnight cultures of *H. volcanii* H26 carrying plasmid pZS103 (*P_tna_::cdpB1-gfp*) or pZS101 (*P_tna_::cdpB1-mCherry*) were diluted 1:100 in fresh Hv.Cab medium with 0.2 mM Trp, and grown at 45˚C to OD_600_ about 0.2. 2 μL of the cultures was spotted on a BSW agarose pad for photography. Scale bar 5 μm.

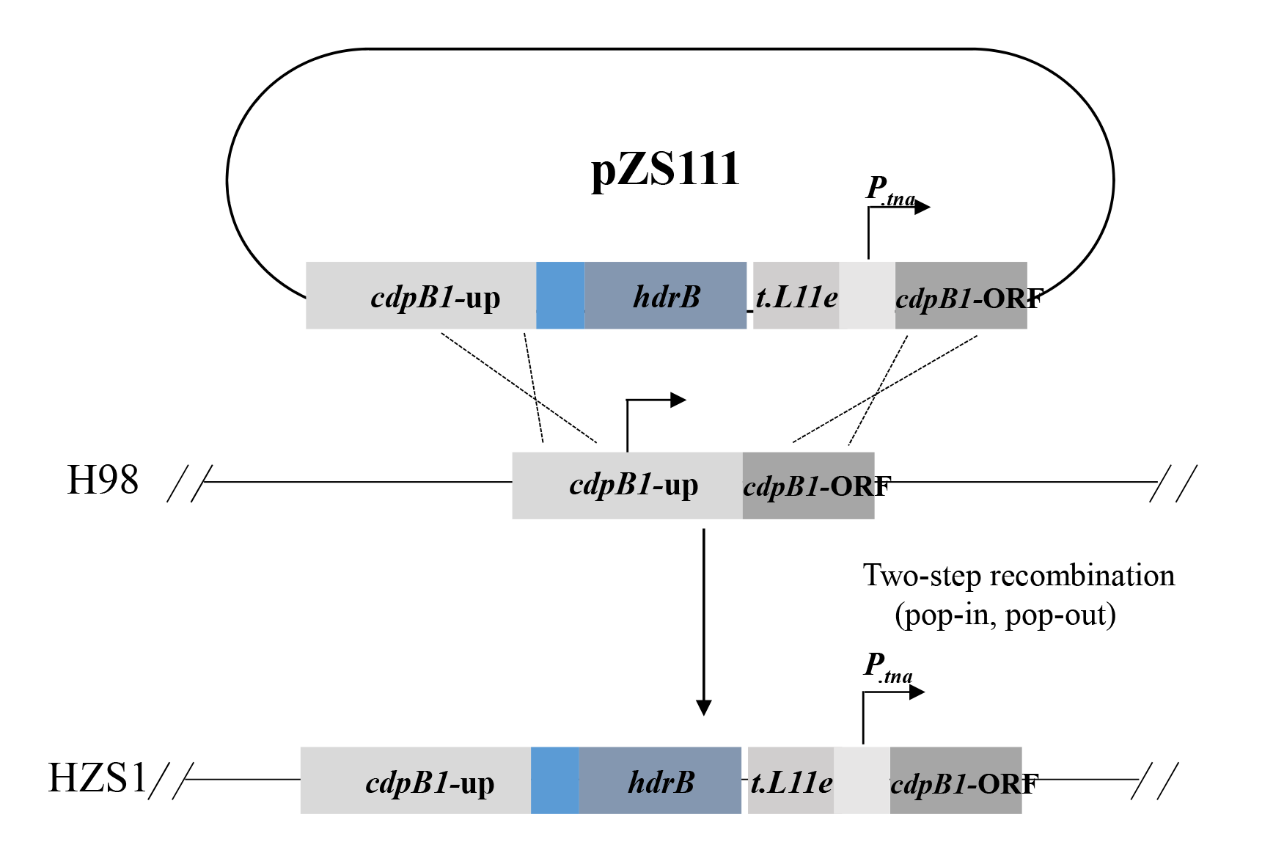

**Supplementary Figure S5. A schematic diagram for the construction of a CdpB1 depletion strain.** A non-replicating plasmid pZS111 was constructed, and then transformed into *H. volcanii* H98 (DS70, Δ*pyrE2* Δ*hdrB*) after demethylation. Recombination at the *cdpB1* locus by two-step homologous recombination yields the genomic structure shown at the bottom. As a consequence, the transcription of *cdpB1* is now under the control of the specific tryptophan-inducible *P_tna_* promoter in HZS1 (H98, *P_tna_::cdpB1*).

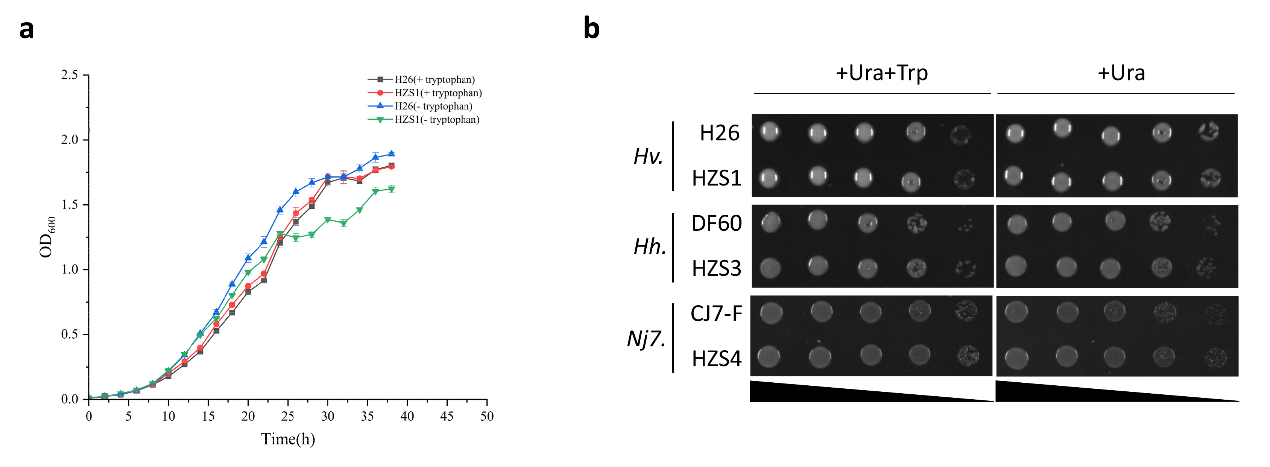

**Supplementary Figure S6. Growth of the CdpB1 depletion strain in the presence or absence of tryptophan.** **a.** Depletion of CdpB1 causes a minor growth defect 24 hours post removal of tryptophan. Growth curves (OD_600_) of H26 (black) and HZS1 (red) in Hv.Cab medium supplemented with 1mM tryptophan, while those of H26 (blue) and HZS1 (green) in the medium without tryptophan. **b.** CdpB1 depleted haloarchaeal cells still grow on plates without tryptophan. Spot tests of CdpB1 depleted cells. The mid-log liquid culture with the same optical density was serially diluted 10 times in Hv.Cab medium, then 4 μL of each dilution was spotted on a plate with or without tryptophan. Before taking photos, the plates were cultured at 45°C for 4-5 days.

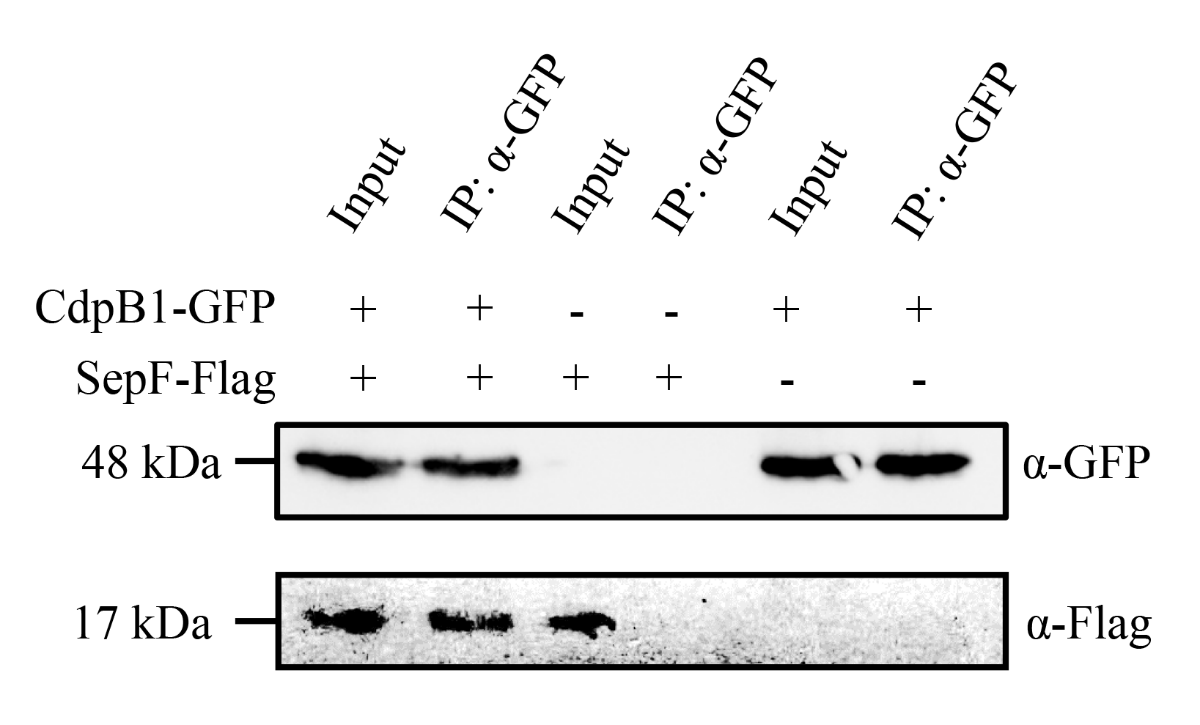

**Supplementary Figure S7. Co-IP experiments shows that CdpB1 interacts with SepF.** The experiment was carried out as in Fig. 3c except that anti-GFP primary antibodies were used for immunoprecipitation.

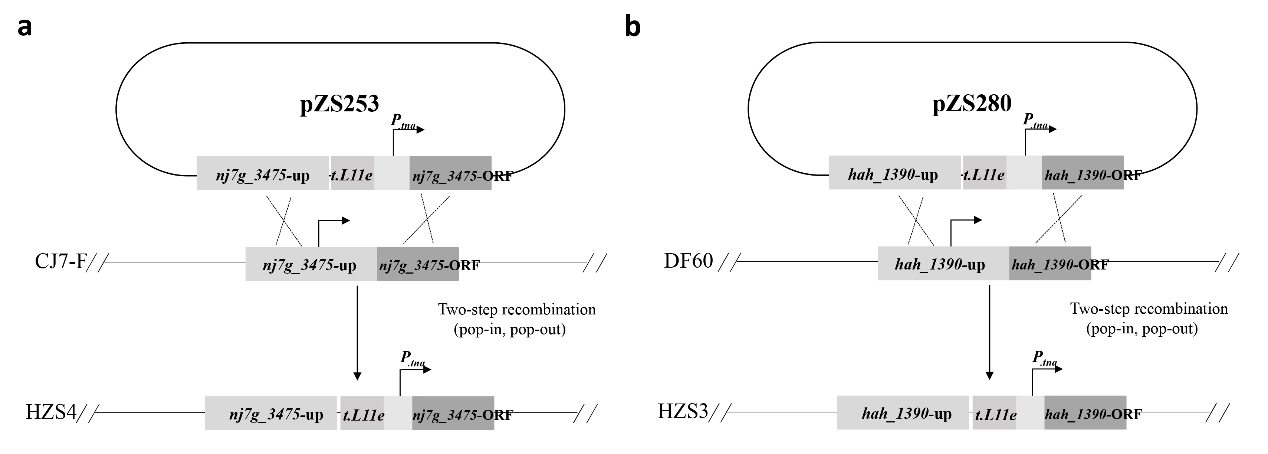

**Supplementary Figure S8. A schematic diagram for the construction of CdpB1 depletion strain of *Natrinema sp J7* and *H. hispanica*.** Non-replicating plasmids pZS253 (pNBK-F, containing *^NJ7G^cdpB1* upstream flanking sequence followed by the cassette *P_tna_::^NJ7G^cdpB1*) or pZS280 (pHAR, containing *^HAH^cdpB1* upstream flanking sequence followed by the cassette *P_tna_::^HAH^cdpB1*) were transformed into *Natrinema sp.* CJ7-F and *H. hispanica* DF60, respectively, resulting in HZS4 (CJ7-F, *P_tna_::^NJ7G^cdpB1*) and HZS3 (DF60, *P_tna_::^HAH^cdpB1*).

| **Table S1. Strains used in this study.** | |  |  |
| --- | --- | --- | --- |
| **Strain** | **Genotype** | **Description** | **Source** |
| ***E. coli*** | | | |
| JS238 | MC1061 *malPp::lacIQ srlC::Tn10 recA1* | General cloning strain for plasmid construction | (1) |
| JM110 | *dam dcm supE*44 *hsdR17 thi leu rpsL lacY galK galT ara tonA thr tsx D(lac-proAB) F'(traD36 proAB+lacIq lacZ* △*M15)* | DNA methylation-deficient strain for preparation of demethylated plasmids for *H. volcanii* transformation | CCTCC |
| BL21(DE3) | *F – ompT hsdSB (rB- mB-) gal dcm (DE3)* | host for protein production | (2) |
| ***Halophilic archaeon*** | | | |
| DS70 | (DS2) ΔpHV2 | Wild-type *H. volcanii* DS2 cured of pHV2 | (3) |
| H26 | (DS70) Δ*pyrE2* | Auxotroph (uracil) | (4) |
| H98 | (DS70) Δ*pyrE2* Δ*hdrB* | Auxotroph (uracil, hypoxanthine and thymidine) | (4) |
| ID56 | (H98) *P_fdx_::hdrB P_tna_*::*ftsZ1* | Trp-regulated expression of genomic *ftsZ1* (HVO_0717) | (5) |
| ID57 | (H98) *P_fdx_::hdrB P_tna_::ftsZ2* | Trp-regulated expression of genomic *ftsZ2* (HVO_0581) | (5) |
| ID112 | Δ*ftsZ2* and Δ*ftsZ1* (Double deletion) | Δ*ftsZ2* and Δ*ftsZ1* (Double deletion) | (5) |
| HZS1 | (H98) *P_fdx_::hdrB P_tna_::cdpB1* | Trp-regulated expression of genomic *cdpB1* (HVO_1691) | This study |
| HZS2 | (H98) *P_fdx_::hdrB P_tna_::sepF* | Trp-regulated expression of genomic *sepF* (HVO_0392) | This study |
| HZS3 | (DF60) *P_tna_::^HAH^cdpB1* | Trp-regulated expression of genomic *hah_1390* | This study |
| HZS4 | (CJ7-F) P_tna_:: *^NJ7G^cdpB1* | Trp-regulated expression of genomic *nj7g_3475* | This study |
| HZS5 | (H26) Δ*cdpB2* | Deletion of *cdpB2* (*HVO_1964*) | This study |
| HZS6 | (H26) Δc*dpB3* | Deletion of *cdpB3* (*HVO_2019*) | This study |
| *H. hispanica* DF60 | △*pyrF* | Auxotroph (uracil) | (6) |
| *Natrinema* sp. CJ7-F | △*pyrF* | Auxotroph (uracil) | (7) |

| **Table S2. Plasmids used in this study.** | | | |
| --- | --- | --- | --- |
| **Plasmid** | **Description/function** | **Primer name** | |
| **Plasmids for gene expression** | | | |
| pTA1228 | *P_tna_*::6XHis expression vector for *H.volcanii* |  | (8) |
| pIDJL40 | pTA962 with *gfp* (BamHI-NotI) |  | (9) |
| pIDJL40-*ftsZ1* | *P_tna_* control of *ftsZ1*-*gfp* |  | (9) |
| pIDJL114 | *P_tna_* control of *ftsZ1-mCherry* |  | (5) |
| pIDJL115 | *P_tna_* control of *ftsZ2-mCherry* |  | (5) |
| pWL502 | carrying *pyrF* and its native promoter for *H.hispanica* DF60 |  | (10) |
| pFJ6-*P_phaR_*-GFP | *pFJ6-MCS with AflII-SphI fragment containing promoter of the P_phaR_ operon from Haloferax mediterranei and GFP* |  | (11) |
| pFJ6-*P_tna_* | *P_tna_* expression vector for *Natrinema* sp. CJ7-F |  | Lab collection |
| pZWC176 | *sepF-flag* under control of the native promoter of sepF | P_native_-sepF-F, P_native_-sepF-R, sepF-F7, sepF-flag-R1, sepF-flag-R2 | This study |
| pZWC185 | *sepF-flag* and *cdpB1-gfp* under control of the native promoter of *sepF* and *cdpB1* | P_native_-cdpB1-F, gfp-R | This study |
| pZS1 | pTA1228 with a ClaI restriction site for construction of genomic library | Clal-EcoRI-F, Clal-BamHI-R | This study |
| pZS101 | *P_tna_* control of *cdpB1-mCherry* | cdpB1-NdeI-F, cdpB1-BamHI-R2 | This study |
| pZS103 | *P_tna_* control of *cdpB1-gfp* | cdpB1-NdeI-F, cdpB1-BamHI-R2 | This study |
| pZS104 | *P_tna_* control of *sepF-gfp* | sepF-NdeI-F, sepF-BamHI-R1 | This study |
| pZS105 | *P_tna_* control of *cdpB1-gfp* and *ftsZ1-mCherry* | ftsZ1-Notl-U-F, mCH-Notl-U-R | This study |
| pZS106 | *P_tna_* control of *cdpB1-gfp* and *ftsZ2-mCherry* | ftsZ2-Notl-U-F, mCH-Notl-U-R | This study |
| pZS107 | *P_tna_* control of *cdpB1-gfp* and *sepF-mCherry* | sepf-Notl-U-F, sepF-BamHI-U-R, mCH-U-F2, mCH-Notl-U-R | This study |
| pZS124 | *P_tna_* control of *ftsZ1* | ftsZ1-NdeI-F, ftsZ1-BamHI-R2 | This study |
| pZS125 | *P_tna_* control of *ftsZ2* | ftsZ2-NdeI-F, ftsZ2-BamHI-R2 | This study |
| pZS126 | *P_tna_* control of *cdpB1* | cdpB1-NdeI-F, cdpB1-BamHI-R1 | This study |
| pZS127 | *P_tna_* control of *sepF* | sepF-NdeI-F, sepF-BamHI-R2 | This study |
| pZS204 | *P_tna_* control of *cdrS* | cdrS -NdeI-F, cdrS-BamHI-R | This study |
| pZS208 | *ftsZ1-gfp* under the control of the native promoter of *ftsZ1* | P_native_-ftsZ1-ApaI-F, ftZ1-BamHI-R1. | This study |
| pZS214 | *cdpB1-gfp* under the control of the native promoter of *cdpB1* | P_native_-cdpB1-ApaI-F, cdpB1-BamHI-R2 | This study |
| pZS217 | *P_tna_* control of *^NJ7G^cdpB1-gfp* | P.tna (J7)-AflII-U-F, P.tna (J7)-U-R, ^NJ7G^cdpB1-U-F, ^NJ7G^cdpB1-U-R, gfp-U-F1, gfp-SphI-U-R | This study |
| pZS234 | *^NJ7G^cdpB1* under the control of its native promoter of | P_native_- ^NJ7G^cdpB1-ApaI-F, P_native_-^NJ7G^cdpB1-BamHI-R | This study |
| pZS236 | *cdpB1* under the control of its native promoter | P_native_- cdpB1-ApaI-F, cdpB1-BamHI-R1 | This study |
| pZS239 | *P_phaR_* control of *cdpB1-gfp* and *ftsZ1-mCherry* | p.phaRP-ApaI-F, p.phaRP-NdeI-R | This study |
| pZS276 | *^HAH^cdpB1* under the control of its native promoter | P_native_-^HAH^cdpB1-ApaI-F, P_native_-^HAH^cdpB1-BamHI-R | This study |
| pZS284 | *P_phaR_* control of *cdpB1-gfp* and *ftsZ2-mCherry* | p.phaRP-U-F, p.phaRP-U-R | This study |
| pZS285 | *P_phaR_* control of *cdpB1-gfp* and *sepF-mCherry* | p.phaRP-U-F, p.phaRP-U-R | This study |
| pZS289 | *P_phaR_* control of *ftsZ1-gfp* and *ftsZ2-mCherry* | ftsZ1-NdeI-U-F, gfp-U-R1, ftsZ2-U-F, mCH-NotI-U-R | This study |
| pZS306 | *P_tna_* control of *^HAH^cdpB1-gfp* | P.tna-EcoRI-U-F, P.tna-U-R, ^HAH^cdpB1-U-F, ^HAH^cdpB1-U-R, gfp-U-F2, gfp-ClaI-U-R | This study |
| pZS322 | *P_phaR_* control of *sepF-gfp* and *ftsZ1-mCherry* | sepF-NdeI-U-F, gfp-U-R2, ftsZ1-U-F, mCH-NotI-U-R | This study |
| pZS324 | *P_phaR_* control of *sepF-gfp* and *ftsZ2-mCherry* | ftsZ2-Notl-U-F, mCH-Notl-U-R | This study |
| pZS336 | *P_tna_* control of *cdpB2-gfp* | cdpB2-NdeI-F, cdpB2-BamHI-R | This study |
| pZS337 | *P_tna_* control of *cdpB3-gfp* | cdpB3-NdeI-F, cdpB3-BamHI-R | This study |
| pZS338 | *cdpB3* under the control of its native promoter | P_native_-cdpB3-ApaI-F, cdpB3-BamHI-R1 | This study |
| pZS339 | *cdpB2* under the control of its native promoter | P_native_-cdpB2-ApaI-F, cdpB2-BamHI-R1 | This study |
| pZS384 | *P_tna_* control of *NJ7G_2729-gfp* | P.tna (J7)-AflII-U-F, P.tna (J7)-U-R1, ^NJ7G^cdpB2-U-F, ^NJ7G^cdpB2-U-R, gfp-U-F3, gfp-SphI-U-R | This study |
| pZS385 | *P_tna_* control of *NJ7G_3497-gfp* | P.tna (J7)-AflII-U-F, P.tna (J7)-U-R2, ^NJ7G^cdpB3-U-F, ^NJ7G^cdpB3-U-R, gfp-U-F4, gfp-SphI-U-R | This study |
| pZS388 | *P_tna_* control of *HAH_0460-gfp* | ^HAH^cdpB2-U-F, ^HAH^cdpB2-U-R, gfp-U-F5, gfp-ClaI-U-R | This study |
| pZS389 | *P_tna_* control of *HAH_5240-gfp* | ^HAH^cdpB3-U-F, ^HAH^cdpB3-U-R, gfp-U-F6, gfp-ClaI-U-R | This study |
| pZS390 | *cdpB2 and cdpB3* under the control of their native promoters | P_native_-cdpB2-BamHI-F, cdpB2-NotI-R | This study |
| pZS407 | *P_phaR_* control of *cdpB3-gfp* and *sepF-mCherry* | cdpB3-NdeI-U-F, gfp-U-R3, sepF-Notl-U-F, mCH-NotI-U-R | This study |
| pZS408 | *P_phaR_* control of *cdpB2-gfp* and *sepF-mCherry* | cdpB2-NdeI-U-F, gfp-U-R3, sepF-Notl-U-F, mCH-NotI-U-R | This study |
| pZS417 | *P_phaR_* control of *cdpB3-gfp* and *cdpB1-mCherry* | mCH-NotI-U-R, cdpB1-NotI-U-F | This study |
| pZS418 | *P_phaR_* control of *cdpB2-gfp* and *cdpB1-mCherry* | mCH-NotI-U-R, cdpB1-NotI-U-F | This study |
| pZS422 | *P_tna_* control of *hvo_1607-gfp* | hvo_1607-NdeI-F, hvo_1607-BamHI-R | This study |
| pZS423 | *P_tna_* control of *gfp- hvo_1607* | hvo_1607-NotI-F, hvo_1607-NotI-R | This study |
| **Plasmids for genomic modification** | | | |
| pTA131 | Cloning vector with *pyrE2* marker, for *H.volcanii* genetic modification |  | (4) |
| pNBK-F | Cloning vector with *pyrF* marker, for *Natrinema sp* genetic modification |  | (7) |
| pHAR | Cloning vector with *pyrF* marker, for *Haloarcula hispanica*genetic modification |  | (6) |
| pZS98 | pTA131 with *cdpB1*-upstream flanking sequence and the *P_tna_::cdpB1* cassette | cdpB1UP-Hindlll-U-F, cdpB1UP-U-R, P.tna-cdpB1-U-F, cdpB1-BamHI-U-R | This study |
| pZS99 | pTA131 with *sepF*-upstream flanking sequence and the *P_tna_::SepF* cassette | sepFUP-Hindlll-U-F, sepFUP-U-R, P.tna-sepF-U-F, sepF-BamHI-U-R | This study |
| pZS100 | pTA131 with *cdpB1*-upstream and downstream flanking sequences | cdpB1UP-Hindlll-U-F, cdpB1UP-U-R, cdpB1down-U-F, cdpB1down-BamHI-U-R | This study |
| pZS111 | pZS98 with *P_fdx_::hdrB* inserted at the sphI site (upstream of *P_tna_*). For replacement of *cdpB1* promoter | p.fdx-hdrB-F, p.fdx-hdrB-R | This study |
| pZS219 | pZS99 with *P_fdx_::hdrB* inserted at the sphI site (upstream of *P_tna_*). For replacement of *sepF* promoter | p.fdx-hdrB-F, p.fdx-hdrB-R | This study |
| pZS253 | pNBK-F with sp *^NJ7G^cdpB1*-upstream flanking sequence and the *P_tna_:: ^NJ7G^cdpB1* cassette. For replacement of *^NJ7G^cdpB1* promoter | ^NJ7G^cdpB1UP-Hindlll-U-F, ^NJ7G^cdpB1UP-U-R, P.tna-^NJ7G^cdpB1-U-F, ^NJ7G^cdpB1-AflII-U-R | This study |
| pZS280 | pHAR with *^HAH^cdpB1*-upstream flanking sequence and the *P_tna_::^HAH^cdpB1* cassette. For replacement of *^HAH^cdpB1* promoter | ^HAH^cdpB1UP- KpnI-U-F, ^HAH^cdpB1UP-U-R, P.tna-^HAH^cdpB1-U-F, ^HAH^cdpB1-HindIII-U-R | This study |
| pZS398 | pTA131 with *cdpB2*-upstream downstream flanking sequence. For *cdpB2* deletion. | cdpB2UP-Hindlll-U-F, cdpB2UP-U-R, cdpB2down-U-F, cdpB2down-BamHI-U-R | This study |
| pZS399 | pTA131 with *cdpB3*-upstream and downstream flanking sequence. For *cdpB3* deletion. | cdpB3UP-Hindlll-U-F, cdpB3UP-U-R, cdpB3down-U-F, cdpB3down-BamHI-U-R | This study |
| **Plasmids for protein purification** | | | |
| pE-SUMO | P_T7_::*his-SUMO*, Amp |  | Lab collection |
| pZS311 | pE-SUMO, P_T7_::*his-SUMO-CdpB1*, Amp | SUMO-SepF-BsaI-F, SepF-XbaI-R | This study |
| pZS288 | pE-SUMO, P_T7_::*his-SUMO-SepF*, Amp | SUMO-CdpB1-BsaI-F, CdpB1-XbaI-R | This study |
| **Plasmids for Split-FP assay** | | | |
| psfGFP-1-9 | carrying the N-terminal fragment GFP1-9 (residues 1-193) of the *sfGFP* |  | (12) |
| psfGFP-10-I | carrying the β-hairpin GFP10 (residues 194-212) of the *sfGFP* with a C-terminal long linker |  | (12) |
| psfGFP-10-II | carrying the β-hairpin GFP10 (residues 194-212) of the *sfGFP* with a C-terminal short linker |  | (12) |
| psfGFP-11-I | carrying the β-hairpin GFP11 (residues 213-233) of the *sfGFP* with a N-terminal long linker |  | (12) |
| psfGFP-11-II | carrying the β-hairpin GFP11 (residues 213-233) of the *sfGFP* with a N-terminal short linker |  | (12) |
| pZWC48 | *P_tna_* control of a *sfGFP10* with a C-terminal long linker and another *sfGFP10* with a C-terminal long linker | GFP10-F1, GFP10-R1, GFP10-F2, GFP10-R2 | This study |
| pZWC50 | *P_tna_* control of a *sfGFP10* with a C-terminal short linker and another *sfGFP10* with a C-terminal short linker | GFP10-F1, GFP10-R3, GFP10-F3, GFP10-R2 | This study |
| pZWC51 | *P_tna_* control of a *sfGFP11* with a N-terminal short linker and another *sfGFP11* with a N-terminal short linker | GFP11-F1, GFP11-R1, GFP11-F2, GFP11-R2 | This study |
| pZWC52 | *P_tna_* control of a *sfGFP11* with a N-terminal long linker and another *sfGFP11* with a N-terminal long linker | GFP11-F1, GFP11-R3, GFP11-F3, GFP11-R2 | This study |
| pZWC55 | *P_tna_* control of *sfGFP1-9*, *sfGFP10* with a C-terminal short linker, and *sfGFP11* with a N-terminal short linker | sfGFP1-F, sfGFP9-R1, sfGFP10S-F, sfGFP10S-R1, sfGFPS11-F1, sfGFPS11-R | This study |
| pZWC56 | *P_tna_* control of *sfGFP1-9*, *sfGFP10* with a C-terminal short linker, and *sfGFP11* with a C-terminal short linker | sfGFP1-F, sfGFP9-R1, sfGFP10S-F, sfGFP10S-R2, sfGFP11S-F, sfGFP11S-R | This study |
| pZWC57 | *P_tna_* control of *sfGFP1-9*, *sfGFP10* with a N-terminal short linker, and *sfGFP11* with a N-terminal short linker | sfGFP1-F, sfGFP9-R2, sfGFPS10-F1, sfGFPS10-R1, sfGFPS11-F2, sfGFPS11-R | This study |
| pZWC61 | *P_tna_* control of *sfGFP1-9*, *sfGFP10* with a C-terminal short linker, and *sfGFP11* with a N-terminal long linker | sfGFP1-F, sfGFP9-R1, sfGFP10S-F, sfGFP10S-R3, sfGFPL11-F, sfGFPS11-R | This study |
| pZWC62 | *P_tna_* control of *sfGFP1-9*, *sfGFP10* with a C-terminal short linker, and *sfGFP11* with a C-terminal long linker | sfGFP1-F, sfGFP9-R1, sfGFP10S-F, sfGFP10S-R2, sfGFP11S-F, sfGFP11L-R | This study |
| pZWC63 | *P_tna_* control of *sfGFP1-9*, *sfGFP10* with a N-terminal short linker, and *sfGFP11* with a N-terminal long linker | sfGFP1-F, sfGFP9-R2, sfGFPS10-F1, sfGFPS10-R2, sfGFPL11-F2, sfGFPS11-R | This study |
| pZWC64 | *P_tna_* control of *sfGFP1-9*, *sfGFP10* with a C-terminal long linker, and *sfGFP11* with a N-terminal short linker | sfGFP1-F, sfGFP9-R1, sfGFP10S-F, sfGFP10L-R, sfGFPS11-F3, sfGFPS11-R | This study |
| pZWC78 | *P_tna_* control of *sfGFP1-9*, *sfGFP10-cdpB1*, and *sfGFP11* with a C-terminal long linker | cdpB1-F1, cdpB1-R1 | This study |
| pZWC91 | *P_tna_* control of *sfGFP1-9*, *sfGFP10-ftsZ1*, and *cdpB1-sfGFP11* | ftsZ1-F1, ftsZ1-R1, cdpB1-F2, cdpB1-R2 | This study |
| pZWC92 | *P_tna_* control of *sfGFP1-9*, *sfGFP10-cdpB1*, and *sfGFP11-ftsZ1* | ftsZ1-F2, ftsZ1-R2, cdpB1-F3, cdpB1-R3 | This study |
| pZWC93 | *P_tna_* control of *sfGFP1-9*, *sfGFP10-cdpB1*, and *ftsZ1-sfGFP11* | ftsZ1-F3, ftsZ1-R3, cdpB1-F4, cdpB1-R4 | This study |
| pZWC94 | *P_tna_* control of *sfGFP1-9*, *cdpB1-sfGFP10*, and *ftsZ1-sfGFP11* | ftsZ1-F4, ftsZ1-R4, cdpB1-F5, cdpB1-R5 | This study |
| pZWC95 | *P_tna_* control of *sfGFP1-9*, *sfGFP10-ftsZ2*, and *cdpB1-sfGFP11* | ftsZ2-F1, ftsZ2-R1, cdpB1-F6, cdpB1-R6 | This study |
| pZWC96 | *P_tna_* control of *sfGFP1-9*, *sfGFP10-cdpB1*, and *sfGFP11-ftsZ2* | ftsZ2-F2, ftsZ2-R2, cdpB1-F7, cdpB1-R7 | This study |
| pZWC97 | *P_tna_* control of *sfGFP1-9*, *sfGFP10-cdpB1*, and *ftsZ2-sfGFP11* | ftsZ2-F3, ftsZ2-R3, cdpB1-F8, cdpB1-R8 | This study |
| pZWC98 | *P_tna_* control of *sfGFP1-9*, *cdpB1-sfGFP10*, and *ftsZ2-sfGFP11* | ftsZ2-F4, ftsZ2-R4, cdpB1-F9, cdpB1-R9 | This study |
| pZWC99 | *P_tna_* control of *sfGFP1-9*, *sfGFP10-sepF*, and *cdpB1-sfGFP11* | sepF-F1, sepF-R1, cdpB1-F10, cdpB1-R10 | This study |
| pZWC100 | *P_tna_* control of *sfGFP1-9*, *sfGFP10-sepF*, and *sfGFP11-cdpB1* | sepF-F2, sepF-R2, cdpB1-F11, cdpB1-R11 | This study |
| pZWC101 | *P_tna_* control of *sfGFP1-9*, *sfGFP10-cdpB1*, and *sepF-sfGFP11* | sepF-F3, sepF-R3, cdpB1-F12, cdpB1-R12 | This study |
| pZWC102 | *P_tna_* control of *sfGFP1-9*, *sepF-sfGFP10*, and *cdpB1-sfGFP11* | sepF-F4, sepF-R4, cdpB1-F13, cdpB1-R13 | This study |
| pZWC144 | *P_tna_* control of *sfGFP1-9*, *sfGFP10* with a C-terminal short linker, and *sfGFP11-ftsZ2* | ftsZ2-F5, ftsZ2-R5 | This study |
| pZWC145 | *P_tna_* control of *sfGFP1-9*, *cdpB1-sfGFP10*, and *sfGFP11* with a N-terminal long linker | cdpB1-F15, cdpB1-R15 | This study |
| pZWC146 | *P_tna_* control of *sfGFP1-9*, *sfGFP10* with a N-terminal short linker, and *ftsZ2-sfGFP11* | ftsZ2-F6, ftsZ2-R6 | This study |
| pZWC147 | *P_tna_* control of *sfGFP1-9*, *sfGFP10-cdpB1*, and *sfGFP11* with a N-terminal short linker | cdpB1-F16, cdpB1-R16 | This study |
| pZWC148 | *P_tna_* control of *sfGFP1-9*, *sfGFP10* with a C-terminal short linker, and *sepF-sfGFP11* | sepF-F5, sepF-R5 | This study |
| pZWC149 | *P_tna_* control of *sfGFP1-9*, *sfGFP10* with a N-terminal short linker, and *cdpB1-sfGFP11* | cdpB1-F17, cdpB1-R17 | This study |
| pZWC150 | *P_tna_* control of *sfGFP1-9*, *sepF-sfGFP10*, and *sfGFP11* with a N-terminal short linker | sepF-F6, sepF-R6 | This study |
| pZWC165 | *P_tna_* control of *sfGFP1-9*, *sfGFP10* with a C-terminal short linker, and *sfGFP11* with a C-terminal long linker | sfGFP10S-F, sfGFP10S-R4 | This study |
| pZWC166 | *P_tna_* control of *sfGFP1-9*, *sfGFP10* with a C-terminal short linker, and *sfGFP11* with a N-terminal long linker | sfGFP10S-F, sfGFP10S-R7 | This study |
| pZWC167 | *P_tna_* control of *sfGFP1-9*, *sfGFP10* with a N-terminal short linker, and *sfGFP11* with a N-terminal long linker | sfGFPS10-F2, sfGFPS10-R3 | This study |
| pZWC168 | *P_tna_* control of *sfGFP1-9*, *sfGFP10* with a C-terminal short linker, and *sfGFP11* with a N-terminal short linker | sfGFP10S-F, sfGFP10S-R6 | This study |
| pZWC169 | *P_tna_* control of *sfGFP1-9*, *sfGFP10* with a N-terminal short linker, and *sfGFP11* with a N-terminal short linker | sfGFPS10-F2, sfGFPS10-R4 | This study |

| **Table S3. Primers used in this study.** | |
| --- | --- |
| **Primer name** | **Oligonucleotides used in construction (5′→3′) *** |
| Clal-EcoRI-F | GCGGAATTCATCGATCTGCAGCCCGGG |
| Clal-BamHI-R | ATCGGGATCCCCCGGGCTGCAGATCGAT |
| cdpB1-NdeI-F | ATGCCATATGGACGGGACGCCCGAAGAGAT |
| cdpB1-BamHI-R1 | ATCGGGATCCTCAGGCGACCAGCGACTCT |
| P_native_- ^NJ7G^cdpB1-ApaI-F | ATCGGGGCCCCCTCACCGGCGGCCACCGG |
| P_native_- ^NJ7G^cdpB1-BamHI-R | ATCGGGATCCTCAGGCCAGCAGTTCGTCCT |
| P_native_- cdpB1-ApaI-F | ATCGGGGCCCGAAATCGCCTACCACAGCC |
| P_native_- ^HAH^cdpB1-ApaI-F | ATCGGGGCCCGGCCGACAGCCGATAGGTA |
| P_native_- ^HAH^cdpB1-BamHI-R | ATGGGATCCCTAGGCGGGGACTTCCTCT |
| cdpB1-BamHI-R2 | ATCGGGATCCGGCGACCAGCGACTCTTCGT |
| cdpB2-NdeI-F | ATGCCATATGGCCGACATACTCGCCGAG |
| cdpB2-BamHI-R | ATGCGGATCCCCGTTGGACGACGATGTAA |
| cdpB3-NdeI-F | ATGCCATATGGTCCCCCAGTCACCGACG |
| cdpB3-BamHI-R | ATCGGGATCCGTCGGCGGCTTCGAGCGCC |
| P.tna (J7)-AflII-U-F | AACCCTTGCAATTGCTTAAGGACTTCGACGACTACTTCGA |
| P.tna (J7)-U-R | CTTGGGGAATGTCGTCCATACCCGCTCAACGCCGTCT |
| ^NJ7G^cdpB1-U-F | TAGACGGCGTTGAGCGGGTATGGACGACATTCCCCAAG |
| ^NJ7G^cdpB1-U-R | GTTCTTCTCCTTTACTCATGGCCAGCAGTTCGTCCTCT |
| gfp-U-F1 | AGAGGACGAACTGCTGGCCATGAGTAAAGGAGAAGAAC |
| gfp-SphI-U-R | ATTACGCCAAGCTTGCATGCTTATTTGTATAGTTCATCC |
| P.tna-EcoRI-U-F | CCTTTCGTCTTCAAGAATTCGACTTCGACGACTACTTCG |
| P.tna-U-R | GGTAGATTCTCCATTCCCATatgcgcaataggtccgcg |
| ^HAH^cdpB1-U-F | tcgcggacctattgcgcatATGGGAATGGAGAATCTACC |
| ^HAH^cdpB1-U-R | tcttctcctttactcatggatccGGCGGGGACTTCCTCTTCCTCG |
| gfp-U-F2 | CGAGGAAGAGGAAGTCCCCGCCggatccatgagtaaaggagaaga |
| gfp-ClaI-U-R | AATCAAAGAAGCTTATCGATTTATTTGTATAGTTCATCCA |
| P_native_-cdpB2-ApaI-F | ATCGGGGCCCAAGCGTATCGGATTGTGAC |
| cdpB2-BamHI-R | GGCGGATCCCTACCGTTGGACGACGATG |
| P_native_-cdpB3-ApaI-F | ATCGGGGCCCGGTGGGTCTACGGAAGGAA |
| cdpB3-BamHI-R | ATCGGGATCCTCAGTCGGCGGCTTCGAGC |
| P_native_-ftsZ1-ApaI-F | ATCGGGGCCCCGACGCCGCGCTCTCCGCC |
| ftZ1-BamHI-R1 | ATCGGGATCCCTCGACGTAGTCGATGTCT |
| ftsZ1-NdeI-F | ATGCCATATGGACTCTATCGTCGGCGAC |
| ftsZ1-BamHI-R2 | ATCGGGATCCCTACTCGACGTAGTCGATG |
| FtsZ2-NdeI-F | ATGCCATATGCAGGATATCGTTCGCGAGG |
| FtsZ2-BamHI-R2 | ATCGGGATCCTTACCGGATGACGTCGAGA |
| sepF-NdeI-F | CGCCATATGGGTATCATGAGTAAGATT |
| sepF-BamHI-R1 | ATCGGGATCCGCCGTTGAGCTTCTGCCGC |
| sepF-BamHI-R2 | ATGCGGATCCTCAGCCGTTGAGCTTCTGC |
| cdrS -NdeI-F | ATGCCATATGGAGCGTGTGACACTACGA |
| cdrS-BamHI-R | ATCGGGATCCTTACACCTTTGCCCAGCC |
| P_native_-cdpB2-BamHI-F | ATCGGGATCCAAGCGTATCGGATTGTGAC |
| cdpB2-NotI-R | GCGGCGGCCGCCTACCGTTGGACGACGATGTA |
| ftsZ1-Notl-U-F | ATGAACTATACAAATAAGCGGCCGCATGGACTCTATCGTCGGC |
| mCH-Notl-U-R | GCGATGGTCCAGAGGTGCGGCCGCTTACTTGTACAGCTCGTCC |
| ftsZ2-Notl-U-F | ATGAACTATACAAATAAGCGGCCGCATGCAGGATATCGTTCGC |
| sepF-Notl-U-F | ATGAACTATACAAATAAGCGGCCGCATGGGTATCATGAGTAAG |
| sepF-BamHl-U-R | cagcggagccagcggatccGCCGTTGAGCTTCTGCCGC |
| mCH-U-F2 | GCGGCAGAAGCTCAACGGCggatccgctggctccgctg |
| p.phaRP-ApaI-F | ATCGGGGCCCTAATCTCGTGGTATCTCT |
| p.phaRP-NdeI-R | CGCCATATGTCAGGATCCCATCTCCTAA |
| ftsZ1-NdeI-U-F | TAGGAGATGGGATCCTGACATATGgactctatcgtcggcga |
| gfp-U-R1 | gaacgatatcctgcatgcggccgcttatttgtatagttcatcc |
| ftsZ2-U-F | ggatgaactatacaaataagcggccgcatgcaggatatcgttc |
| sepF-NdeI-U-F | GTTAGGAGATGGGATCCTGACATATGGGTATCATGAGTAAGAT |
| gfp-U-R2 | gccgacgatagagtccatgcggccgcttatttgtatagttcatccat |
| ftsZ1-U-F | atggatgaactatacaaataagcggccgcatggactctatcgtcggc |
| cdpB2-NdeI-U-F | TTAGGAGATGGGATCCTGACATATGGCCGACATACTCGCCG |
| cdpB3-NdeI-U-F | TTAGGAGATGGGATCCTGACATATGGTCCCCCAGTCACCGA |
| gfp-U-R3 | CATGATACCCATgcggccgcttatttgtatagttcatcc |
| cdpB1-NotI-U-F | gatgaactatacaaataagcggccgcATGGACGGGACGCCCGAA |
| hvo_1607-NdeI-F | ATGCcatATGTCAGTTCTCTCGACCG |
| hvo_1607-BamHI-R | CGCggatccGAGATCCTTGGCGACGCGA |
| hvo_1607-NotI-F | ATGCggatccATGTCAGTTCTCTCGACCG |
| hvo_1607-NotI-R | CGCgcggccgcTCAGAGATCCTTGGCGACG |
| cdpB1UP-Hindlll-U-F | GAGGTCGACGGTATCGATAAGCTTTGCTGGACAACGGCGCTGAC |
| cdpB1UP-U-R | CGAAGTAGTCGTCGAAGTCGCATGCATCCGCCAACATGGGATGCC |
| p.tna-cdpB1-U-F | GGCATCCCATGTTGGCGGATGCATGCGACTTCGACGACTACTTCG |
| cdpB1-BamHI-U-R | GGCCGCTCTAGAACTAGTGGATCCTCAGGCGACCAGCGACTCT |
| p.fdx-hdrB-F | GTCGCATGCCGTGGATAAAACCCCTCGTT |
| p.fdx-hdrB-R | ATCGGCATGCTTACTCATCGGATTCCTCC |
| sepFUP-Hindlll-U-F | ATCGAAGCTTAAGGTCGTCGAATCCATCCA |
| sepFUP-U-R | CGAAGTAGTCGTCGAAGTCGCATGCAGGACGTAGAGAGGCCACA |
| p.tna-sepF-U-F | TGTGGCCTCTCTACGTCCTGCATGCGACTTCGACGACTACTTCG |
| sepF-BamHI-U-R | CTCTAGAACTAGTGGATCCTCAGCCGTTGAGCTTCTGCC |
| ^NJ7G^cdpB1UP-Hindlll-U-F | GGCGGCCGCTCCGCGAGTAAGCTTGATCTCGACGCCGAGATGC |
| ^NJ7G^cdpB1UP-U-R | CGAAGTAGTCGTCGAAGTCAAGTGGCCTCATGCGTGGC |
| P.tna-^NJ7G^cdpB1-U-F | GCCACGCATGAGGCCACTTGACTTCGACGACTACTTCG |
| ^NJ7G^cdpB1-AflII-U-R | CGTAATACGACTCACTTAAGTCAGGCCAGCAGTTCGTCCT |
| ^HAH^cdpB1UP- KpnI-U-F | CAGTTCCGCTAAGGTACCGAGGAGGGTCCGCGAATC |
| ^HAH^cdpB1UP-U-R | CGAAGTAGTCGTCGAAGTCAAGTGGCCTCATGCGTGGC |
| P.tna-^HAH^cdpB1-U-F | GCCACGCATGAGGCCACTTGACTTCGACGACTACTTCG |
| ^HAH^cdpB1- HindIII-U-R | TCACTATAGGGAGAAGCTTCTAGGCGGGGACTTCCTCT |
| SUMO-SepF-BsaI-F | GATCGGTCTCAAGGTATGGACGGGACGCCCGAAG |
| SepF-XbaI-R | ATGGTCTAGATCAGGCGACCAGCGACTCT |
| SUMO-CdpB-BsaI-F | GATCGGTCTCAAGGTATGGACGGGACGCCCGAAG |
| CdpB-XbaI-R | ATGGTCTAGATCAGGCGACCAGCGACTCT |
| cdpB1UP-Hindlll-U-F | GACGGTATCGATAAGCTTTGCTGGACAACGGCGCTGA |
| cdpB1UP-U-R | CGGAAACGGGAGTTAGAACGATCCGCCAACATGGGATGCC |
| cdpB1down-U-F | GGCATCCCATGTTGGCGGATCGTTCTAACTCCCGTTTCCG |
| cdpB1down-BamHI-U-R | CGCTCTAGAACTAGTGGATCCGAGACGCTCGACTTCAAGC |
| cdpB2UP-Hindlll-U-F | GACGGTATCGATAAGCTTGATGAACGGCGTCCCCGT |
| cdpB2UP-U-R | TGAGAATCTATCCTGAATACACACCCCTTCCTCGTC |
| cdpB2down-U-F | GACGAGGAAGGGGTGTGTATTCAGGATAGATTCTCA |
| cdpB2down-BamHI-U-R | CTCTAGAACTAGTGGATCCGGGAACCTGCTCGTCGTCG |
| cdpB3UP-Hindlll-U-F | GACGGTATCGATAAGCTTAAGCGAGACGACGACTGA |
| cdpB3UP-U-R | TCGAGCGAAGTCGCCGCGGACCTCCGGTATCGGACGT |
| cdpB3down-U-F | ACGTCCGATACCGGAGGTCCGCGGCGACTTCGCTCGA |
| cdpB3down-BamHI-U-R | CTCTAGAACTAGTGGATCCGGCGTCTCCGACGACGGCCG |
| P.tna (J7)-U-R1 | TCAGCGAGTATATCGCTCATacccgctcaacgccgtcta |
| ^NJ7G^cdpB2-U-F | tagacggcgttgagcgggtATGAGCGATATACTCGCTGA |
| ^NJ7G^cdpB2-U-R | gttcttctcctttactcatGCGCTGGACGACAATGTAGT |
| gfp-U-F3 | ACTACATTGTCGTCCAGCGCatgagtaaaggagaagaac |
| P.tna (J7)-U-R2 | TTCGTCCCGTCGGAGCCCATacccgctcaacgccgtcta |
| ^NJ7G^cdpB3-U-F | tagacggcgttgagcgggtATGGGCTCCGACGGGACGAA |
| ^NJ7G^cdpB3-U-R | gttcttctcctttactcatCGTGCCGGCGTTTCGGCCC |
| gfp-U-F4 | GGGCCGAAACGCCGGCACGatgagtaaaggagaagaac |
| ^HAH^cdpB2-U-F | tcgcggacctattgcgcatATGGCTGCAATACTCGCAGA |
| ^HAH^cdpB2-U-R | tctcctttactcatggatccGCGTTTGACGATCATGTGGT |
| gfp-U-F5 | CCACATGATCGTCAAACGCggatccatgagtaaaggag |
| ^HAH^cdpB3-U-F | tcgcggacctattgcgcatATGCGAACGGTCCTCGCAAA |
| ^HAH^cdpB3-U-R | tctcctttactcatggatccCGAGTCCACAGTATCGCCGC |
| gfp-U-F6 | CGGCGATACTGTGGACTCGggatccatgagtaaaggag |
| GFP10-F1 | GCGGACCTATTGCGCATATGGACTCACTATAGGGAGACCC |
| GFP10-R1 | TAGTGGTCGTCAGGCAGGTCGGATCCCCCGCCAGCGCTCC |
| GFP10-F2 | GGAGCGCTGGCGGGGGATCCGACCTGCCTGACGACCACTA |
| GFP10-R2 | GAGGTGCGGCCTTAGCTAGCACTGTGCTGGATATCTGCAG |
| GFP10-F3 | GTTCCGGTGGTGGAGGATCCGACCTGCCTGACGACCACTA |
| GFP10-R3 | TAGTGGTCGTCAGGCAGGTCGGATCCTCCACCACCGGAAC |
| GFP11-F1 | GCGGACCTATTGCGCATATGGGCTAGCGTTTAAACTTAAG |
| GFP11-R1 | GAACCACCGCCACCGGATCCGCTAGCATCGGTAATGCCAG |
| GFP11-F2 | CTGGCATTACCGATGCTAGCGGATCCGGTGGCGGTGGTTC |
| GFP11-R2 | GAGGTGCGGCCTTAGCTAGCTCGAGGCTGATCAGCGGGCT |
| GFP11-F3 | CTGGCATTACCGATGCTAGCGGATCCGGCGCTGGCGGAAG |
| GFP11-R3 | CTTCCGCCAGCGCCGGATCCGCTAGCATCGGTAATGCCAG |
| sfGFP1-F | TTCGCGGACCTATTGCGCATATGCGCAAAGGCGAAGAACT |
| sfGFP9-R1 | TGGTCGTCAGGCAGGTCCATGAATTCTTAAACCGGGCCATCGCCAA |
| sfGFP10S-F | TTGGCGATGGCCCGGTTTAAGAATTCATGGACCTGCCTGACGACCA |
| sfGFP10S-R1 | GAACCACCGCCACCGGATCCTCTAGAAAGCTTGGATCCTCCACCACCGGAAC |
| sfGFPS11-F1 | GTTCCGGTGGTGGAGGATCCAAGCTTTCTAGAGGATCCGGTGGCGGTGGTTC |
| sfGFPS11-R | CCAGAGGTGCGGCCTTAGCTAGCATCGGTAATGCCAGCCG |
| sfGFP10S-R2 | ACCATATGGTCTCGCTTTTCCATAAGCTTGGATCCTCCACCACCGGAAC |
| sfGFP11S-F | GTTCCGGTGGTGGAGGATCCAAGCTTATGGAAAAGCGAGACCATATGGT |
| sfGFP11S-R | GGTCCAGAGGTGCGGCCTTATCTAGAAGATGTTGATCCGCCACCAG |
| sfGFP9-R2 | CCAGACCCAACGTCAGTTCCAAGCTTGAATTCTTAAACCGGGCCATCGCCAA |
| sfGFPS10-F1 | TTGGCGATGGCCCGGTTTAAGAATTCAAGCTTGGAACTGACGTTGGGTCTGG |
| sfGFPS10-R1 | GAACCACCGCCACCGGATCCTCTAGATTAGTTCAGGTCCTTGGACAGGA |
| sfGFPS11-F2 | TCCTGTCCAAGGACCTGAACTAATCTAGAGGATCCGGTGGCGGTGGTTC |
| sfGFP10S-R3 | CTTCCGCCAGCGCCGGATCCTCTAGAAAGCTTGGATCCTCCACCACCGGAAC |
| sfGFPL11-F1 | GTTCCGGTGGTGGAGGATCCAAGCTTTCTAGAGGATCCGGCGCTGGCGGAAG |
| sfGFP11L-R | GGTCCAGAGGTGCGGCCTTATCTAGAAGATGTTGAGCCGCCACTAG |
| sfGFPS10-R2 | CTTCCGCCAGCGCCGGATCCTCTAGATTAGTTCAGGTCCTTGGACAGGA |
| sfGFPL11-F2 | TCCTGTCCAAGGACCTGAACTAATCTAGAGGATCCGGCGCTGGCGGAAG |
| sfGFP10L-R | GAACCACCGCCACCGGATCCTCTAGAAAGCTTGGATCCCCCGCCAGCGCTCC |
| sfGFPS11-F3 | GGAGCGCTGGCGGGGGATCCAAGCTTTCTAGAGGATCCGGTGGCGGTGGTTC |
| cdpB-F1 | GGTGGTGGAGGATCCATGGACGGGACGCCC |
| cdpB-R1 | GTCTCGCTTTTCCATTTAGGCGACCAGCGACTCTTC |
| ftsZ1-F1 | GCTGGCGGGGGATCCATGGACTCTATCGTCGGCGA |
| ftsZ1-R1 | CCCGTCCATTCTAGATTACTCGACGTAGTCGATGTCTT |
| cdpB-F2 | AAGACATCGACTACGTCGAGTAATCTAGAATGGACGGG |
| cdpB-R2 | TGCTCTAGAGGCGACCAGCGACTCTTC |
| cdpB-F3 | GGTGGTGGAGGATCCATGGACGGGACGCCC |
| cdpB-R3 | GTCTCGCTTTTCCATTTAGGCGACCAGCGACTCTTC |
| ftsZ1-F2 | GGCGGCTCAACATCTATGGACTCTATCGTCGGCGA |
| ftsZ1-R2 | AGAGGTGCGGCCTTACTCGACGTAGTCGATGTCTT |
| cdpB-F4 | GGTGGTGGAGGATCCATGGACGGGACGCCC |
| ftsZ1-F3 | TCGCTGGTCGCCTAAATGGACTCTATCGTCGGCGA |
| ftsZ1-R3 | GCCAGCGCCGGATCCCTCGACGTAGTCGATGTCTT |
| cdpB-F5 | GCCCGGTTTAAGAATTCATGGACGGGACGCCC |
| cdpB-R5 | CCCAACGTCAGTTCCGGCGACCAGCGACTCTTC |
| ftsZ1-F4 | AAGGACCTGAACTAAATGGACTCTATCGTCGGCGA |
| ftsZ1-R4 | GCCAGCGCCGGATCCCTCGACGTAGTCGATGTCTT |
| ftsZ2-F1 | GCTGGCGGGGGATCCATGCAGGATATCGTTCGC |
| ftsZ2-R1 | CCCGTCCATTCTAGATTACCGGATGACGTCGAGACC |
| cdpB-F6 | GGTCTCGACGTCATCCGGTAATCTAGAATGGACGGG |
| cdpB-R6 | GCCACCGGATCCTCTAGAGGCGACCAGCGACTCTTC |
| cdpB-F7 | GGTGGTGGAGGATCCATGGACGGGACGCCC |
| cdpB-R7 | GCTTTTCCATTTAGGCGACCAGCGACTCTTC |
| ftsZ2-F2 | GGCGGCTCAACATCTATGCAGGATATCGTTCGC |
| ftsZ2-R2 | AGAGGTGCGGCCTTACCGGATGACGTCGAGACC |
| cdpB-F8 | GGTGGTGGAGGATCCATGGACGGGACGCCC |
| cdpB-R8 | GCGAACGATATCCTGCATTTAGGCGACCAGCGACTCTTC |
| ftsZ2-F3 | TCGCTGGTCGCCTAAATGCAGGATATCGTTCGC |
| ftsZ2-R3 | CGCCAGCGCCGGATCCCGGATGACGTCGAGACC |
| cdpB-F9 | CCGGTTTAAGAATTCATGGACGGGACGCCC |
| cdpB-R9 | CCCAACGTCAGTTCCGGCGACCAGCGACTCTTC |
| ftsZ2-F4 | AAGGACCTGAACTAAATGCAGGATATCGTTCGC |
| ftsZ2-R4 | GCCAGCGCCGGATCCCCGGATGACGTCGAGACC |
| sepF-F1 | GGTGGTGGAGGATCCATGGGTATCATGAGTAAGAT |
| sepF-R1 | CCCGTCCATTCTAGATTAGCCGTTGAGCTTCTGCC |
| cdpB-F10 | AAGCTCAACGGCTAATCTAGAATGGACGGGACGCCC |
| cdpB-R10 | CCGCCACCGGATCCTCTAGAGGCGACCAGCGACTCTTC |
| sepF-F2 | GGTGGTGGAGGATCCATGGGTATCATGAGTAAGAT |
| sepF-R2 | GTCTCGCTTTTCCATTTAGCCGTTGAGCTTCTGCC |
| cdpB-F11 | TCAACATCTTCTAGAATGGACGGGACGCCC |
| cdpB-R11 | GCCTTATCTAGATTAGGCGACCAGCGACTCTTC |
| cdpB-F12 | GGAGGATCCAAGCTTATGGACGGGACGCCC |
| cdpB-R12 | ACCCATAAGCTTTTAGGCGACCAGCGACTCTTC |
| sepF-F3 | GTCGCCTAAAAGCTTATGGGTATCATGAGTAAGAT |
| sepF-R3 | ACCGCCACCGGATCCGCCGTTGAGCTTCTGCC |
| sepF-F4 | CCGGTTTAAGAATTCATGGGTATCATGAGTAAGAT |
| sepF-R4 | CCCAACGTCAGTTCCGCCGTTGAGCTTCTGCC |
| cdpB-F13 | AAGGACCTGAACTAAATGGACGGGACGCCC |
| cdpB-R13 | ACCGCCACCGGATCCGGCGACCAGCGACTCTTC |
| cdpB-F14 | GGTGGTGGAGGATCCATGGACGGGACGCCC |
| cdpB-R14 | GCCAGCGCCGGATCCCATTTAGGCGACCAGCG |
| ftsZ2-F5 | GGCGGCTCAACATCTATGCAGGATATCGTTCGC |
| ftsZ2-R5 | AGAGGTGCGGCCTTACCGGATGACGTCGAGACC |
| cdpB-F15 | CCGGTTTAAGAATTCATGGACGGGACGCCC |
| cdpB-R15 | CCCAACGTCAGTTCCGGCGACCAGCGACTCTTC |
| ftsZ2-F6 | AAGGACCTGAACTAAATGCAGGATATCGTTCGC |
| ftsZ2-R6 | GCCAGCGCCGGATCCCCGGATGACGTCGAGACC |
| cdpB-F16 | TGGAGGATCCAAGCTTATGGACGGGACGCCC |
| cdpB-R16 | ACCGCCACCGGATCCCATAAGCTTTTAGGCGACCA |
| sepF-F5 | GGTGGTGGAGGATCCTAAAAGCTTATGGGTATCATGAG |
| sepF-R5 | ACCGCCACCGGATCCGCCGTTGAGCTTCTGCC |
| cdpB-F17 | AAGGACCTGAACTAAATGGACGGGACGCCC |
| cdpB-R17 | ACCGCCACCGGATCCGGCGACCAGCGACTCTTC |
| sepF-F6 | CCGGTTTAAGAATTCATGGGTATCATGAGTAAGAT |
| sepF-R6 | CCCAACGTCAGTTCCGCCGTTGAGCTTCTGCC |
| sfGFP10S-R4 | GTCTCGCTTTTCCATTTAGGATCCTCCACCACCG |
| sfGFP10S-R5 | GCCAGCGCCGGATCCCATTTAGGATCCTCCACCACCG |
| sfGFPS10-F2 | GCGATGGCCCGGTTTAAATGGGAACTGACGTTGGG |
| sfGFPS10-R3 | GCCAGCGCCGGATCCCATTTAGTTCAGGTCCTTGGACAGG |
| sfGFP10S-R6 | ACCGCCACCGGATCCCATTTAGGATCCTCCACCACCG |
| sfGFP10S-R7 | GCCAGCGCCGGATCCCATTTAGGATCCTCCACCACCG |
| sfGFPS10-R4 | ACCGCCACCGGATCCCATTTAGTTCAGGTCCTTGGACAGG |
| P_native_-sepF-F | CCCGGTACCGGGCCCAACCGACGCCGTGGA |
| P_native_-sepF-R | CGAGAATCTTACTCATGATACCCATAGGACGTAGAGAGGCCAC |
| sepF-F7 | GTGGCCTCTCTACGTCCTATGGGTATCATGAGTAAGATTCTCG |
| sepF-flag-R1 | CTTATCGTCGTCATCCTTGTAATCGGATCCTCCACCACCGGA |
| sepF-flag-R2 | TCCAGAGGTGCGGCCGCTTACTTATCGTCGTCATCCTTGTAA |
| P_native_-cdpB1-F | CGACGATAAGTAAGCGGCCGCGAAATCGCCTACCACAGCC |
| gfp-R | CAGAGGTGCGGCCGCTTATTTGTATAGTTCATCCATGCCA |

| **Table S4. Chemicals and antibodies used in this study.** | | |
| --- | --- | --- |
| **REAGENT or RESOURCE** | **SOURCE** | **IDENTIFIER** |
| Yeast Extract | Becton-Dickinson（BD） | 212720 |
| Tryptone | Becton-Dickinson（BD） | 211699 |
| Peptone | Oxoid | LP0037B |
| Casamino Acids | Becton-Dickinson（BD） | 223050 |
| Agar | Becton-Dickinson（BD） | 214010 |
| NaOH | Sinopharm Chemical Reagent Co. Ltd | 10019762 |
| NaCl | Sinopharm Chemical Reagent Co. Ltd | 10019308 |
| MgSO_4_·7H_2_O | Sinopharm Chemical Reagent Co. Ltd | 10013018 |
| MgCl_2_·6H_2_O | Sinopharm Chemical Reagent Co. Ltd | 20191016 |
| KCl | Sinopharm Chemical Reagent Co. Ltd | 10016318 |
| FeSO_4_·7H_2_O | Sinopharm Chemical Reagent Co. Ltd | 10012118 |
| MnCl_2_·4H_2_O | Sinopharm Chemical Reagent Co. Ltd | TM209525G |
| CaCl_2_ | Sinopharm Chemical Reagent Co. Ltd | 100195861 |
| FeCl_3_ | Sinopharm Chemical Reagent Co. Ltd | 81015228 |
| ZnCl_2_ | Sinopharm Chemical Reagent Co. Ltd | 10023883 |
| CuCl_2_ | Sinopharm Chemical Reagent Co. Ltd | 10007816 |
| H_3_BO_3_ | Sinopharm Chemical Reagent Co. Ltd | 10004817 |
| NiSO_4_·6H_2_O | Sinopharm Chemical Reagent Co. Ltd | 10014418 |
| Na_2_MoO_4_.2H_2_O | Sinopharm Chemical Reagent Co. Ltd | 10019816 |
| Trisodium citrate | Aladdin | S189183 |
| Sodium glutamate | Solarbio | G8010 |
| 2×Hieff Canace gold PCR Master Mix | YEASEN | 10136ES03 |
| Hieff Clone Plus One Step Cloning Kit | YEASEN | 10911ES20 |
| 2×F8 FastLong PCR MasterMix | Aidlab Biotechnologies | PC8001 |
| T4 DNA ligase | Thermo Fisher | EL0011 |
| 10×T4 DNA ligase Buffer | Thermo Fisher | B69 |
| All restriction enzymes | Thermo Fisher |  |
| 6×DNA Loading Buffer | TransGen Biotech | GH101-01 |
| 1Kb DNA Ladder | TransGen Biotech | BM201-01 |
| Uracil | Solarbio | U8010 |
| Thiamine | Solarbio | V8020 |
| Biotin | Solarbio | IB0250 |
| L-Tryptophan | Solarbio | T0011 |
| PEG600 | Sigma-Fluka | 8074861000 |
| Na_2_EDTA.2H_2_O | Solarbio | E8030 |
| Sucrose | Sinopharm Chemical Reagent Co. Ltd | 10021418 |
| HCl | Sinopharm Chemical Reagent Co. Ltd | 10011028 |
| Glycerol | Solarbio | G8190 |
| Tris-base | BioFroxx | 1115 |
| SDS-PAGE Separating Gel Buffer | Solarbio | S1051 |
| SDS-PAGE spacer Gel Buffer | Solarbio | S1052 |
| SDS | Solarbio | S8010 |
| TEMED | Fdbio Science | FD2100 |
| 30% acrylamide-bisacrylamide | Biosharp | BL513A |
| Agarose | US Everbright Inc | A2015 |
| 5-FOA | Sangon Biotech | A601555 |
| Imidazole | Solarbio | I8090 |
| DTT | Solarbio | D8220 |
| APS | Fdbio science | FD2050 |
| ColorMixed Protein Marker | Solarbio | PR1910 |
| Tween-20 | Solarbio | T8220 |
| PBS | Solarbio | P1000 |
| 20x TBST | Solarbio | T1082 |
| Skim milk powder | BioFroxx | 1172 |
| Glycine | BioFroxx | 1275 |
| Methanol | Sinopharm Chemical Reagent Co. Ltd | 10014118 |
| Ethanol | Sinopharm Chemical Reagent Co. Ltd | 10009218 |
| DMSO | Macklin | D807289 |
| Rabbit anti GFP-Tag mAb | ABclonal | AE078 |
| Rabbit anti DDDDK-Tag pAb | ABclonal | AE004 |
| ProteinFind® Anti-GFP Mouse Monoclonal Antibody | Transgen | HT801-01 |
| ProteinFind® Anti-DYKDDDDK Mouse Monoclonal Antibody | Transgen | HT201-01 |
| Protein A/G Magnetic Beads | MedChemExpress | HY-K0202 |
| Protease Inhibitor Cocktail | MedChemExpress | HY-K0010 |
| Magnetic Stand | MedChemExpress | HY-K0200 |
| Hybond-NC | Pall Corporation | 66485 |
| Smart-ECL Enhanced | Smart-Lifesciences | H31500 |
| E.Z.N.A.®Gel Extraction Kit | Omega | D2500-01 |
| TIANprep Mini Plasmid Kit | TIANGEN | 4992420 |
| [Wizard®Genomic DNA Purification Kit](https://www.promega.com.cn/products/nucleic-acid-extraction/genomic-dna/wizard-genomic-dna-purification-kit/) | Promega | A1120 |

**Construction of plasmids**

pZS1

The plasmid pZS1 (pTA1228, *P_tna_*::MCS *bla* with a ClaI site) was constructed by ligation of an EcoRI/BamHI digested DNA fragment containing a ClaI site into pTA1228 digested with the same enzymes. The DNA fragment was amplified from plasmid pTA1228 using primers Clal-EcoRI-F and Clal-BamHI-R.

pZS98

The plasmid pZS98 (pTA131, containing *cdpB1* upstream flanking sequence followed by the *P_tna_::cdpB1* cassette) was constructed by ligation of pTA131 (digested with HindIII and BamHI) with a DNA fragment containing the upstream flanking sequence of *cdpB1* followed by *P_tna_::cdpB1*. To construct the plasmid, a DNA fragment harboring the *cdpB1* upstream flanking sequence was first amplified from *H. volcanii* DS70 (DS2, Δ*pHV2*) chromosomal DNA using primers cdpB1UP-Hindlll-U-F and cdpB1UP-U-R. The *P_tna_::cdpB1* cassette was amplified from plasmid pZS103 (containing the downstream flanking sequence of *cdpB1* followed by the L11e transcription terminator and the *P_tna_::cdpB1* cassette) using primers P.tna-cdpB1-U-F and cdpB1-BamHI-U-R. The two DNA fragments were joined by overlap extension PCR to generate the fragment to be inserted into pTA131.

pZS99

The plasmid pZS99 (pTA131, containing *sepF* upstream flanking sequence followed by the cassette *P_tna_::sepF*) was constructed by recombination of pTA131 (digested with HindIII and BamHI) and a DNA fragment containing *sepF* upstream flanking sequencing followed by *P_tna_::sepF* (containing the terminal homologous sequence of pTA131). To construct the plasmid, a DNA fragment harboring the *sepF* upstream flanking sequence was amplified from *H. volcanii* DS70 (DS2, *ΔpHV2*) chromosomal DNA using primers sepFUP-Hindlll-U-F and sepFUP-U-R. The cassette *P_tna_::sepF* was amplified from plasmid pZS104 (containing sepF downstream flanking sequence followed by the L11e transcription terminator and *P_tna_::sepF*) using primers P.tna- sepF-U-F and sepF-BamHI-U-R. The two DNA fragments were joined by overlap extension PCR to generate the fragment to be inserted into pTA131.

pZS100

The plasmid pZS100 (pTA131, containing both *cdpB1* upstream and downstream flanking sequence) is a non-replicating plasmid for deletion of *cdpB1* using the pop-in/pop-out approach. It was constructed by recombination of pTA131 (digested with HindIII and BamHI) and a DNA fragment harboring the upstream and downstream flanking sequence of *cdpB1* as well as the terminal homologous sequence of pTA131. The upstream and downstream flanking sequences of *cdpB1* were amplified from *H. volcanii* DS70 (DS2, *ΔpHV2*) chromosomal DNA using primer pairs cdpB1UP-Hindlll-U-F/cdpB1UP-U-R and cdpB1down-U-F/cdpB1down-BamHI-U-R, respectively. The upstream and downstream flanking sequences of *cdpB1* were then joined by overlap extension PCR using primers cdpB1UP-Hindlll-U-F and cdpB1down-BamHI-U-R.

pZS101

The plasmid pZS101 (pIDJL114, *P_tna_::cdpB1-mCherry bla*) was constructed by ligation of a NdeI/BamHI digested DNA fragment containing *cdpB1* into pIDJL114 digested with the same enzymes. The DNA fragment was amplified from *H. volcanii* DS70 (DS2, *ΔpHV2*) chromosomal DNA using primers cdpB1-NdeI-F and cdpB1-BamHI-R2.

pZS103

The plasmid pZS103 (pIDJL40-*ftsZ1*, *P_tna_::cdpB1-gfp bla*) was constructed by ligation of a NdeI/BamHI digested DNA fragment containing *cdpB1* into pIDJL40-*ftsZ1* digested with the same enzymes. The DNA fragment was amplified from *H. volcanii* DS70 (DS2, *ΔpHV2*) chromosomal DNA using primers cdpB1-NdeI-F and cdpB1-BamHI-R2.

pZS104

The plasmid pZS104 (pIDJL40-*ftsZ1*, *P_tna_::sepF-gfp bla*) was constructed by ligation of a NdeI/BamHI digested DNA fragment containing *sepF* into pIDJL40-*ftsZ1* digested with the same enzymes. The DNA fragment was amplified from *H. volcanii* DS70 (DS2, *ΔpHV2*) chromosomal DNA using primers sepF-NdeI-F and sepF-BamHI-R1.

pZS105

The plasmid pZS105 (pZS103, *P_tna_::cdpB1-gfp-ftsZ1-mCherry bla*) was constructed by recombination of pZS103 (digested with NotI) and a DNA fragment harboring *ftsZ1-mCherry* followed by the terminal homologous sequence of pZS103. The DNA fragment was amplified from plasmid pIDJL114 (*P_tna_::ftsZ1-mCherry bla*) using primers ftsZ1-Notl-U-F and mCH-Notl-U-R.

pZS106

The plasmid pZS106 (pZS103, *P_tna_::cdpB1-gfp-ftsZ2-mCherry bla*) was constructed by recombination of pZS103 (digested with NotI) and a DNA fragment harboring *ftsZ2-mCherry* followed by the terminal homologous sequence of pZS103. The DNA fragment was amplified from plasmid pIDJL115 (*P_tna_::ftsZ2-mCherry bla*) using primers ftsZ2-Notl-U-F and mCH-Notl-U-R.

pZS107

The plasmid pZS107 (pZS103, *P_tna_::cdpB1-gfp-sepF-mCherry bla*) was constructed by recombination of pZS103 (digested with NotI) and a DNA fragment harboring *sepF-mCherry* followed by the terminal homologous sequence of pZS103. *sepF* coding sequence was amplified from *H. volcanii* DS70 (DS2, *ΔpHV2*) chromosomal DNA using primers sepF-Notl-U-F and sepF-BamHl-U-R. *mCherry* coding sequence was amplified from plasmid pIDJL114 (*P_tna_::ftsZ1-mCherry bla*) using primers mCH-U-F2 and mCH-Notl-U-R. *sepF* and *mCherry* were joined by overlap extension PCR using primers sepF-Notl-U-F and mCH-Notl-U-R to generate the *sepF-mCherry* cassette.

pZS111

The plasmid pZS111 (pTA131, containing *cdpB1* upstream flanking sequence followed by the cassettes *P_fdx_::hdrB and P_tna_::cdpB1*) is a non-replicating plasmid for the generation of CdpB1 depletion strain. It was constructed by ligation of a SphI digested DNA fragment containing *P_fdx_::hdrB* into pZS98 digested with the same enzymes. The *P_fdx_::hdrB* cassette was amplified from *H. volcanii* DS70 (DS2, *ΔpHV2*) chromosomal DNA using primers p.fdx-hdrB-F and p.fdx-hdrB-R.

pZS124

The plasmid pZS124 (pTA1228, *P_tna_::ftsZ1 bla*) was constructed by ligation of a NdeI/BamHI digested DNA fragment containing *ftsZ1* into pTA1228 digested with the same enzymes. The DNA fragment was amplified from *H. volcanii* DS70 (DS2, *ΔpHV2*) chromosomal DNA using primers ftsZ1-NdeI-F and ftsZ1-BamHI-R2.

pZS125

The plasmid pZS125 (pTA1228, *P_tna_::ftsZ2 bla*) was constructed by ligation of a NdeI/BamHI digested DNA fragment containing *ftsZ2* into pTA1228 digested with the same enzymes. The DNA fragment was amplified from *H. volcanii* DS70 (DS2, *ΔpHV2*) chromosomal DNA using primers ftsZ2-NdeI-F and ftsZ2-BamHI-R2.

pZS126

The plasmid pZS126 (pTA1228, *P_tna_::cdpB1 bla*) was constructed by ligation of a NdeI/BamHI digested DNA fragment containing *cdpB1* into pTA1228 digested with the same enzymes. The DNA fragment was amplified from *H. volcanii* DS70 (DS2, *ΔpHV2*) chromosomal DNA using primers cdpB1-NdeI-F and cdpB1-BamHI-R1.

pZS127

The plasmid pZS127 (pTA1228, *P_tna_::sepF bla*) was constructed by ligation of a NdeI/BamHI digested DNA fragment containing *sepF* into pTA1228 digested with the same enzymes. The DNA fragment was amplified from *H. volcanii* DS70 (DS2, *ΔpHV2*) chromosomal DNA using primers sepF-NdeI-F and sepF-BamHI-R2.

pZS204

The plasmid pZS204 (pTA1228, *P_tna_::cdrS bla*) was constructed by ligation of a NdeI/BamHI digested DNA fragment containing *cdrS* into pTA1228 digested with the same enzymes. The DNA fragment was amplified from *H. volcanii* DS70 (DS2, *ΔpHV2*) chromosomal DNA using primers cdrS-NdeI-F and cdrS-BamHI-R.

pZS208

The plasmid pZS208 (pIDJL40-*ftsZ1*, *P_native_::ftsZ1-gfp bla*) was constructed by ligation of an ApaI/BamHI digested DNA fragment containing *P_native_::ftsZ1* into pIDJL40-*ftsZ1* digested with the same enzymes. The DNA fragment was amplified from *H. volcanii* DS70 (DS2, *ΔpHV2*) chromosomal DNA using primers P_native_-ftsZ1-ApaI-F and ftZ1-BamHI-R1.

pZS214

The plasmid pZS214 (pIDJL40-*ftsZ1*, *p_native_::cdpB1-gfp bla*) was constructed by ligation of an ApaI/BamHI digested DNA fragment containing *p_native_::cdpB1* into pIDJL40-*ftsZ1* digested with the same enzymes. The DNA fragment was amplified from *H. volcanii* DS70 (DS2, *ΔpHV2*) chromosomal DNA using primers P_native_-cdpB1-ApaI-F and cdpB1-BamHI-R2.

pZS217

The plasmid pZS217 (pFJ6-*P_tna_*, *P_tna_:: ^NJ7G^cdpB1-gfp bla*) was constructed by recombination of pFJ6-*P_tna_* (digested with AflII and SphI) and a DNA fragment harboring the cassette *P_tna_::^NJ7G^cdpB1-gfp* followed by the terminal homologous sequence of pFJ6-*P_tna_*. The DNA fragment harboring the *P_tna_* promoter was amplified from plasmid pFJ6-*P_tna_* using primers P.tna (J7)-AflII-U-F and P.tna (J7)-U-R. *^NJ7G^cdpB1* was amplified from *Natrinema sp*.CJ7-F (*ΔpyrF*) chromosomal DNA using primers ^NJ7G^cdpB1-U-F and ^NJ7G^cdpB1-U-R. *gfp* was amplified from plasmid pIDJL40-*ftsZ1*(*P_tna_::ftsZ1-gfp bla*) using primers gfp-U-F1 and gfp-SphI-U-R. The three fragments *P_tna_, ^NJ7G^cdpB1* and *gfp* were joined by overlap extension PCR using primers P.tna (J7)-AflII-U-F and gfp-SphI-U-R to generate the cassette *P_tna_:: ^NJ7G^cdpB1-gfp.*

pZS219

The plasmid pZS219 (pTA131, harboring *sepF* upstream flanking sequence followed by the cassette P*_fdx_::hdrB* and *P_tna_::sepF*) is a non-replicating plasmid for the generation of SepF depletion strain. It was constructed by ligation of a SphI digested DNA fragment containing *P_fdx_::hdrB* into pZS98 digested with the same enzymes. The DNA fragment harboring *P_fdx_::hdrB* was amplified from *H. volcanii* DS70 (DS2, *ΔpHV2*) chromosomal DNA using primers p.fdx-hdrB-F and p.fdx-hdrB-R.

pZS234

The plasmid pZS234 (pTA1228, *P_native_::^NJ7G^cdpB1 bla*) was constructed by ligation of an ApaI/BamHI digested DNA fragment containing *P_native_::^NJ7G^cdpB1* into pTA1228 digested with the same enzymes. The DNA fragment was amplified from *Natrinema sp*. CJ7-F (*ΔpyrF*) chromosomal DNA using primers P_native_- ^NJ7G^cdpB1-ApaI-F and P_native_- ^NJ7G^cdpB1-BamHI-R.

pZS236

The plasmid pZS236 (pTA1228, *P_native_::cdpB1 bla*) was constructed by ligation of an ApaI/BamHI digested DNA fragment containing *P_native_-cdpB1* into pTA1228 digested with the same enzymes. The DNA fragment was amplified from *H.volcanii* DS70 (DS2, *ΔpHV2*) chromosomal DNA using primers P_native_-cdpB1-ApaI-Fand cdpB1-BamHI-R1.

pZS239

The plasmid pZS239 (pZS105, *P_phaR_::cdpB1-gfp-ftsZ1-mCherry bla*) was constructed by replacing the *P_tna_* promoter with the P_phaR_ promoter via ligation of an ApaI/NdeI digested DNA fragment containing *P_phaR_* into pZS105 digested with the same enzymes. The P_phaR_ promoter was amplified from plasmid pFJ6-*P_phaR_*-GFP (*P_phaR_::gfp bla*) using primers p.phaRP-ApaI-F and p.phaRP-NdeI-R.

pZS253

The plasmid pZS253 (pNBK-F, containing *^NJ7G^cdpB1* upstream flanking sequence followed by the cassette *P_tna_::^NJ7G^cdpB1*) is a non-replicating plasmid for the generation of ^NJ7G^CdpB1 depletion strain. It was constructed by recombination of pNBK-F (digested with HindIII and AflII) and a DNA fragment harboring *^NJ7G^cdpB1* upstream flanking sequence followed by the cassette *P_tna_::^NJ7G^cdpB1* and the terminal homologous sequence of pNBK-F. The upstream flanking sequence of *^NJ7G^cdpB1* was amplified from *Natrinema sp*.CJ7-F (*ΔpyrF*) chromosomal DNA using primers ^NJ7G^cdpB1UP-Hindlll-U-F and ^NJ7G^cdpB1UP-U-R. The cassette *P_tna_::^NJ7G^cdpB1* was amplified from plasmid pZS217 (containing the L11e transcription terminator and the *P_tna::_^NJ7G^cdpB1* cassette to be inserted) using primers P.tna-^NJ7G^cdpB1-U-F and ^NJ7G^cdpB1- AflII-U-R. The two DNA fragments were joined by overlap extension PCR using primers ^NJ7G^cdpB1UP-Hindlll-U-F and ^NJ7G^cdpB1- AflII-U-R and then inserted into pNBK-F.

pZS276

The plasmid pZS276 (pTA1228, *P_native_:: ^HAH^cdpB1 bla*) was constructed by ligation of an ApaI/BamHI digested DNA fragment containing *P_native_::^HAH^cdpB1* into pTA1228 digested with the same enzymes. The DNA fragment was amplified from *H. hispanica* DF60 (*ΔpyrF*) chromosomal DNA using primers P_native_- ^HAH^cdpB1-ApaI-F and P_native_- ^HAH^cdpB1-BamHI-R.

pZS280

The plasmid pZS280 (pHAR, containing *^HAH^cdpB1* upstream flanking sequence followed by the cassette *P_tna_::^HAH^cdpB1*) is a non-replicating plasmid for the generation of ^HAH^CdpB1 depletion strain. It was constructed by recombination of pHAR (digested with KpnI and HindIII) and a DNA fragment harboring *^HAH^cdpB1* upstream flanking sequence followed by the cassette *P_tna_::^HAH^cdpB1* and the terminal homologous sequence of pHAR. The upstream flanking sequence of *^HAH^cdpB1* was amplified from *H. hispanica* DF60 (*ΔpyrF*) chromosomal DNA using primers ^HAH^cdpB1UP- KpnI-U-F and ^HAH^cdpB1UP-U-R. *P_tna_::^HAH^cdpB1* was amplified from plasmid pZS306 (containing the L11e transcription terminator and *p.tna::^HAH^cdpB1* cassette to be inserted) using primers P.tna-^HAH^cdpB1-U-F and ^HAH^cdpB1-HindIII-U-R. The two DNA fragments were joined by overlap extension PCR using primers ^HAH^cdpB1UP- KpnI-U-F and ^HAH^cdpB1- HindIII-U-R and inserted into pHAR.

pZS284

The plasmid pZS284 (pZS106, *P_phaR_::cdpB1-gfp-ftsZ2-mCherry bla*) was constructed by ligation of an ApaI/NdeI digested DNA fragment containing *P_phaR_* into pZS106 digested with the same enzymes. The DNA fragment was amplified from plasmid pFJ6-*P_phaR_*-GFP (*P_phaR_::gfp bla*) using primers p.phaRP-ApaI-F and p.phaRP-NdeI-R.

pZS285

The plasmid pZS285 (pZS107, *P_phaR_::cdpB1-gfp-sepF-mCherry bla*) was constructed by ligation of an ApaI/NdeI digested DNA fragment containing *P_phaR_* into pZS107 digested with the same enzymes. The DNA fragment was amplified from plasmid pFJ6-*P_phaR_*-GFP (*P_phaR_::gfp bla*) using primers p.phaRP-ApaI-F and p.phaRP-NdeI-R.

pZS288

The plasmid pZS288 (pE-SUMO, P_T7_::*his-SUMO-SepF*, *bla*) was constructed by ligation of an BsaI/BamHI digested DNA fragment containing *sepF* into pE-SUMO digested with the same enzymes. The DNA fragment was amplified from *H. volcanii* DS70 (DS2, *ΔpHV2*) chromosomal DNA using primers SUMO-SepF-BsaI-F and SepF-XbaI-R.

pZS289

The plasmid pZS289 (pZS285, *P_phaR_::ftsZ1-gfp-ftsZ2-mCherry bla*) was constructed by recombination of pZS285 (digested with NdeI and NotI) and a DNA fragment containing the cassette *ftsZ1-gfp-ftsZ2-mCherry* followed by the terminal homologous sequence of pZS285. *ftsZ1-gfp* was amplified from plasmid pIDJL40-*ftsZ1* using primers *ftsZ1*-NdeI-U-F and *gfp*-U-R1, whereas *ftsZ2-mCherry* was amplified from plasmid pIDJL115 (*P_tna_::ftsZ2-mCherry*) using primers ftsZ2-U-F and mCH-NotI-U-R. *ftsZ1-gfp* and *ftsZ2-mCherry* were joined by overlap extension PCR using primers *ftsZ1*-NdeI-U-F and *mCH*-NotI-U-R and then inserted into pZS285 as described above.

pZS311

The plasmid pZS311 (pE-SUMO, P_T7_::*his-SUMO-CdpB1*, *bla*) was constructed by ligation of an BsaI/BamHI digested DNA fragment containing *cdpB1* into pE-SUMO digested with the same enzymes. The DNA fragment was amplified from *H. volcanii* DS70 (DS2, *ΔpHV2*) chromosomal DNA using primers SUMO-CdpB1-BsaI-F and CdpB1-XbaI-R.

pZS322

The plasmid pZS322 (pZS285, *P_phaR_::sepF-gfp-ftsZ1-mCherry bla*) was constructed by recombination of pZS285 (digested with NdeI and NotI) and a DNA fragment containing *sepF-gfp-ftsZ1-mCherry* followed by the terminal homologous sequence of pZS285. *sepF-gfp* was amplified from plasmid pZS104 (*P_tna_::sepF-gfp*) using primers sepF-NdeI-U-F and gfp-U-R2, whereas *ftsZ1-mCherry* was amplified from plasmid pIDJL114 (*P_tna_::ftsZ1-mCherry*) using primers ftsZ1-U-F and mCH-NotI-U-R. The two DNA fragments were joined by overlap extension PCR using primers *sepF*-NdeI-U-F and *mCH*-NotI-U-R and then inserted into pZS285.

pZS324

The plasmid pZS324 (pZS322, *P_phaR_::sepF-gfp-ftsZ2-mCherry bla*) was constructed by recombination of pZS322 (digested with NotI) and a DNA fragment containing *ftsZ2-mCherry* (containing the terminal homologous sequence of pZS322). The DNA fragment was amplified from plasmid pIDJL115 (*P_tna_::ftsZ2-mCherry bla*) using primers ftsZ2-Notl-U-F and mCH-Notl-U-R.

pZS306

The plasmid pZS306 (pWL502, *P_tna_::^HAH^cdpB1-gfp bla*) was constructed by recombination of pWL502 (digested with EcoRI and ClaI) and a DNA fragment containing *P_tna_::^HAH^cdpB1-gfp* followed by the terminal homologous sequence of pWL502. The *P_tna_* promoter was amplified from plasmid pIDJL40-*ftsZ1* using primers P.tna-EcoRI-U-F and P.tna-U-R, *^HAH^cdpB1* was amplified from *H. hispanica* DF60 (*ΔpyrF*) chromosomal DNA using primers ^HAH^cdpB1-U-F and ^HAH^cdpB1-U-R, and *gfp* was amplified from plasmid pIDJL40-*ftsZ1* (*P_tna_::ftsZ1-gfp bla*) using primers gfp-U-F2 and gfp-ClaI-U-R. The three DNA fragments were joined by overlap extension PCR using primers P.tna-EcoRI-U-F and gfp-ClaI-U-R to generate the *P_tna_*:: *^HAH^cdpB1-gfp* cassette.

pZS336

The plasmid pZS336 (pIDJL40-*ftsZ1*, *P_tna_::cdpB2-gfp bla*) was constructed by ligation of a NdeI/BamHI digested DNA fragment containing *cdpB2* into pIDJL40-*ftsZ1* digested with the same enzymes. The DNA fragment was amplified from *H. volcanii* DS70 (DS2, *ΔpHV2*) chromosomal DNA using primers cdpB2-NdeI-F and cdpB2-BamHI-R.

pZS337

The plasmid pZS337 (pIDJL40-*ftsZ1*, *P_tna_::cdpB3-gfp bla*) was constructed by ligation of an NdeI/BamHI digested DNA fragment containing *cdpB3* into pIDJL40-*ftsZ1* digested with the same enzymes. The DNA fragment was amplified from *H. volcanii* DS70 (DS2, *ΔpHV2*) chromosomal DNA using primers cdpB3-NdeI-F and cdpB3-BamHI-R.

pZS338

The plasmid pZS338 (pTA1228, *P_native_::cdpB3 bla*) was constructed by ligation of an ApaI/BamHI digested DNA fragment containing *P_native_::cdpB3* into pTA1228 digested with the same enzymes. The DNA fragment was amplified from *H. volcanii* DS70 (DS2, *ΔpHV2*) chromosomal DNA using primers P_native_-cdpB3-ApaI-Fand cdpB3-BamHI-R1.

pZS339

The plasmid pZS339 (pTA1228, *P_native_::cdpB2 bla*) was constructed by ligation of an ApaI/BamHI digested DNA fragment containing *P_native_::cdpB2* into pTA1228 digested with the same enzymes. The DNA fragment was amplified from *H. volcanii* DS70 (DS2, *ΔpHV2*) chromosomal DNA using primers P_native_-cdpB2-ApaI-F and cdpB2-BamHI-R.

pZS384

The plasmid pZS384 (pFJ6-*P_tna_*, *P_tna_::NJ7G_2729-gfp bla*) was constructed by recombination of pFJ6-*P_tna_* (digested with AflII and SphI) and a DNA fragment harboring the cassette *P_tna_::^NJ7G^cdpB2-gfp* followed by the terminal homologous sequence of pFJ6-*P_tna_*. The DNA fragment harboring the *P_tna_* promoter was amplified from plasmid pFJ6-*P_tna_* using primers P.tna (J7)-AflII-U-F and P.tna (J7)-U-R1. *^NJ7G^cdpB2* was amplified from *Natrinema sp*.CJ7-F (*ΔpyrF*) chromosomal DNA using primers ^NJ7G^cdpB2-U-F and ^NJ7G^cdpB2-U-R. *gfp* was amplified from plasmid pIDJL40-*ftsZ1*(*P_tna_::ftsZ1-gfp bla*) using primers gfp-U-F3 and gfp-SphI-U-R. The three fragments *P_tna_, ^NJ7G^cdpB2* and *gfp* were joined by overlap extension PCR using primers P.tna (J7)-AflII-U-F and gfp-SphI-U-R to generate the cassette *P_tna_:: ^NJ7G^cdpB2-gfp.*

pZS385

The plasmid pZS385 (pFJ6-*P_tna_*, *P_tna_::NJ7G_3497-gfp bla*) was constructed by recombination of pFJ6-*P_tna_* (digested with AflII and SphI) and a DNA fragment harboring the cassette *P_tna_::^NJ7G^cdpB3-gfp* followed by the terminal homologous sequence of pFJ6-*P_tna_*. The DNA fragment harboring the *P_tna_* promoter was amplified from plasmid pFJ6-*P_tna_* using primers P.tna (J7)-AflII-U-F and P.tna (J7)-U-R2. *^NJ7G^cdpB3* was amplified from *Natrinema sp*.CJ7-F (*ΔpyrF*) chromosomal DNA using primers ^NJ7G^cdpB3-U-F and ^NJ7G^cdpB3-U-R. *gfp* was amplified from plasmid pIDJL40-*ftsZ1*(*P_tna_::ftsZ1-gfp bla*) using primers gfp-U-F4 and gfp-SphI-U-R. The three fragments *P_tna_, ^NJ7G^cdpB3* and *gfp* were joined by overlap extension PCR using primers P.tna (J7)-AflII-U-F and gfp-SphI-U-R to generate the cassette *P_tna_:: ^NJ7G^cdpB3-gfp.*

pZS388

The plasmid pZS388 (pZS306, *P_tna_::HAH_0460-gfp bla*) was constructed by recombination of pZS306 (digested with NdeI and ClaI) and a DNA fragment containing *^HAH^cdpB2-gfp* followed by the terminal homologous sequence of pZS306. The *^HAH^cdpB2* was amplified from *H. hispanica* DF60 (*ΔpyrF*) chromosomal DNA using primers ^HAH^cdpB2-U-F and ^HAH^cdpB2-U-R, whereas *gfp* was amplified from plasmid pIDJL40-*ftsZ1* (*P_tna_::ftsZ1-gfp bla*) using primers gfp-U-F5 and gfp-ClaI-U-R. The two DNA fragments were joined by overlap extension PCR using primers ^HAH^cdpB2-U-F and gfp-ClaI-U-R and then inserted into pZS306.

pZS389

The plasmid pZS389 (pZS306, *P_tna_::HAH_5240-gfp bla*) was constructed by recombination of pZS306 (digested with NdeI and ClaI) and a DNA fragment containing *^HAH^cdpB3-gfp* followed by the terminal homologous sequence of pZS306. The *^HAH^cdpB3* was amplified from *H. hispanica* DF60 (*ΔpyrF*) chromosomal DNA using primers ^HAH^cdpB3-U-F and ^HAH^cdpB3-U-R, whereas *gfp* was amplified from plasmid pIDJL40-*ftsZ1* (*P_tna_::ftsZ1-gfp bla*) using primers gfp-U-F6 and gfp-ClaI-U-R. The two DNA fragments were joined by overlap extension PCR using primers ^HAH^cdpB3-U-F and gfp-ClaI-U-R and then inserted into pZS306.

pZS390

The plasmid pZS390 (pZS338, *P_native_::cdpB2*-*P_native_::cdpB3 bla*) was constructed by ligation of a BamHI/NotI digested DNA fragment containing *P_native_::cdpB2* into pZS338 digested with the same enzymes. The DNA fragment was amplified from *H. volcanii* DS70 (DS2, *ΔpHV2*) chromosomal DNA using primers P_native_-cdpB2-BamHI-F and cdpB2-NotI-R.

pZS398

The plasmid pZS398 (pTA131, containing upstream and downstream flanking sequences of *cdpB2*) is a non-replicating plasmid for the generation of CdpB2 deletion strain. It was constructed by recombination of pTA131 (digested with HindIII and BamHI) and a DNA fragment harboring the upstream and downstream flanking sequences of *cdpB2* followed by the terminal homologous sequence of pTA131. The upstream and downstream flanking sequences of *cdpB1* were amplified from *H. volcanii* DS70 (DS2, *ΔpHV2*) chromosomal DNA using primer pairs cdpB2UP-Hindlll-U-F/cdpB2UP-U-R and cdpB2down-U-F/cdpB2down-BamHI-U-R, respectively. The two DNA fragments were joined by overlap extension PCR using primers cdpB2UP-Hindlll-U-F and cdpB2down-BamHI-U-R and then inserted into pTA131.

pZS399

The plasmid pZS399 (pTA131, containing upstream and downstream flanking sequences of *cdpB3*) is a non-replicating plasmid for the generation of *cdpB3* deletion strain. It was constructed by recombination of pTA131 (digested with HindIII and BamHI) and a DNA fragment harboring the upstream and downstream flanking sequences of *cdpB3* followed by the terminal homologous sequence of pTA131. The *cdpB1* upstream and downstream flanking sequences were amplified from *H. volcanii* DS70 (DS2, *ΔpHV2*) chromosomal DNA using primer pairs cdpB3UP-Hindlll-U-F/cdpB3UP-U-R and cdpB3down-U-F/cdpB3down-BamHI-U-R, respectively. The two DNA fragments were joined by overlap extension PCR using primers cdpB3UP-Hindlll-U-F and cdpB3down-BamHI-U-R and then inserted into pTA131.

pZS407

The plasmid pZS407 (pZS285, *P_phaR_::cdpB3-gfp-sepF-mCherry bla*) was constructed by recombination pZS285 (digested with NdeI and NotI) and a DNA fragment containing *cdpB3-gfp-sepF-mCherry* followed by the terminal homologous sequence of pZS285. *cdpB3-gfp* was amplified from plasmid pZS337 (*P_tna_::cdpB3-gfp*) using primers cdpB3-NdeI-U-F and gfp-U-R3, whereas *sepF-mCherry* was amplified from plasmid pZS107 (*P._tna_::cdpB1-gfp-sepF-mCherry*) using primers sepF-Notl-U-F and mCH-NotI-U-R. The two DNA fragments were joined by overlap extension PCR using primers cdpB3-NdeI-U-F and mCH-NotI-U-R and then inserted into pZS285.

pZS408

The plasmid pZS408 (pZS285, *P_phaR_::cdpB2-gfp-sepF-mCherry bla*) was constructed by recombination pZS285 (digested with NdeI and NotI) and a DNA fragment containing *cdpB2-gfp-sepF-mCherry* followed by the terminal homologous sequence of pZS285. *cdpB2-gfp* was amplified from plasmid pZS336 (*P_tna_::cdpB2-gfp*) using primers cdpB2-NdeI-U-F and gfp-U-R3, whereas *sepF-mCherry* was amplified from plasmid pZS107 (*P._tna_::cdpB1-gfp-sepF-mCherry*) using primers sepF-Notl-U-F and mCH-NotI-U-R. The two DNA fragments were joined by overlap extension PCR using primers cdpB2-NdeI-U-F and mCH-NotI-U-R and then inserted into pZS285.

pZS417

The plasmid pZS417 (pZS407, *P_phaR_::cdpB3-gfp-cdpB1-mCherry bla*) was constructed by recombination of pZS407 (digested with NotI) and a DNA fragment harboring *cdpB1-mCherry* followed by the terminal homologous sequence of pZS407. The DNA fragment was amplified from plasmid pZS101 (*P_tna_::cdpB1-mCherry bla*) using primers *cdpB1*-Notl-U-F and mCH-Notl-U-R.

pZS418

The plasmid pZS418 (pZS408, *P_phaR_::cdpB2-gfp-cdpB1-mCherry bla*) was constructed by recombination of pZS408 (digested with NotI) and a DNA fragment harboring *cdpB1-mCherry* followed by the terminal homologous sequence of pZS408. The DNA fragment was amplified from plasmid pZS101 (*P_tna_::cdpB1-mCherry bla*) using primers *cdpB1*-Notl-U-F and mCH-Notl-U-R.

pZS422

The plasmid pZS422 (pIDJL40-*ftsZ1*, *P_tna_::hvo_1607-gfp bla*) was constructed by ligation of a NdeI/BamHI digested DNA fragment containing *hvo_1607* into pIDJL40-*ftsZ1* digested with the same enzymes. The DNA fragment was amplified from *H. volcanii* DS70 (DS2, *ΔpHV2*) chromosomal DNA using primers hvo_1607-NdeI-F and hvo_1607-BamHI-R.

pZS423

The plasmid pZS423 (pIDJL40, *P_tna_::gfp-hvo_1607 bla*) was constructed by ligation of a NotI digested DNA fragment containing *hvo_1607* into pIDJL40 digested with the same enzymes. The DNA fragment was amplified from *H. volcanii* DS70 (DS2, *ΔpHV2*) chromosomal DNA using primers hvo_1607- NotI-F and hvo_1607- NotI-R.

pZWC48

The plasmid pZWC48 (pTA1228, *P_tna_*::*sfGFP10-I*-*sfGFP10-I* *bla*) was constructed by recombination of pTA1228 (digested with NdeI and NheI) and a DNA fragment harboring *sfGFP10-I*-*sfGFP10-I* followed by the terminal homologous sequence of pTA1228. *sfGFP10-I-a* and *sfGFP10-I-b* were amplified from plasmid psfGFP-10-I (pUC-19, *lacZ*::*sfGFP10-I bla*) using primer pairs GFP10-F1/GFP10-R1 and GFP10-F2/GFP10-R2, respectively. The two DNA fragments were joined by overlap extension PCR using primers GFP10-F1 and GFP10-R2 to generate the cassette *sfGFP10-I*-*sfGFP10-I* and then insert into pTA1228.

pZWC50

The plasmid pZWC50 (pTA1228, *P_tna_*::*sfGFP10-II*-*sfGFP10-II* *bla*) was constructed by recombination of pTA1228 (digested with NdeI and NheI) and a DNA fragment harboring *sfGFP10-II*-*sfGFP10-II* followed by the terminal homologous sequence of pTA1228. *sfGFP10-II-a* and *sfGFP10-II-b* were amplified from plasmid psfGFP-10-II (pUC-19, *lacZ*::*sfGFP10-II* bla) using primer pairs GFP10-F1/GFP10-R3 and GFP10-F3/GFP10-R2, respectively. The two DNA fragments were joined by overlap extension PCR using primers GFP10-F1 and GFP10-R2 to generate the cassette *sfGFP10-II*-*sfGFP10-II* and then insert into pTA1228.

pZWC51

The plasmid pZWC51 (pTA1228, *P_tna_*::*II*-*sfGFP11-II-sfGFP11* *bla*) was constructed by recombination of pTA1228 (digested with NdeI and NheI) and a DNA fragment harboring *II*-*sfGFP11-II*-*sfGFP11* followed by the terminal homologous sequence of pTA1228. *II-sfGFP11-a* and *II-sfGFP11-b* were amplified from plasmid psfGFP-11-II (pUC-19, *lacZ*::*II-sfGFP11* *bla*) using primer pairs GFP11-F1/GFP11-R1 and GFP11-F2/GFP11-R2, respectively. The two DNA fragments were joined by overlap extension PCR using primers GFP11-F1 and GFP11-R2 to generate the cassette *II*-*sfGFP11-II-sfGFP11* and then insert into pTA1228.

pZWC52

The plasmid pZWC52 (pTA1228, *P_tna_*::*I*-*sfGFP11-I-sfGFP11* *bla*) was constructed by recombination of pTA1228 (digested with NdeI and NheI) and a DNA fragment harboring *I*-*sfGFP11-I-sfGFP11* followed by the terminal homologous sequence of pTA1228. *I-sfGFP11-a* *I-sfGFP11-b* were amplified from plasmid psfGFP-11-I (pUC-19, *lacZ*::*I-sfGFP11* *bla*) using primer pairs GFP11-F1/GFP11-R3 and GFP11-F3/GFP11-R2, respectively. The two DNA fragments were joined by overlap extension PCR using primers GFP11-F1 and GFP11-R2 to generate the cassette *I*-*sfGFP11-I-sfGFP11* and then insert into pTA1228.

pZWC55

The plasmid pZWC55 (pTA1228, *P_tna_*::*sfGFP1-9-sfGFP10-II-II-sfGFP11* *bla*) was constructed by recombination of pTA1228 (digested with NdeI and NheI) and a DNA fragment harboring *sfGFP1-9-sfGFP10-II-II-sfGFP11* followed by the terminal homologous sequence of pTA1228. *sfGFP1-9* was amplified from plasmid psfGFP-1-9 (pUC-19, *lacZ*::*sfGFP1–9* *bla*) using primers sfGFP1-F and sfGFP9-R1, *sfGFP10-II* was amplified from plasmid psfGFP-10-II (pUC-19, *lacZ*::*sfGFP10-II* *bla*) using primers sfGFP10S-F and sfGFP10S-R1, and the cassette *II-sfGFP11* was amplified from plasmid pZWC51 (pTA1228, *P_tna_*::*sfGFP11-II*-*sfGFP11-II* *bla*) using primers sfGFPS11-F1 and sfGFPS11-R. *sfGFP1-9, sfGFP10-II* and *II-sfGFP11* were joined by overlap extension PCR using primers sfGFP10S-F and sfGFPS11-R to generate the cassette *sfGFP1-9*-*sfGFP10-II-II-sfGFP11.*

pZWC56

The plasmid pZWC56 (pTA1228, *P_tna_*::*sfGFP1-9-sfGFP10-II-sfGFP11-II bla*) was constructed by recombination of pTA1228 (digested with NdeI and NheI) and a DNA fragment harboring *sfGFP1-9-sfGFP10-II-sfGFP11-II* followed by the terminal homologous sequence of pTA1228. *sfGFP1-9* was amplified from plasmid psfGFP-1-9 (pUC-19, *lacZ*::*sfGFP1–9* *bla*) using primers sfGFP1-F and sfGFP9-R1, *sfGFP10-II* was amplified from plasmid psfGFP-10-II (pUC-19, *lacZ*::*sfGFP10-II* *bla*) using primers sfGFP10S-F and sfGFP10S-R2, and *sfGFP11-II* was amplified from plasmid pZWC51 (pTA1228, *P_tna_*::*II*-*sfGFP11-II-sfGFP11* *bla*) using primers sfGFP11S-F and sfGFP11S-R. The cassette *sfGFP10-II-sfGFP11-II* was amplified by overlap extension PCR using *sfGFP10-II* and *sfGFP11-II* as templates and primers sfGFP10S-F and sfGFP11S-R. *sfGFP1-9* and *sfGFP10-II-sfGFP11-II* were joined by overlap extension PCR to generate *sfGFP1-9*-*sfGFP10-II-sfGFP11-II* using primers sfGFP1-F and sfGFP11S-R, and the PCR product was cut and inserted into pTA1228.

pZWC57

The plasmid pZWC57 (pTA1228, *P_tna_*::*sfGFP1-9-II-sfGFP10-II-sfGFP11 bla*) was constructed by recombination of pTA1228 (digested with NdeI and NheI) and a DNA fragment *sfGFP1-9-II-sfGFP10-II-sfGFP11* followed by the terminal homologous sequence of pTA1228.*sfGFP1-9* was amplified from plasmid psfGFP-1-9 (pUC-19, *lacZ*::*sfGFP1–9* *bla*) using primers sfGFP1-F and sfGFP9-R2, whereas *II-sfGFP10* was amplified from plasmid pZWC50 (pTA1228, *P_tna_*::*sfGFP10-II*-*sfGFP10-II* *bla*) using primers sfGFPS10-F1 and sfGFPS10-R1. *II-sfGFP11* was amplified from plasmid pZWC51 (pTA1228, *P_tna_*::*sfGFP11-II*-*sfGFP11-II* *bla*) using primers sfGFPS11-F2 and sfGFPS11-R. *II-sfGFP10* and *II-sfGFP11* were joined by overlap extension PCR to generate *II-sfGFP10-II-sfGFP11* using primers sfGFPS10-F1 and sfGFPS11-R. *sfGFP1-9* and *II-sfGFP10-II-sfGFP11* were joined by overlap extension PCR to generate *sfGFP1-9*-*II-sfGFP10-II-sfGFP11* using primers sfGFP1-F and sfGFPS11-R. The PCR product was digested and ligated into pTA1228.

pZWC61

The plasmid pZWC61 (pTA1228, *P_tna_*::*sfGFP1-9-sfGFP10-II-I-sfGFP11 bla*) was constructed by recombination of pTA1228 (digested with NdeI and NheI) and a DNA fragment harboring *sfGFP1-9-sfGFP10-II-I-sfGFP11* followed by the terminal homologous sequence of pTA1228. *sfGFP1-9* was amplified from plasmid psfGFP-1-9 (pUC-19, *lacZ*::*sfGFP1-9* *bla*) using primers sfGFP1-F and sfGFP9-R1, whereas *sfGFP10-II* was amplified from plasmid psfGFP-10-II (pUC-19, *lacZ*::*sfGFP10-II* *bla*) using primers sfGFP10S-F and sfGFP10S-R3, and *I-sfGFP11* was amplified from plasmid pZWC52 (pTA1228, *P.tna*::*sfGFP11-I*-*sfGFP11-I* *bla*) using primers sfGFPL11-F and sfGFP11S-R. *sfGFP10-II* and *I-sfGFP11* were joined by by overlap extension PCR using primers sfGFP10S-F and sfGFPS11-R to generate *sfGFP10-II-I-sfGFP1*1, which was further joined with *sfPGFP1-9* by overlap extension PCR using primers sfGFP1-F and sfGFPS11-R.

pZWC62

The plasmid pZWC62 (pTA1228, *P_tna_*::*sfGFP1-9-sfGFP10-II-sfGFP11-I bla*) was constructed by recombination of pTA1228 (digested with NdeI and NheI) and a DNA fragment containing *sfGFP1-9-sfGFP10-II-sfGFP11-I* followed by the terminal homologous sequence of pTA1228. *sfGFP1-9* was amplified from plasmid psfGFP-1-9 (pUC-19, *lacZ*::*sfGFP1–9* *bla*) using primers sfGFP1-F and sfGFP9-R1, *sfGFP10-II* was amplified from plasmid psfGFP-10-II (pUC-19, *lacZ*::*sfGFP10-II* *bla*) using primers sfGFP10S-F and sfGFP10S-R2, and *sfGFP11-I* was amplified from plasmid pZWC52 (pTA1228, *P_tna_*::*sfGFP11-I*-*sfGFP11-I* *bla*) using primers sfGFP11S-F and sfGFP11L-R. *sfGFP10-II* and *sfGFP11-I* were joined by overlap extension PCR using primers sfGFP10S-F and sfGFP11L-R to generate *sfGFP10-II-sfGFP11-I*, which was further joined with sfGFP1-9 by overlap extension PCR to generate *sfGFP1-9-sfGFP10-II-sfGFP11-I*. The final PCR product was then inserted into pTA1228 to generate pZWC62

pZWC63

The plasmid pZWC63 (pTA1228, *P_tna_*::*sfGFP1-9-II-sfGFP10-I-sfGFP11 bla*) was constructed by recombination of pTA1228 (digested with NdeI and NheI) and a DNA fragment harboring *sfGFP1-9-II-sfGFP10-I-sfGFP11* followed by the terminal homologous sequence of pTA1228. *sfGFP1-9* was amplified from plasmid psfGFP-1-9 (pUC-19, *lacZ*::*sfGFP1–9* *bla*) using primers sfGFP1-F and sfGFP9-R2, *II-sfGFP10* was amplified from plasmid pZWC50 (pTA1228, *P_tna_*::*sfGFP10-II*-*sfGFP10-II* *bla*) using primers sfGFPS10-F1 and sfGFPS10-R2, and *I-sfGFP11* was amplified from plasmid pZWC52 (pTA1228, *P_tna_*::*sfGFP11-I*-*sfGFP11-I* *bla*) using primers sfGFPL11-F2 and sfGFPS11-R. *II-sfGFP10* and *I-sfGFP11* were joined by overlap extension PCR using primers sfGFPS10-F1 and sfGFPS11-R to generate *II-sfGFP10-I-sfGFP11*, which was further joined with *sfGFP1-9* to generate the cassette *sfGFP1-9-II-sfGFP10-I-sfGFP11* by overlap extension PCR using primers sfGFP1-F and sfGFPS11-R. The final PCR product was inserted into pTA1228 to generate pZWC63.

pZWC64

The plasmid pZWC64 (pTA1228, *P_tna_*::*sfGFP1-9-sfGFP10-I-II-sfGFP11 bla*) was constructed by recombination of pTA1228 (digested with NdeI and NheI) and a DNA fragment harboring *sfGFP1-9-sfGFP10-I-II-sfGFP11* followed by the terminal homologous sequence of pTA1228. *sfGFP1-9* was amplified from plasmid psfGFP-1-9 (pUC-19, *lacZ*::*sfGFP1–9* *bla*) using primers sfGFP1-F and sfGFP9-R1, *sfGFP10-I* was amplified from plasmid psfGFP-10-I (pUC-19, *lacZ*::*sfGFP10-I* *bla*) using primers sfGFP10S-F and sfGFP10L-R, and *II-sfGFP11* was amplified from plasmid pZWC51 (pTA1228, *P_tna_*::*sfGFP11-II*-*sfGFP11-II* *bla*) using primers sfGFPS11-F3 and sfGFPS11-R. *sfGFP10-I* and *II-sfGFP11* were joined by overlap extension PCR to generate *sfGFP10-I-II-sfGFP11*, which was further joined with *sfGFP1-9* to generate the cassette *sfGFP1-9-sfGFP10-I-II-sfGFP11* by overlap extension PCR using primers sfGFP1-F and sfGFPS11-R. The final PCR product was inserted into pTA1228 to generate pZWC64.

pZWC78

The plasmid pZWC78 (pTA1228, *P_tna_*::*sfGFP1-9-sfGFP10-cdpB1-sfGFP11-I bla*) was constructed by recombination of pZWC62 (digested with HindIII) and a DNA fragment containing *cdpB1* followed by the terminal homologous sequence of pZWC62. The DNA fragment *cdpB1* was amplified from *H. volcanii* DS70 (DS2, *ΔpHV2*) chromosomal DNA using primers cdpB1-F1 and cdpB1-R1.

pZWC91

The plasmid pZWC91 (pTA1228, *P_tna_*::*sfGFP1-9-sfGFP10-ftsZ1-cdpB1-sfGFP11 bla*) was constructed by recombination of pZWC64 (digested with HindIII and XbaI) and a DNA harboring fragment *ftsZ1-cdpB1* followed by the terminal homologous sequence of pZWC64. *ftsZ1* and *cdpB1* were amplified from *H. volcanii* DS70 (DS2, *ΔpHV2*) chromosomal DNA using primer pairs ftsZ1-F1 /ftsZ1-R1 and cdpB1-F2/cdpB1-R2, respectively. *ftsZ1* and *cdpB1* were joined by overlap extension PCR using primers ftsZ1-F1 and cdpB1-R2 and then inserted into pZWC64.

pZWC92

The plasmid pZWC92 (pTA1228, *P_tna_*::*sfGFP1-9-sfGFP10-cdpB1-sfGFP11-ftsZ1 bla*) was constructed by two steps of recombination. To generate the plasmid, *cdpB1 and ftsZ1* were first amplified from *H. volcanii* DS70 (DS2, *ΔpHV2*) chromosomal DNA using primer pairs cdpB1-F3/cdpB1-R3 and ftsZ1-F2/ftsZ1-R2, respectively. The plasmid was then constructed by recombination of pZWC62 (digested with HindIII) and a DNA fragment containing *cdpB1* followed by the terminal homologous sequence of pZWC62. The intermediate plasmid was digested with XbaI and then recombined with a DNA fragment containing *ftsZ1* followed by the terminal homologous sequence of pZWC62.

pZWC93

The plasmid pZWC93 (pTA1228, *P_tna_*::*sfGFP1-9-sfGFP10-cdpB1-ftsZ1-sfGFP11 bla*) was constructed by recombination of pZWC61 (digested with HindIII and XbaI) and a DNA fragment containing *cdpB1-ftsZ1* followed by the terminal homologous sequence of pZWC61. *cdpB1 and ftsZ1* were amplified from *H. volcanii* DS70 (DS2, *ΔpHV2*) chromosomal DNA using primer pairs cdpB1-F4/cdpB1-R4 and ftsZ1-F3/ ftsZ1-R3, respectively. *cdpB1* and *ftsZ1* were joined together by overlap extension PCR to generate the cassette *cdpB1*- *ftsZ1* using primers cdpB1-F4 and ftsZ1-R3.

pZWC94

The plasmid pZWC94 (pTA1228, *P_tna_*::*sfGFP1-9-cdpB1-sfGFP10-ftsZ1-sfGFP11 bla*) was constructed by two steps of recombination. *cdpB1 and ftsZ1* were amplified from *H. volcanii* DS70 (DS2, *ΔpHV2*) chromosomal DNA using primer pairs cdpB1-F5/cdpB1-R5 and ftsZ1-F4/ ftsZ1-R4, respectively. pZWC63 (digested with HindIII) was recombined with the *cdpB1* cassette containing the terminal homologous sequence of pZWC63 to generate an intermediate plasmid, which was then cut with XbaI and recombined with *ftsZ1* digested with XbaI.

pZWC95

The plasmid pZWC95 (pTA1228, *P_tna_*::*sfGFP1-9-sfGFP10-ftsZ2-cdpB1-sfGFP11 bla*) was constructed by recombination of pZWC64 (digested with HindIII and XbaI) and a DNA fragment containing *ftsZ2-cdpB1* and the terminal homologous sequence of pZWC64. *ftsZ2* and cdpB1 were amplified from *H. volcanii* DS70 (DS2, *ΔpHV2*) chromosomal DNA using primer pairs ftsZ2-F1/ftsZ2-R1 and cdpB1-F6/cdpB1-R6, respectively. The two DNA fragments were joined together by overlap extension PCR to generate the cassette *ftsZ2-cdpB1* using primers ftsZ2-F1 and cdpB1-R6.

pZWC96

The plasmid pZWC96 (pTA1228, *P_tna_*::*sfGFP1-9-sfGFP10-cdpB1-sfGFP11-ftsZ2 bla*) was constructed by two steps of recombination. *cdpB1* and *ftsZ2* were amplified from *H. volcanii* DS70 (DS2, *ΔpHV2*) chromosoml DNA using primer pairs cdpB1-F7/cdpB1-R7 and ftsZ2-F2/ftsZ2-R2, respectively. pZWC62 (digested with HindIII) was recombined with the DNA fragment containing *cdpB1* and the terminal homologous sequence of pZWC62 to produce an intermediate plasmid, which was digested with XbaI and then recombined with the *ftsZ2* cassette.

pZWC97

The plasmid pZWC97 (pTA1228, *P_tna_*::*sfGFP1-9-sfGFP10-cdpB1-ftsZ2-sfGFP11 bla*) was constructed by recombination of pZWC61 (digested with HindIII and XbaI) and a DNA fragment containing *cdpB1-ftsZ2* and the terminal homologous sequence of pZWC61. *cdpB1* and *ftsZ* were amplified from *H. volcanii* DS70 (DS2, *ΔpHV2*) chromosomal DNA using primer pairs cdpB1-F8/cdpB1-R8 and ftsZ2-F3/ftsZ2-R3, respectively. The two DNA fragments were joined together by overlap extension PCR to generate the cassette *cdpB1-ftsZ2* using primers cdpB1-F8 and ftsZ2-R3.

pZWC98

The plasmid pZWC98 (pTA1228, *P_tna_*::*sfGFP1-9-cdpB1-sfGFP10-ftsZ2-sfGFP11 bla*) was constructed by two steps of recombination. *cdpB1* and *ftsZ2* were amplified from *H. volcanii* DS70 (DS2, *ΔpHV2*) chromosomal DNA using primer pairs cdpB1-F9/dpB1-R9 and ftsZ2-F4/ftsZ2-R4, respectively. pZWC63 (digested with HindIII) was recombined with the *cdpB1* cassette containing the terminal homologous sequence of pZWC63 to generate an intermediate plasmid, which was then digested with XbaI and recombined with the *ftsZ2 cassette.*

pZWC99

The plasmid pZWC99 (pTA1228, *P_tna_*::*sfGFP1-9- sfGFP10-sepF-cdpB1-sfGFP11 bla*) was constructed by recombination of pZWC55 (digested with HindIII and XbaI) and a DNA fragment containing the cassette *sepF-cdpB1* and the terminal homologous sequence of pZWC55. *sepF* and *cdpB1* were amplified from *H. volcanii* DS70 (DS2, *ΔpHV2*) chromosomal DNA using primer pairs sepF-F1/sepF-R1 and cdpB1-F10/cdpB1-R10, respectively. The two DNA fragments were joined by overlap extension PCR using primers sepF-F1 and cdpB1-R10.

pZWC100

The plasmid pZWC100 (pTA1228, *P_tna_*::*sfGFP1-9-sfGFP10-sepF-sfGFP11-cdpB1 bla*) was constructed by two steps of recombination. *sepF* and *cdpB1* were amplified from *H. volcanii* DS70 (DS2, *ΔpHV2*) chromosomal DNA using primer pairs sepF-F2/sepF-R2 and cdpB1-F11/cdpB1-R11, respectively. pZWC56 (digested with HindIII) was recombined with the DNA fragment containing *sepF* and the terminal homologous sequence of pZWC56 to generate an intermediate plasmid, which was digested with XbaI and then recombined with *cdpB1* to produce pZWC100.

pZWC101

The plasmid pZWC101 (pTA1228, *P_tna_*::*sfGFP1-9-sfGFP10-cdpB1-sepF-sfGFP11 bla*) was constructed by recombination of pZWC55 (digested with HindIII and XbaI) and a DNA fragment containing *cdpB1-sepF* and the terminal homologous sequence of pZWC55. *cdpB1* and *sepF* were amplified from *H. volcanii* DS70 (DS2, *ΔpHV2*) chromosomal DNA using primer pairs cdpB1-F12/cdpB1-R12 and sepF-F3/sepF-R3, respectively. The two DNA fragments were joined by overlap extension PCR using primers cdpB1-F12 and sepF-R3, the PCR product was cut and ligated into pZWC55.

pZWC102

The plasmid pZWC102 (pTA1228, *P_tna_*::*sfGFP1-9-sepF-sfGFP10-cdpB1-sfGFP11 bla*) was constructed by two steps of recombination. *sepF* and *cdpB1* were amplified from *H. volcanii* DS70 (DS2, *ΔpHV2*) chromosomal DNA using primer pairs sepF-F4/sepF-R4 and cdpB1-F13/cdpB1-R13, respectively. pZWC57 (digested with HindIII) was recombined with the *sepF* cassette containing the terminal homologous sequence of pZWC57 to generate an intermediate plasmid, which was digested with XbaI and then recombined with *cdpB1* to generate pZWC102.

pZWC144

The plasmid pZWC144 (pTA1228, *P_tna_*::*sfGFP1-9-sfGFP10-II-sfGFP11-ftsZ2 bla* ) was constructed by recombination of pZWC62 (digested with XbaI) and a DNA fragment containing *ftsZ2* and the terminal homologous sequence of pZWC62. The DNA fragment *ftsZ2* was amplified from *H. volcanii* DS70 (DS2, *ΔpHV2*) chromosomal DNA using primers ftsZ2-F5 and ftsZ2*-*R5.

pZWC145

The plasmid pZWC145 (pTA1228, pTA1228, *P_tna_*::*sfGFP1-9-cdpB1-sfGFP10-I-sfGFP11 bla*) was constructed by recombination of pZWC63 (digested with HindIII) and a DNA fragment containing *cdpB1* and the terminal homologous sequence of pZWC63. The DNA fragment containg *cdpB1* was amplified from *H. volcanii* DS70 (DS2, ΔpHV2) chromosome DNA using primers cdpB1-F15 and cdpB1-R15.

pZWC146

The plasmid pZWC146 (pTA1228, *P_tna_*::*sfGFP1-9-II-sfGFP10-ftsZ2-sfGFP11 bla* ) was constructed by recombination of pZWC63 (digested with XbaI) and a DNA fragment containing *ftsZ2* and the terminal homologous sequence of pZWC63. The DNA fragment containing *ftsZ2* was amplified from *H. volcanii* DS70 (DS2, *ΔpHV2*) chromosomal DNA using primers ftsZ2-F6 and ftsZ2-R6.

pZWC147

The plasmid pZWC147 (pTA1228, *P_tna_*::*sfGFP1-9-sfGFP10-cdpB1-II-sfGFP11 bla* ) was constructed by recombination of pZWC55 (digested with HindIII) and a DNA fragment containing *cdpB1* and the terminal homologous sequence of pZWC55. The DNA fragment containing *cdpB1* was amplified from *H. volcanii* DS70 (DS2, *ΔpHV2*) chromosomal DNA using primers cdpB1-F16 and cdpB1-R16.

pZWC148

The plasmid pZWC148 (pTA1228, *P_tna_*::*sfGFP1-9-sfGFP10-II-sepF-sfGFP11 bla* ) was constructed by recombination of pZWC55 (digested with XbaI) and a DNA fragment containing *sepF* and the terminal homologous sequence of pZWC55. The DNA fragment containing *sepF* was amplified from *H. volcanii* DS70 (DS2, *ΔpHV2*) chromosomal DNA using primers sepF-F5 and sepF-R5.

pZWC149

The plasmid pZWC149 (pTA1228, *P_tna_*::*sfGFP1-9-II-sfGFP10-cdpB1-sfGFP11 bla* ) was constructed by recombination of pZWC57 (digested with XbaI) and a DNA fragment containing *cdpB1* and the terminal homologous sequence of pZWC55. The DNA fragment containing *cdpB1* was amplified from *H. volcanii* DS70 (DS2, *ΔpHV2*) chromosomal DNA using primers cdpB1-F17 and cdpB1-R17.

pZWC150

The plasmid pZWC150 (pTA1228, *P_tna_*::*sfGFP1-9-sepF-sfGFP10-II-sfGFP11 bla* ) was constructed by recombination of pZWC57 (digested with HindIII) and a DNA fragment containing *sepF* and the terminal homologous sequence of pZWC57. The DNA fragment containing *sepF* was amplified from *H. volcanii* DS70 (DS2, *ΔpHV2*) chromosomal DNA using primers sepF-F6 and sepF-R6.

pZWC165

The plasmid pZWC165 (pTA1228, *P_tna_*::*sfGFP1-9-sfGFP10-II-sfGFP11-I bla* ) was constructed by recombination of pZWC62 (digested with EcoRI and HindIII) and a DNA fragment containing *sfGFP10-II* and the terminal homologous sequence of pZWC62. The DNA fragment *sfGFP10-II* was amplified from plasmid psfGFP-10-II (pUC-19, *lacZ*::*sfGFP10-II* *bla*) using primers sfGFP10S-F and sfGFP10S-R4.

pZWC166

The plasmid pZWC166 (pTA1228, *P_tna_*::*sfGFP1-9-sfGFP10-II-I-sfGFP11 bla* ) was constructed by recombination of pZWC61 (digested with EcoRI and XbaI) and a DNA fragment containing *sfGFP10*-*II* and the terminal homologous sequence of pZWC61. The DNA fragment containing *sfGFP10*-*II* was amplified from plasmid pZWC50 (pTA1228, *P_tna_*::*sfGFP10-II*-*sfGFP10-II* bla) using primers sfGFP10S-F and sfGFP10S-R7

pZWC167

The plasmid pZWC167 (pTA1228, *P_tna_*::*sfGFP1-9-II-sfGFP10-I-sfGFP11 bla* ) was constructed by recombination of pZWC63 (digested with EcoRI and XbaI) and a DNA fragment containing *II-sfGFP10* and the terminal homologous sequence of pZWC63. The DNA fragment containing *II-sfGFP10* was amplified from plasmid pZWC50 (pTA1228, *P_tna_*::*sfGFP10-II*-*sfGFP10-II* bla) using primers sfGFPS10-F2 and sfGFPS10-R3.

pZWC168

The plasmid pZWC168 (pTA1228, *P_tna_*::*sfGFP1-9-sfGFP10-II-II-sfGFP11 bla*) was constructed by recombination of pZWC61 (digested with EcoRI and HindIII) and a DNA fragment containing *sfGFP10-II* and the terminal homologous sequence of pZWC61. The DNA fragment containing *sfGFP10-II* was amplified from plasmid psfGFP-10-II (pUC-19, *lacZ*::*sfGFP10-II* *bla*) using primers sfGFP10S-F and sfGFP10S-R6.

pZWC169

The plasmid pZWC169 (pTA1228, *P_tna_*::*sfGFP1-9-II-sfGFP10-II-sfGFP11 bla*) was constructed by recombination of pZWC57 (digested with EcoRI and XbaI) and a DNA fragment containing *II-sfGFP10* and the terminal homologous sequence of pZWC57. The DNA fragment containing *II-sfGFP10* was amplified from plasmid pZWC50 (pTA1228, *P_tna_*::sfGFP1-9-II-*sfGFP10-II*-*sfGFP10* bla) using primers sfGFPS10-F2 and sfGFPS10-R4.

pZWC176

The plasmid pZWC176 (pIDJL40-*ftsZ1*, *P_native_*::*sepF-flag bla*) was constructed by recombination of pIDJL40-*ftsZ1* (digested with ApaI and NotI) and a DNA fragment containg *P_native_*-*sepF-flag* and the terminal homologous sequence of pIDJL40-*ftsZ1*. *P_native_* was amplified from *H. volcanii* DS70 (DS2, ΔpHV2) chromosomal DNA using primers P_native_-sepF-F and P_native_-sepF-R. *sepF-flag* was amplified from a DNA fragment harboring *sepF* using primers sepF-F7 and sepF-flag-R2. *sepF* was amplified from plasmid pZWC102 (pTA1228, *P_tna_*::*sfGFP1-9-sepF-sfGFP10-cdpB1-sfGFP11* bla) using primers sepF-F7 and sepF-flag-R1. P_native_ and *sepF-flag* were joined by overlap extension PCR of using primers P_native_-sepF-F and sepF-flag-R2.

pZWC185

The plasmid pZWC185 (pZWC176, *P_native_*::*sepF-flag P_native_*::*cdpB1-gfp bla*) was constructed by recombination of pZWC176 (digested with NotI) and a DNA fragment containing *P_native_::cdpB1-gfp* and the terminal homologous sequence of pZWC176. The DNA fragment containing *P_native_::cdpB1-gfp* was amplified from plasmid pZS214 (pIDJL40-*ftsZ1*, *P_native_::cdpB1-gfp bla*) using primers P_native_-cdpB1-F and gfp-R.

**References**

1.	Pichoff S, Vollrath B, Touriol C, Bouche JP. Deletion analysis of gene minE which encodes the topological specificity factor of cell division in Escherichia coli. Mol Microbiol. 1995;18(2):321-9.

2.	Miroux B, Walker JE. Over-production of proteins in Escherichia coli: mutant hosts that allow synthesis of some membrane proteins and globular proteins at high levels. J Mol Biol. 1996;260(3):289-98.

3.	Wendoloski D, Ferrer C, Dyall-Smith ML. A new simvastatin (mevinolin)-resistance marker from Haloarcula hispanica and a new Haloferax volcanii strain cured of plasmid pHV2. Microbiology (Reading). 2001;147(Pt 4):959-64.

4.	Allers T, Ngo HP, Mevarech M, Lloyd RG. Development of additional selectable markers for the halophilic archaeon Haloferax volcanii based on the leuB and trpA genes. Appl Environ Microbiol. 2004;70(2):943-53.

5.	Liao Y, Ithurbide S, Evenhuis C, Lowe J, Duggin IG. Cell division in the archaeon Haloferax volcanii relies on two FtsZ proteins with distinct functions in division ring assembly and constriction. Nat Microbiol. 2021;6(5):594-605.

6.	Liu H, Han J, Liu X, Zhou J, Xiang H. Development of pyrF-based gene knockout systems for genome-wide manipulation of the archaea Haloferax mediterranei and Haloarcula hispanica. J Genet Genomics. 2011;38(6):261-9.

7.	Wang J, Liu Y, Liu Y, Du K, Xu S, Wang Y, et al. A novel family of tyrosine integrases encoded by the temperate pleolipovirus SNJ2. Nucleic Acids Res. 2018;46(5):2521-36.

8.	Allers T, Barak S, Liddell S, Wardell K, Mevarech M. Improved strains and plasmid vectors for conditional overexpression of His-tagged proteins in Haloferax volcanii. Appl Environ Microbiol. 2010;76(6):1759-69.

9.	Duggin IG, Aylett CH, Walsh JC, Michie KA, Wang Q, Turnbull L, et al. CetZ tubulin-like proteins control archaeal cell shape. Nature. 2015;519(7543):362-5.

10.	Cai S, Cai L, Liu H, Liu X, Han J, Zhou J, et al. Identification of the haloarchaeal phasin (PhaP) that functions in polyhydroxyalkanoate accumulation and granule formation in Haloferax mediterranei. Appl Environ Microbiol. 2012;78(6):1946-52.

11.	Chen B, Chen Z, Wang Y, Gong H, Sima L, Wang J, et al. ORF4 of the Temperate Archaeal Virus SNJ1 Governs the Lysis-Lysogeny Switch and Superinfection Immunity. J Virol. 2020;94(16).

12.	Cabantous S, Nguyen HB, Pedelacq JD, Koraichi F, Chaudhary A, Ganguly K, et al. A new protein-protein interaction sensor based on tripartite split-GFP association. Sci Rep. 2013;3:2854.
